# Desert Cucurbit Microbiomes: Spatiotemporal Dynamics and Functional Adaptations

**DOI:** 10.64898/2026.05.07.723578

**Authors:** Miranda Procter, Biduth Kundu, Naganeeswaran Sudalaimuthuasari, Raja S. AlMaskari, Iltaf Shah, Shaima Alnuaimi, Farah Husain, Khawla Aldhaheri, Khaled M. Hazzouri, Khaled M. A. Amiri

## Abstract

**Background:** Plant microbiomes can contribute to host adaptation in extreme environments, particularly deserts where high temperatures, intense radiation, water limitation, and nutrient-poor soils constrain plant survival. *Citrullus colocynthis* is a desert-adapted cucurbit with medicinal and agricultural relevance, yet integrated understanding of its microbiome across seasons, tissues, habitats, and functional traits remains limited. Here, we asked whether spatiotemporal microbiome dynamics in *C. colocynthis* are linked to microbial functional potential and cultivable traits relevant to persistence under arid conditions.

**Results:** To address this, we profiled the microbiome of *C. colocynthis* across two sites, two seasons, and multiple compartments using 16S rRNA amplicon screening, shotgun metagenomics, culture-based phenotyping, and genome analysis. Amplicon profiling provided an exploratory framework and showed that plant-associated bacterial communities were shaped primarily by season and tissue type, with roots emerging as the most season-responsive compartment. Shotgun metagenomics confirmed stronger seasonal restructuring in root bacterial communities than in leaves and extended taxonomic profiling to fungal and archaeal fractions, which were interpreted descriptively because of low read representation. Summer root communities were enriched in actinobacterial genera, including *Saccharopolyspora*, *Amycolatopsis*, *Pseudonocardia*, and *Nonomuraea*, while functional profiling indicated coordinated shifts in central metabolism, cofactor salvage, exopolysaccharide-related pathways, and redox-associated functions. Twenty-four cultured bacterial isolates exhibited diverse stress-tolerance and plant growth-promoting traits, and whole-genome analyses identified biosynthetic, osmoprotective, oxidative-stress, and phytohormone-related gene content. Pangenome analysis of *Pseudomonas orientalis* revealed an open pangenome and lifestyle-associated accessory genes linked to host-associated functions.

**Conclusions:** Together, these findings connect seasonal bacterial community turnover, metagenomic functional potential, and cultivable microbial traits in a desert plant holobiont, highlighting desert-native bacteria and pathways relevant to microbiome-guided strategies for arid agriculture.

## 1. Introduction

Increasing heat, drought, and salinity are major constraints on global agriculture and make crop resilience a growing priority [1, 2] [3, 4]. In desert systems, plant persistence is not explained by plant physiology alone; microbial partners can contribute to tolerance under water limitation, high temperature, and poor nutrient availability [5–9]. However, the extent to which these associations vary across plant habitats and seasons remains somewhat unresolved. Several studies show that plant-associated microbial communities change between wet and dry periods, with leaves and roots often responding more strongly than surrounding soils [10–14]. These temporal and spatial patterns are important for linking community change to possible functions.

*Citrullus colocynthis* provides a useful system for studying these processes under natural desert conditions. This cucurbit has both medicinal and agricultural relevance [15–19]and shows physiological adaptation to extreme temperatures [20]. Previous studies have reported disease-suppressive and growth-promoting endophytes and rhizobacteria [21], antibacterial endophytic actinobacteria [22], tissue-and site-dependent fungal endophytes [23], and limited fungal diversity in wider desert surveys that included this species [24]. Our earlier bacterial microbiome study also showed diverse plant-associated bacteria, many with known plant growth-promoting potential [4], and later culture-based work showed that rhizosphere isolates can carry multiple arid-adaptation traits [25]. These findings support examining the *C. colocynthis* holobiont in a way that connects community dynamics with functional traits.

The holobiont concept treats the plant and its associated microbes as an interacting system shaped by both host and microbial processes [26, 27]. Within this view, plant-associated and free-living microbes may differ in traits relevant to host interaction. Plant-associated *Pseudomonas*, for example, often contain genes linked to rhizosphere competence, including antibiotic and siderophore pathways, as well as functions involved in induced systemic resistance. Many beneficial bacteria also produce indole-3-acetic acid (IAA), commonly via tryptophan-dependent pathways, which affect plant development and defense [28–31]. These observations provide a basis for asking whether seasonal community changes, habitat specificity in leaves, roots, and soils, and microbial functions associated with stress tolerance are connected.

Here, we examined whether spatiotemporal microbiome dynamics and microbial traits reflect the ecological demands of a desert plant. We profiled *C. colocynthis* tissues and surrounding soils from two UAE emirates during rainy and dry seasons using 16S rRNA amplicon sequencing and shotgun metagenomics. We also isolated endophytic strains from leaves and roots, sequenced their genomes, and evaluated stress tolerance and plant growth-promoting traits, including in vitro phytohormone production and antimicrobial activity. By combining this data with previous work, we aim to clarify how the *C. colocynthis* holobiont may contribute to resilience in arid environments.

## 2. Methods

### 2.1. Sample collection

Leaf, root, root zone, and bulk soil samples were collected from three *C. colocynthis* plants in natural populations in Al Ain city (24°27’50.8“ N 55°38’37.1” E) and in the Ras Al Khaimah (RAK) emirate (25°33’11.7“ N 55°53’18.9” E), UAE, as described previously [4]. The study included three biological replicates, each with three technical replicates per sample type and season. The locations were selected for their distinct rainfall patterns and geography. Endophytic bacteria were isolated from freshly collected leaf and root samples from the Al Ain site in January 2022, and samples collected in March and August 2022 were used for microbiome analysis.

### 2.2. Amplicon Profiling and Metagenomic Analyses

#### 2.2.1. DNA extraction

Root and leaf samples were surface sterilized (1% bleach, 3 min; two sterile water washes; 70% ethanol, 3 min; six sterile water washes) [3]. The final wash water was plated on nutrient agar (NA, HiMedia, Mumbai, India) and potato dextrose agar (PDA, Conda Lab, Madrid, Spain) to assess sterility. DNA was extracted using the XpressDNA Plant Kit (MagGenome, India) following the manufacturer’s protocol. However, root samples were incubated at 56 °C for an additional 30 minutes during the lysis step. DNA was extracted from 2g of soil using a combination of the DNeasy PowerSoil Pro (Qiagen, Hilden, Germany) and XpressDNA Soil Kit (MagGenome). These extractions were repeated and pooled within the same biological replicate (i.e., technical replicates were pooled) until sufficient DNA was obtained for the respective sequencing strategies. The quality and quantity of the extracted DNA were confirmed as previously described [4].

#### 2.2.2. 16S rRNA Amplicon-based Bacterial Profiling

16S rRNA amplicon sequencing was used to assess DNA suitability for downstream metagenomics and to provide preliminary insights into seasonal and geographic variation in bacterial community structure. DNA samples were amplified, trimmed, and quality assessed as described previously [4]. The generated ASV files were used for analyses with MicrobiomeAnalyst2 [32]and the updated Greengenes database [33]. Alpha diversity was assessed using Shannon [34] and Chao1 [35] indices, while NMDS was used to examine beta diversity, which guided subsequent metagenomic analysis.

#### 2.2.3. Metagenome-based Microbial Profiling

Shotgun metagenomic sequencing was performed on an Illumina NovaSeq PE150 (150 bp paired-end) platform, generating approximately 20 GB of raw data per sample. The quality of the PE reads was assessed with FastQC v.0.11.9 [36]. Low-quality reads and adapter sequences were removed using Trimmomatic v.0.39 [37]. The trimmed reads were taxonomically profiled using Kraken 2 [38].

#### 2.2.4. Seasonal and Geographic Comparisons of Metagenome-derived Microbial Communities

Kraken-based reports were combined with metadata to compare leaf and root microbial communities across seasons and locations. Analyses were restricted to leaf and root profiles, with viral assignments and other sample classes excluded. Bacterial, fungal, and archaeal assignments were retained, but low representation of fungal and archaeal reads limited these groups to descriptive phylum-and order-level summaries. Kraken lineage-style reports were parsed to retain terminal assignments, collapse profiles to comparable taxonomic levels, and generate relative-abundance tables for downstream analyses.

For bacterial genus-level comparisons, alpha diversity was summarized using observed genera, Shannon diversity, and Pielou evenness [39]. Compositional turnover was assessed using Bray-Curtis dissimilarities [40], metric multidimensional scaling, and PERMANOVA [41] within each location-tissue subgroup. Seasonal changes in alpha diversity and genus-level relative abundance were tested using exact two-sided Mann-Whitney tests, Cliff’s delta effect sizes, and false discovery rate correction within each test family. Interpretation emphasized effect sizes, multivariate patterns, and directional consistency.

#### 2.2.5. Functional Profiling using Root Samples

Functional profiling was restricted to root samples because this compartment exhibited the strongest seasonal restructuring and serves as the main interface for plant–microbe interactions under abiotic stress. Analyses used HUMAnN gene-family, pathway-abundance, pathway-coverage, and merged relative-abundance outputs [42, 43]. The merged pathway-abundance matrix was used for downstream analyses. Unstratified pathway outputs were used for community-level ordination and seasonal comparisons, while stratified outputs identified contributing taxa. Pathway names with provenance tags, including plant, yeast, and archaeal labels, were retained when linked to MetaCyc reactions; these labels indicate pathway provenance rather than quantitative profiling of those groups. Analyses focused on pathways related to exopolysaccharide production, biofilm formation, one-carbon metabolism, amino-acid biosynthesis, cofactor salvage, and redox buffering.

### 2.3. Bacterial Culture Identification and Genome Annotation

#### 2.3.1. Isolation

Root and leaf samples were surface-sterilized as described above, homogenized in 10 mL sterile water, diluted 1:10, and stored at 4°C for 7 days before 6 serial dilutions. The final three dilutions were plated on half-strength NA and starch nitrate agar (SNA) supplemented with cycloheximide (100 mg/L), dried at room temperature, and incubated inverted at 28°C. Colonies were purified by repeated sub-culturing, grouped by morphotype, and maintained on antibiotic-free Luria-Bertani (LB) agar (HiMedia).

#### 2.3.2. Initial Identification

Pure cultures were grown in LB broth at 30°C with agitation at 150 rpm for 48 hours, and DNA was extracted using the Xpress DNA Bacteria Kit (MagGenome) per the manufacturer’s protocol. DNA quantity and quality were assessed as previously described [4]. Isolates were initially identified by 16S rRNA gene amplification with primers 27F (AGAGTTTGATCMTGGCTCAG) and 1492R (GGTTACCTTGTTACGACTT) in 20 μL reactions using DreamTaq Polymerase (Thermo Fisher Scientific, California, USA). PCR products were analyzed on 1% agarose gels, purified with ExoSAP-IT (Applied Biosystems, Thermo Fisher Scientific) and sequenced by Sanger capillary sequencing using the BigDye Terminator v3.1 kit (Applied Biosystems) on an Applied Biosystems 3500 Series Genetic Analyzer. Consensus sequences were assembled in BioEdit [44], and homologous sequences were identified using BLASTn within BLAST (https://blast.ncbi.nlm.nih.gov/Blast.cgi).

#### 2.3.3. Whole-Genome Sequencing (WGS)

The DNA used for initial identification was also used to prepare Illumina and ONT WGS libraries. Reads were trimmed and hybrid assemblies generated as described previously [45], using Unicycler v.0.4.8 [46], MaSuRCA v.4.0.4 [47], and CANU v.2.2.[48]. Assemblies were polished with Pilon [49], assessed with BUSCO [50], and submitted to TYGS for species-level identification. For isolates flagged as potential new species, ANI was calculated against the closest TYGS/BLAST reference using the CJ Bioscience ANI calculator [51]. ANI values ≥95–96% were considered consistent with the same species, whereas lower values indicated potential novel species.

#### 2.3.4. Functional Annotation

Assembled genomes were annotated during GenBank submission using the NCBI Prokaryotic Annotation Pipeline [52]. Protein similarities were identified with BLASTp against UniProt [53, 54]. Metabolic pathway genes were detected via KEGG-KAAS using GHOSTZ [55] and mapped with KEGG Mapper. COG classifications were assigned using COGclassifier v.1.0.5, and Gene Ontology data were obtained from UniProt. Biosynthetic gene clusters (BGCs) were annotated with antiSMASH 7.0 [56], and OrthoVenn3 [57] was used to compare orthologous clusters with reference genomes.

### 2.4. Culture-based Biochemical Assays and Stress Tolerance

#### 2.4.1. Stress Tolerance

Drought tolerance was tested in LB broth supplemented with 10-30% PEG8000 [58]. Cultures were incubated overnight at 30°C and 150 rpm, and OD600 readings were used to classify isolates as sensitive (≤0.29), moderately tolerant (0.3–0.39), or tolerant (≥0.4). Salinity tolerance was assessed on LB agar containing 2-12% NaCl (w/v) in 2% increments and incubated for 7 days. Heat tolerance was tested at 22°C and 30–55°C in 5°C increments; plates with no growth after 7 days were transferred to 22°C to assess recovery. pH tolerance was tested in LB broth adjusted to pH 4-10 with 1 M HCl or NaOH, and the broth was incubated under the same conditions as the drought assay.

#### 2.4.2. Qualitative Enzyme Activity Assays

Qualitative screening used milk agar (HiMedia) for nitrogen dissolution [59], Pikovskaya’s agar (HiMedia) with 5% bromophenol blue for phosphate solubilization, M9 salts (HiMedia) with 1% (w/v) citrus pectin for pectinase, or 1% CMC for cellulase, starch agar (HiMedia) for amylase, Aleksandrow agar (HiMedia) for potassium solubilization, and nitrogen-free Dworkin and Foster’s salts minimal agar (DF; [60]) with 2 g (NH_4_)_2_SO_4_ or 3 mM ACC (Sigma-Aldrich, Maryland, USA) as sole nitrogen source. Plates were incubated at 35 for up to 7 days; pectinase, amylase, and cellulase activity were visualized with Gram’s iodine or Congo red. All assays were done in triplicate. IAA production was assessed in LB broth containing 2 g/L L-tryptophan after incubation at 37 and 150 rpm for 24 hours, followed by Salkowski’s reagent addition (2:1) and dark incubation for 30 min [61]; a red color indicated IAA production.

#### 2.4.3. Phytohormone Production

Spent LB culture medium was used for LC-MS/MS analysis to examine whether the bacteria can produce phytohormones associated with PGP. We tested for 6-benzylaminopurine (6-BA), naphthaleneacetic acid, IAA, salicylic acid (SA), indole-3-butyric acid (IBA), isopentenyl adenine (iPA), gibberellic acid (GA3), and abscisic acid (ABA) using available standards. The methodology followed is described in Supplementary Note S1.

### 2.5. Pangenome analysis of Pseudomonas orientalis

Publicly available *P. orientalis* genomes were downloaded from NCBI in FASTA format and analyzed with the B20 genome. Genomes were annotated with Prokka v.5.30.2 [62], assessed with BUSCO, and high-quality GFF3 files were used for Roary pan-genome analysis [63] at ≥95% BLASTp identity. The resulting gene presence/absence matrix was visualized in R [64]. Pan-genome openness was modeled using K = κ⋅n^−γ^ [65], where γ > 1 indicates a closed pan-genome and 0 < γ ≤ 1 an open pan-genome. TWILIGHT [66] was used to identify lineage-specific accessory-gene adaptations in complete free-living and plant-associated genomes, including B20. Genes were classified as core, accessory, intermediate, or unique, and phylogenetic mapping was used to assess selection and ecological associations. Pan-genome fluidity and diversity were estimated with ANGSD [67] using *P. aeruginosa* mutation rates (10 ¹¹), and effective population size was calculated as θ = 2Neμ. Lifestyle differences were tested using Wilcoxon tests in R (P < 0.05).

## 3. Results

### 3.1. Seasonal dynamics of microbial communities reflect environmental changes

#### 3.1.1. Amplicon profiling provides an exploratory overview of bacterial community structure

Amplicon profiling provided an exploratory overview of the bacterial community structure prior to shotgun metagenomic analysis. Plant-associated communities were shaped primarily by season and tissue type, with little effect of sampling location. Richness decreased from winter to summer in both leaves and roots, while ordination showed clear seasonal separation across plant tissues. In contrast, root-zone soil communities remained comparatively stable across seasons, although root-zone and bulk soil samples differed compositionally. Together, these patterns identified leaf and root tissues, particularly roots, as the most responsive compartments for downstream metagenomic analysis. Full amplicon diversity, ordination and taxonomic summaries are provided in the Supplement (Suppl. Notes S2–3 and Figures S1–4).

#### 3.1.2. Metagenomic profiling reveals root-associated seasonal restructuring

Shotgun metagenomic sequencing and Kraken2 classification were used to characterize bacterial genus-level profiles across leaf and root samples from both sites and seasons. Alpha diversity was summarized using observed genera, Shannon diversity, and Pielou evenness, with full seasonal comparison statistics provided in Supplementary Table S1. Leaf communities showed little seasonal change in diversity at either site and consistently had higher Shannon diversity and evenness than root communities. In contrast, root communities showed consistent summer declines in Shannon diversity and evenness at both sites, while observed genus richness remained comparatively stable (Figure 1A–C).

**Figure 1.**
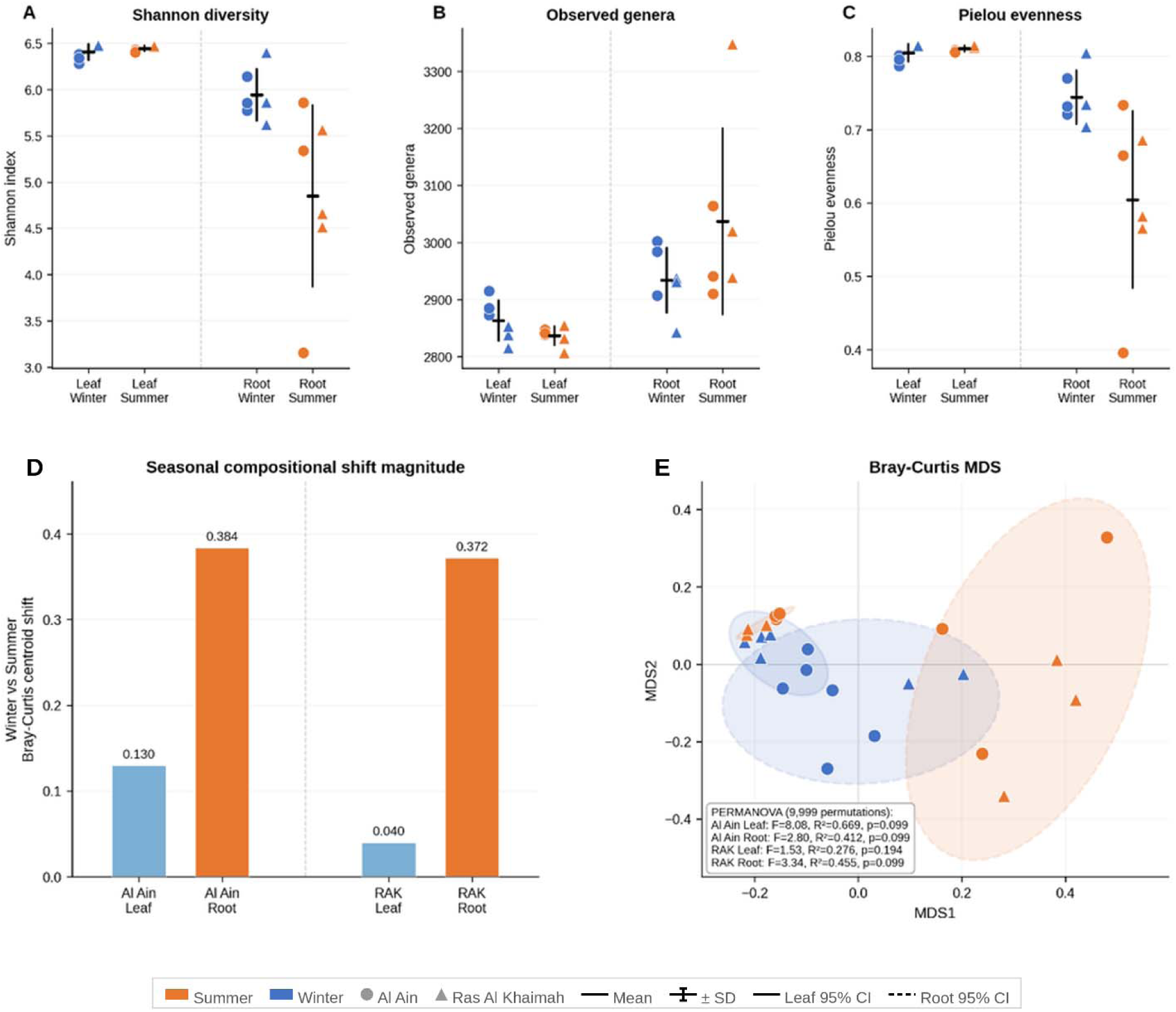
Diversity and compositional turnover of metagenome-derived bacterial communities across tissue types, sampling locations, and seasons. (A) Shannon diversity, (B) observed bacterial genera, and (C) Pielou evenness. Individual points represent biological replicates (n = 3 per group); horizontal bars and error bars indicate mean ± SD. (D) Winter-to-summer Bray-Curtis centroid shifts for each location–tissue subgroup. (E) Metric multidimensional scaling ordination based on Bray-Curtis dissimilarities. Dashed ellipses represent 95% confidence intervals for root communities and solid ellipses for leaf communities. Color indicates season (orange, summer; blue, winter), and marker shape indicates sampling location (circle, Al Ain; triangle, Ras Al Khaimah). PERMANOVA results were calculated using 9,999 permutations; full seasonal comparison statistics are provided in Supplementary Table S1.

Bray-Curtis-based beta diversity showed stronger seasonal compositional turnover in roots than in leaves. Winter-to-summer centroid shifts were larger in root communities at both Al Ain and Ras Al Khaimah, whereas leaf communities showed tighter cross-seasonal clustering, particularly in Ras Al Khaimah. PERMANOVA supported moderate-to-large seasonal effects in root communities, and MDS ordination confirmed greater seasonal separation in roots than leaves at both sites (Figure 1D–E; Suppl. Table S1).

#### 3.1.3. Taxonomic composition identified key taxa driving seasonal turnover across methods

Amplicon taxonomic profiles supported the exploratory diversity patterns and showed seasonal and tissue-associated bacterial *taxonomic* shifts; full phylum-and order-level summaries are provided in Supplementary Note S2 and Supplementary Figures S3–S4.

Shotgun metagenomes extended these bacterial patterns and provided broader taxonomic resolution across bacterial, fungal, and archaeal fractions. Most classified reads in leaf and root samples mapped to Eukaryota, whereas Bacteria dominated soil profiles (>50%; Suppl. Fig. S5). Viral assignments were excluded from downstream analyses, and unclassified and non-fungal eukaryotic assignments were removed to focus on microbial taxa relevant to this study. Among retained profiles, phylum-level distributions showed stronger seasonal than geographic differences, with Al Ain and RAK samples clustering more closely within seasons than between (Figure 2). Actinomycetota and Pseudomonadota dominated bacterial profiles, with Actinomycetota particularly abundant in soil (Figure 2A-D), while fungal profiles were dominated by Ascomycota and Basidiomycota (Figure 2E-H). Archaeal profiles showed comparatively little seasonal or geographic restructuring (Figure 2I-L). Because fungal and archaeal reads represented a minor fraction of the shotgun profiles, these groups were interpreted descriptively.

**Figure 2.**
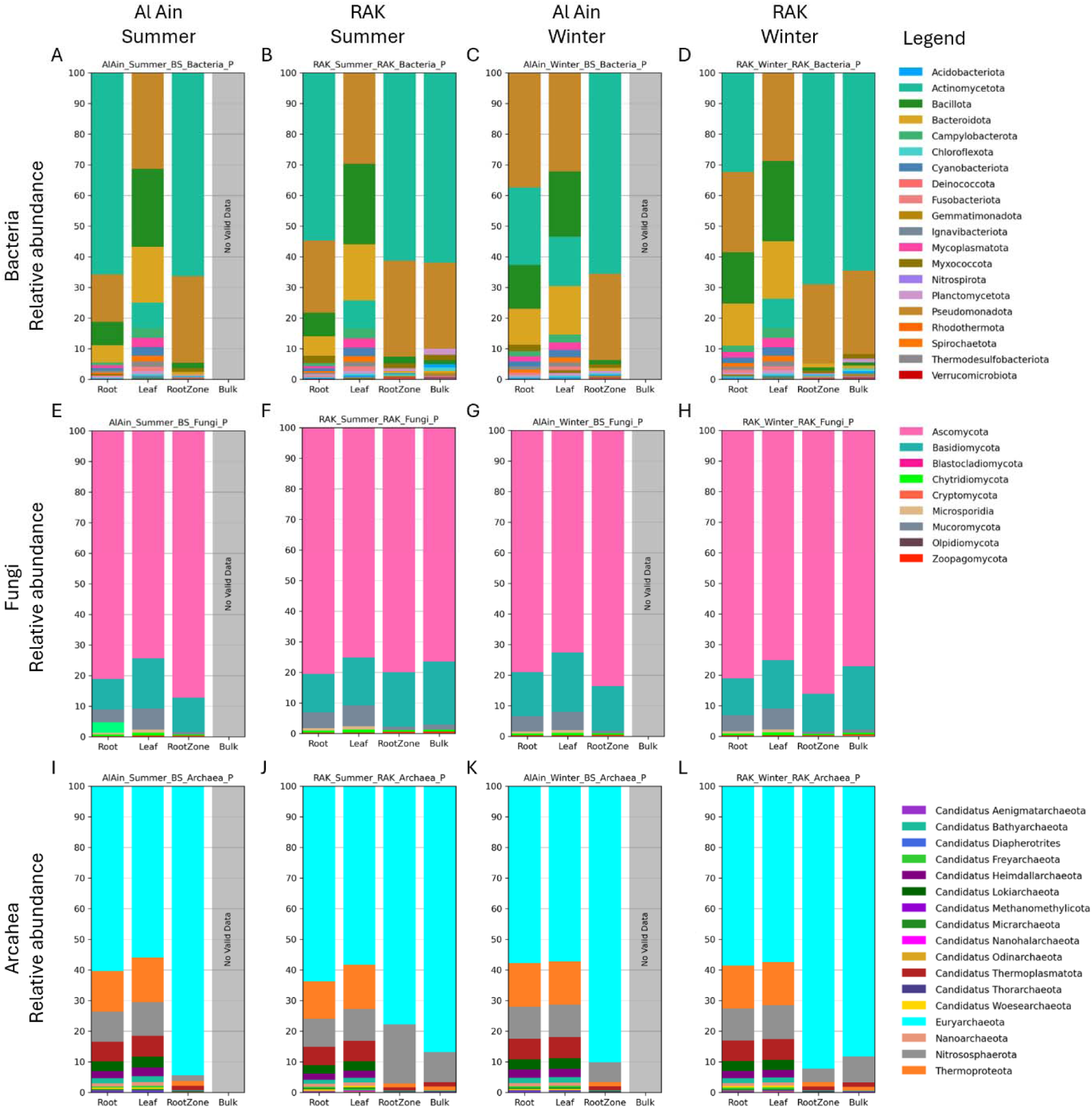
Phylum-level relative abundance of microbial communities in Al Ain and RAK samples based on metagenomic data. Bar graphs show the relative abundance of the top 15 bacterial (A–D), fungal (E–H), and archaeal (I–L) phyla across different sample types collected during summer and winter from Al Ain and Ras Al Khaimah (RAK). Each panel represents seasonal and site-specific variation within a specific microbial group.

Order-level profiles resolved the main bacterial turnover underlying these phylum-level patterns (Figure 3A-D). Pseudomonadales, Burkholderiales, Bacillales, Kitasatosporales, Pseudonocardiales, and Micrococcales showed the strongest seasonal variation. Leaves were enriched in Bacillales and Flavobacteriales, whereas roots and root-zone soils were dominated by Kitasatosporales, Pseudonocardiales, and Micrococcales. Seasonal restructuring was most evident in plant tissues, particularly roots, where Pseudonocardiales decreased from summer to winter in Al Ain and showed a similar, albeit weaker, pattern in RAK. Fungal order-level profiles also shifted descriptively (Figure 3E-H), with Pleosporales increasing from summer to winter across several tissues and sites, while Eurotiales and Sordariales declined in selected Al Ain root compartments.

**Figure 3.**
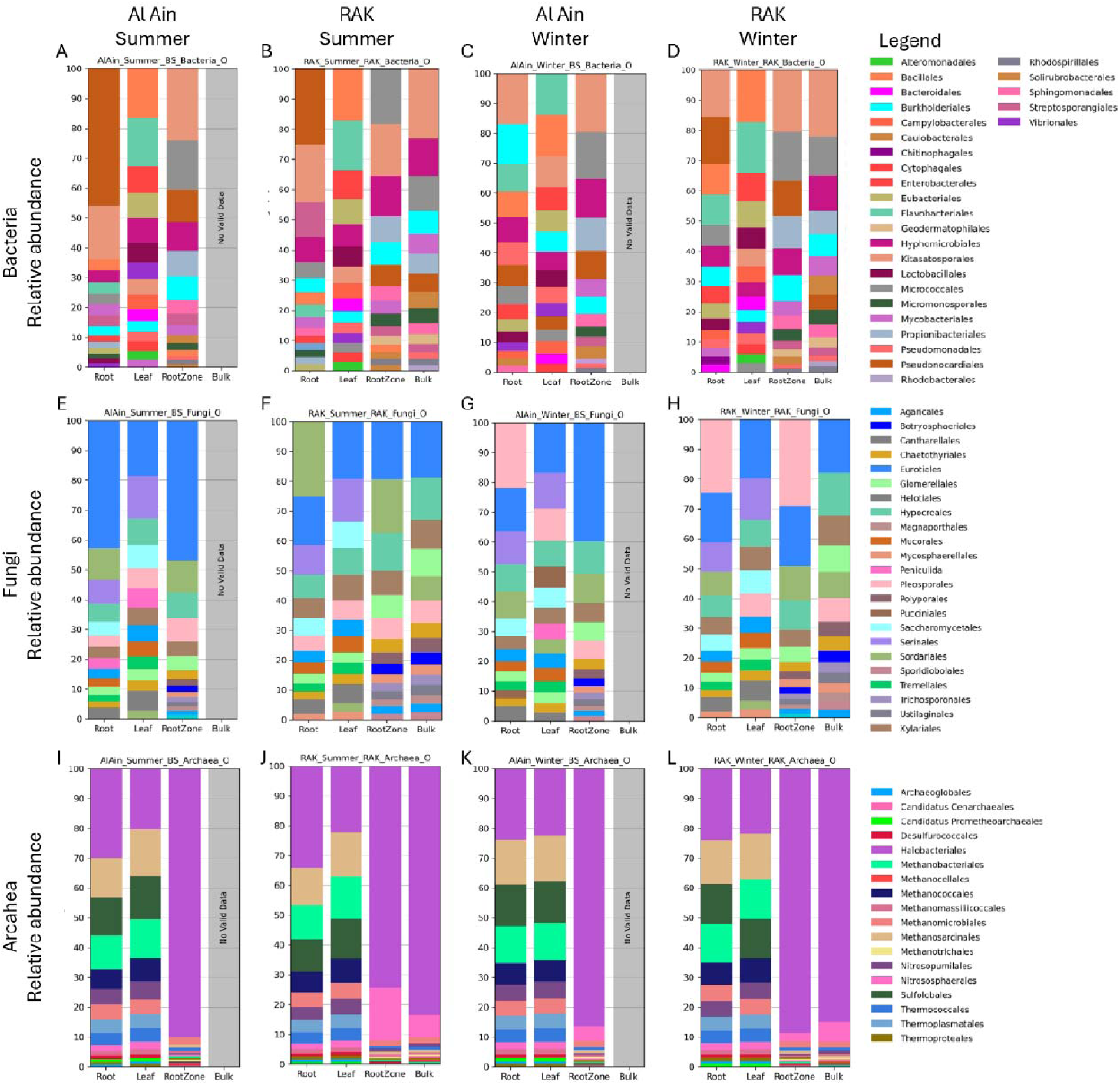
Order-level relative abundance of microbial communities in Al Ain and RAK samples based on metagenomic data. Bar graphs show the relative abundance of the top 15 bacterial (A–D), fungal (E–H), and archaeal (I–L) orders across different sample types collected during summer and winter from Al Ain and Ras Al Khaimah (RAK). Each panel represents seasonal and site-specific variation within a specifi microbial group.

At the genus level, seasonal changes were concentrated in root bacterial communities. Among the 40 most abundant bacterial genera analyzed, root communities showed the strongest seasonal shifts. Al Ain roots showed summer enrichment of *Saccharopolyspora* (log2FC = 3.91), *Amycolatopsis* (log2FC = 3.28), and *Pseudonocardia* (log2FC = 3.24), alongside declines in *Pseudomonas* (log2FC = −2.54) and *Massilia* (log2FC = −4.74). In Ras Al Khaimah roots, *Nonomuraea* (log2FC = 5.04), *Amycolatopsis* (log2FC = 1.82), and *Saccharopolyspora* (log2FC = 2.09) increased in summer, while *Clostridium*, *Vibrio*, and *Flavobacterium* declined. Leaf communities showed no consistent directional shifts in dominant genera at either site (Figure 4A), whereas root-specific seasonal shifts are shown in Figure 4B-C and Supplementary Table S2.

**Figure 4.**
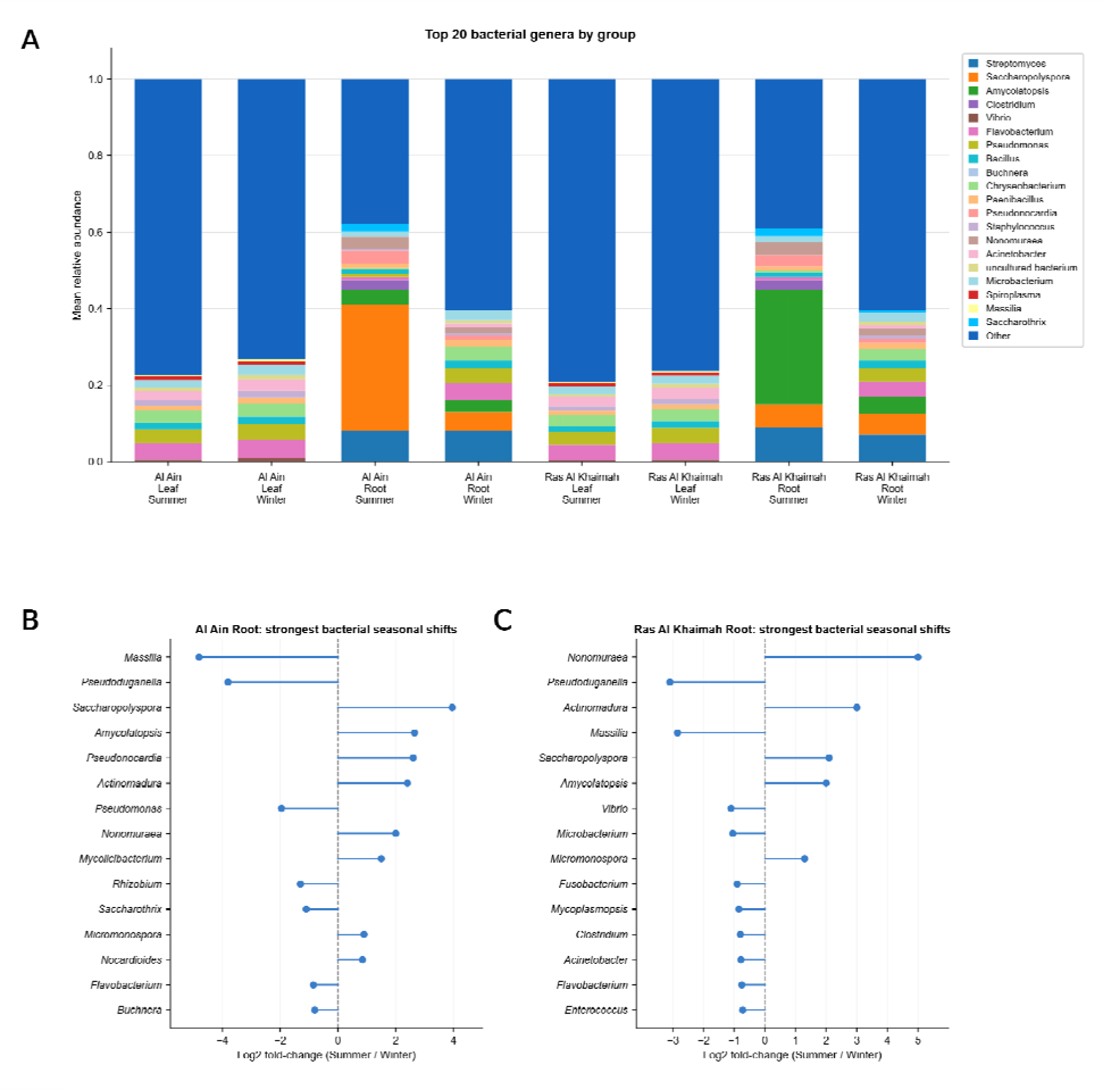
Dominant bacterial genera and seasonal shifts in root communities based on metagenomic data. (A) Mean relative abundance of the 20 most abundant bacterial genera across tissue types, sampling locations, and seasons. Each bar represents the group mean relative abundance (n = 3 biological replicates); remaining taxa are pooled as “Other.” (B–C) Seasonal log2 fold-changes in the relative abundance of dominant bacterial genera in root communities from (B) Al Ain and (C) Ras Al Khaimah. Positive values indicate summer enrichment and negative values indicate winter enrichment. Only genera with the strongest directional shifts are shown; full seasonal shift statistics for all 40 genera are provided in Supplementary Table S2.

### 3.1.4. Functional profiling of root communities shows seasonal restructuring

All 12 root metagenomic libraries produced usable functional profiles after filtering. Feature recovery differed across groups, with RAK summer samples showing the highest mean numbers of mapped enzyme classes, gene families, pathways, total reads, and mapped taxa, while RAK winter samples showed the lowest mean pathway recovery and the lowest mean mapped-taxon richness (Suppl. Table S3).

After the removal of unmapped and unclassified bins, the dominant named pathways were consistently central metabolic modules, including branched-chain amino acid biosynthesis, de novo biosynthesis of purine and guanosine nucleotides, the glyoxylate cycle, and TCA-associated routes (Figure 5A). These functions suggest maintenance of biosynthetic capacity, carbon flexibility, and nucleotide turnover across root communities, rather than reliance on a single stress-specialist pathway. Sample-level pathway variation was driven mainly by a strong deviation in the RAK winter sample RAK2R (Suppl. Fig. S6).

**Figure 5.**
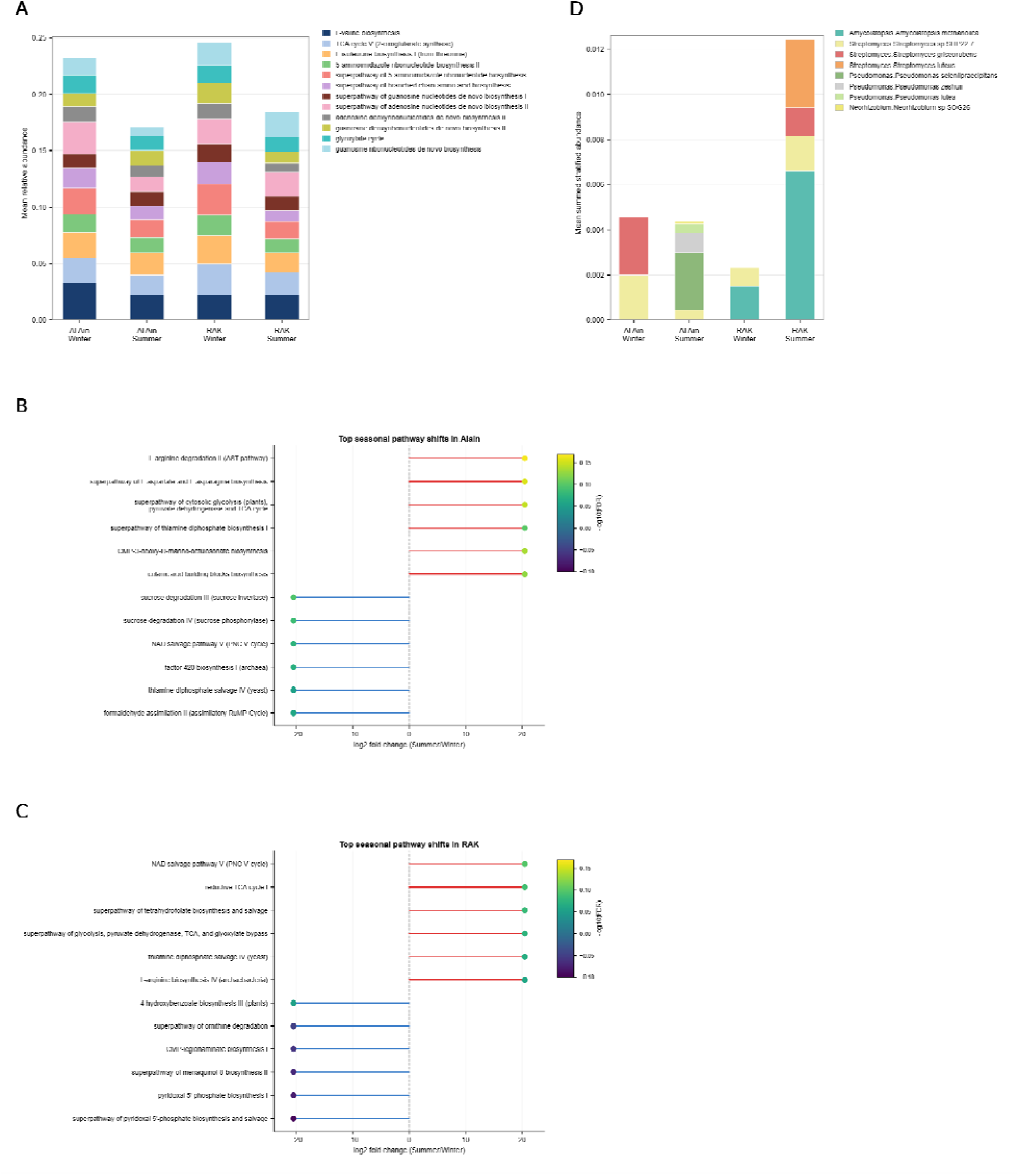
Functional pathway composition, seasonal shifts, and dominant bacterial contributors in root microbial communities. (A) Mean relative abundance of the 12 most abundant mapped functional pathways across sampling locations and seasons, based on HUMAnN pathway profiling after removal of unmapped and unclassified bins. (B–C) Seasonal pathway contrasts in root microbial communities from (B) Al Ain and (C) Ras Al Khaimah. Positive values indicate summer enrichment and negative value indicate winter enrichment; pathways are ranked by effect size within each site. (D) Mean summed stratified pathway abundance for dominant bacterial taxa contributing to functional profiles across sampling locations and seasons, based on HUMAnN stratified output. March and August correspond to winter and summer, respectively. Pathways and taxa are color-coded as shown in their respective legends.

Site-specific pathway contrasts revealed distinct seasonal functional trends in the roots of Al Ain and RAK (Figure 5B–C; Suppl. Table S4). In Al Ain, summer-enriched pathways included colanic acid building blocks and CMP-KDO biosynthesis, thiamine diphosphate biosynthesis, amino acid metabolism, and glucose/xylose degradation, consistent with envelope remodeling, exopolysaccharide production, carbon scavenging, and nitrogen metabolism. Winter-enriched pathways were associated with formaldehyde assimilation, tetrahydrofolate and NAD salvage, factor 420 biosynthesis, and sucrose degradation. In RAK, summer-enriched pathways were dominated by central carbon metabolism, cofactor maintenance, arginine biosynthesis, reductive TCA, NAD salvage, and formaldehyde assimilation, whereas winter-enriched pathways were linked to pyridoxal-5-phosphate, menaquinol-8, polyamine, and norspermidine metabolism. Because no individual pathway passed the stringent FDR threshold, these shifts were interpreted as coordinated, hypothesis-generating functional trends rather than confirmed pathway-level effects.

Stratified HUMAnN profiles identified the main bacterial contributors to these site-specific functional trends (Figure 5D; Suppl. Table S4). In Al Ain roots, functional contributions shifted from winter profiles dominated by *Streptomyces griseorubens* and *Streptomyces* sp. SHP22-7 to summer profiles dominated by *Pseudomonas* taxa, particularly *P. seleniipraecipitans*, *P. zeshuii*, and *P. lutea*, with a smaller *Neorhizobium* contribution. In RAK roots, summer profiles instead strengthened an actinobacterial signal, mainly from *Amycolatopsis methanolica*, *Streptomyces luteus*, *S. griseorubens*, and *Streptomyces* sp. SHP22-7. These contrasting contributors indicate that seasonal functional restructuring differed by site, with Al Ain associated with a more pseudomonad-rich profile and RAK with a stronger actinobacterial profile.

### 3.2. Cultured isolates mirror *in situ* taxa and express stress-ready, plant-beneficial functions

PCR-based 16S rRNA gene sequencing classified the 24 cultured isolates into the genera *Bacillus* (B04, B08, B11, B12, B14, B18, B19), *Pseudomonas* (B03, B05, B06, B10, B13, B15, B17, B20, B23, B26, B27), *Plantibacter* (B07), *Paenibacillus* (B09), *Brevibacillus* (B16), *Pantoea* (B22), *Streptomyces* (B24), and *Arthrobacter* (F02). These identifications informed the taxonomy used for subsequent assays and overlapped with lineages detected in the community profiles (Figure 2; Suppl. Fig. S3). Abiotic stress tolerance varied across isolates (Table 1). Several isolates grew under osmotic stress up to 20% PEG. Heat and salinity tolerance were concentrated in a smaller subset: B04 and B11 grew at 55 °C; B04, B11, and B14 grew at 10% NaCl; and B11 tolerated 12% NaCl. Most isolates grew at pH 6–8, whereas B22 grew across pH 4–10 and several *Bacillus* isolates (B05, B08, B12, B18, B19) tolerated pH 6–12. Overall, stress tolerance was distributed across multiple genera rather than restricted to a single lineage.

**Table 1.**
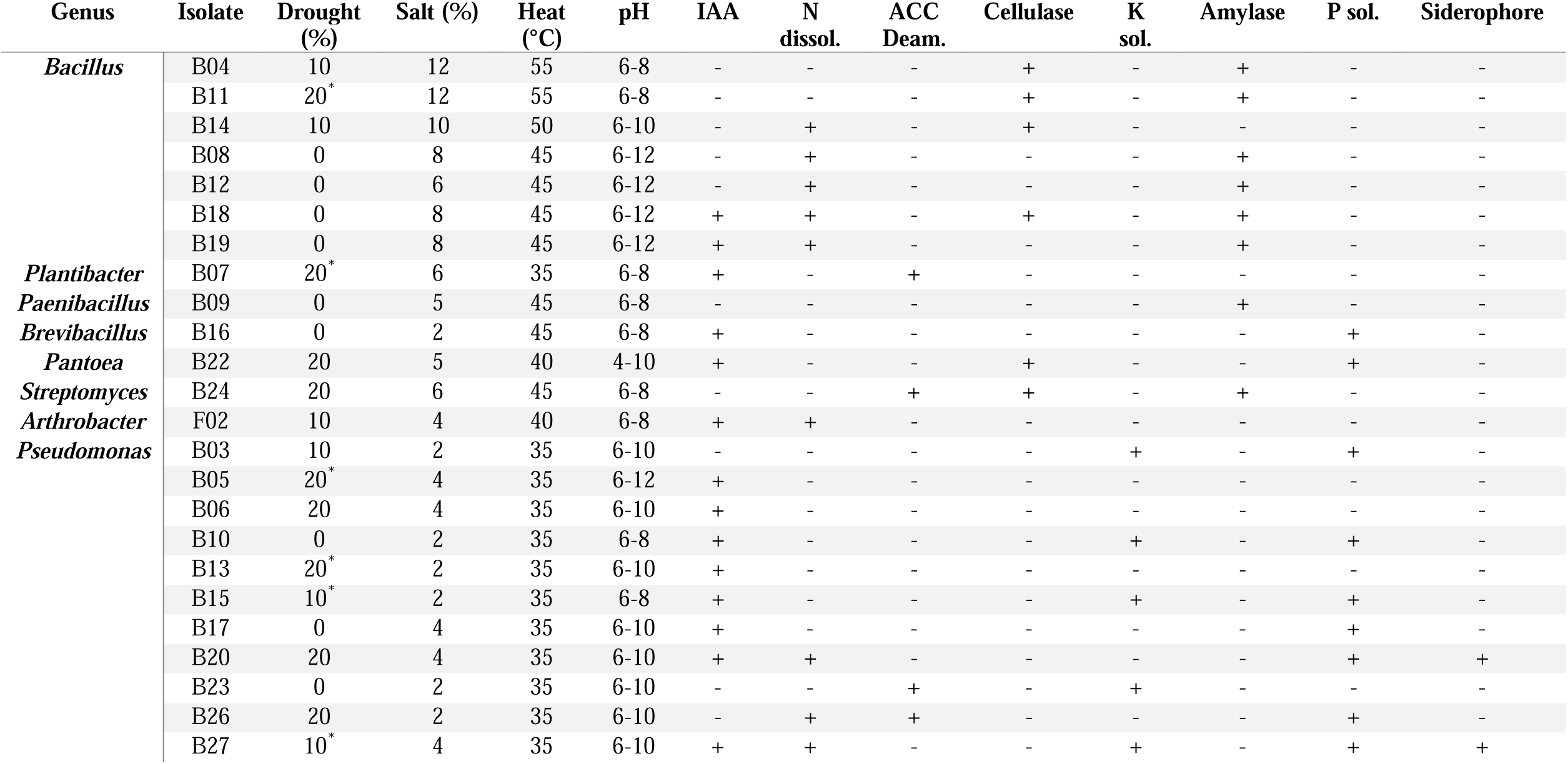
Overview of Stress Tolerance and PGP Assay Results.

| Genus | Isolate | Drought (%) | Salt (%) | Heat (°C) | pH | IAA | N dissol. | ACC Deam. | Cellulase | K sol. | Amylase | P sol. | Siderophore |
| --- | --- | --- | --- | --- | --- | --- | --- | --- | --- | --- | --- | --- | --- |
| *Bacillus* | B04 | 10 | 12 | 55 | 6-8 | - | - | - | + | - | + | - | - |
|  | B11 | 20* | 12 | 55 | 6-8 | - | - | - | + | - | + | - | - |
|  | B14 | 10 | 10 | 50 | 6-10 | - | + | - | + | - | - | - | - |
|  | B08 | 0 | 8 | 45 | 6-12 | - | + | - | - | - | + | - | - |
|  | B12 | 0 | 6 | 45 | 6-12 | - | + | - | - | - | + | - | - |
|  | B18 | 0 | 8 | 45 | 6-12 | + | + | - | + | - | + | - | - |
|  | B19 | 0 | 8 | 45 | 6-12 | + | + | - | - | - | + | - | - |
| *Plantibacter* | B07 | 20* | 6 | 35 | 6-8 | + | - | + | - | - | - | - | - |
| *Paenibacillus* | B09 | 0 | 5 | 45 | 6-8 | - | - | - | - | - | + | - | - |
| *Brevibacillus* | B16 | 0 | 2 | 45 | 6-8 | + | - | - | - | - | - | + | - |
| *Pantoea* | B22 | 20 | 5 | 40 | 4-10 | + | - | - | + | - | - | + | - |
| *Streptomyces* | B24 | 20 | 6 | 45 | 6-8 | - | - | + | + | - | + | - | - |
| *Arthrobacter* | F02 | 10 | 4 | 40 | 6-8 | + | + | - | - | - | - | - | - |
| *Pseudomonas* | B03 | 10 | 2 | 35 | 6-10 | - | - | - | - | + | - | + | - |
|  | B05 | 20* | 4 | 35 | 6-12 | + | - | - | - | - | - | - | - |
|  | B06 | 20 | 4 | 35 | 6-10 | + | - | - | - | - | - | - | - |
|  | B10 | 0 | 2 | 35 | 6-8 | + | - | - | - | + | - | + | - |
|  | B13 | 20* | 2 | 35 | 6-10 | + | - | - | - | - | - | - | - |
|  | B15 | 10* | 2 | 35 | 6-8 | + | - | - | - | + | - | + | - |
|  | B17 | 0 | 4 | 35 | 6-10 | + | - | - | - | - | - | + | - |
|  | B20 | 20 | 4 | 35 | 6-10 | + | + | - | - | - | - | + | + |
|  | B23 | 0 | 2 | 35 | 6-10 | - | - | + | - | + | - | - | - |
|  | B26 | 20 | 2 | 35 | 6-10 | - | + | + | - | - | - | + | - |
|  | B27 | 10* | 4 | 35 | 6-10 | + | + | - | - | + | - | + | + |

Plant growth–promoting activities were common but varied among isolates (Table 1). Nitrogen dissolution was observed in five *Bacillus* and two *Pseudomonas* isolates; ACC deaminase activity in B07, B24, and B26; cellulase activity in six isolates; amylase in eight isolates; and phosphate solubilization in ten isolates. Growth on CAS medium was frequent, although clearing halos were generally weak, with B20 and B27 showing the clearest zones. IAA screening showed strong positivity in B07 and B22, with variable results in other isolates. LC–MS/MS confirmed multiple phytohormones, with 6-BA and NAA below detection and consistently low ABA levels (Suppl. Fig. S7). Overall, enzyme, siderophore, and phytohormone-associated activities were distributed across several genera, with broader enzyme activity among *Bacillus* isolates and clearer siderophore-associated activity among selected *Pseudomonas* isolates.

Whole-genome sequencing confirmed species-level identities and resolved borderline assignments (Table 2; Suppl. Table S6). TYGS-based identification used digital DNA-DNA hybridization scores, followed by ANI analysis where needed. Four isolates initially assigned to *Bacillus anthracis* (B08, B12, B18, B19) were resolved as *B. thuringiensis*, consistent with ANI comparisons within the highly similar *B. cereus* group [24] and supported by motility assays on semi-solid agar. Among isolates flagged by TYGS as potential novel species (B03, B07, B17, B20, B23, B24, B26, B27, F02), only B03, B23, and F02 fell below the 95% ANI threshold, indicating potential novel *Pseudomonas* and *Arthrobacter* species; the remaining isolates showed ANI values ≥95% to known species (Table 2). Genome-derived context for these culture-based traits is presented in the following section.

**Table 2.**
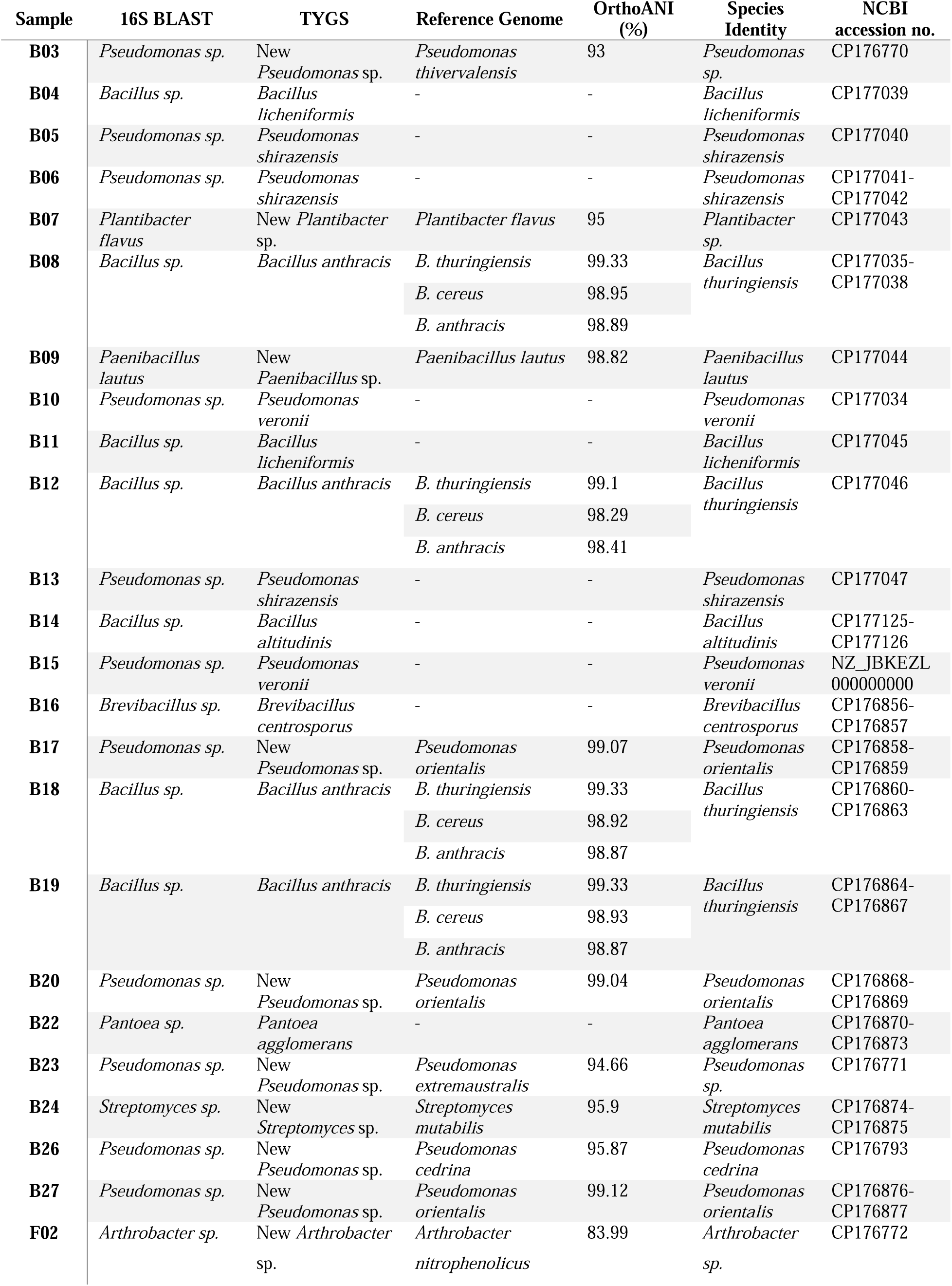
TYGS and OrthoANI Results for Bacterial Genomes and Assigned Species Identity.

### 3.3. Genome-resolved signals link observed phenotypes to adaptive gene content

Genome mining linked the cultured phenotypes to stress-and competition-related gene content. KEGG/COG profiling showed broad coverage of central metabolism, with *Pseudomonas* genomes enriched for secondary metabolite biosynthesis, biofilm formation, and two-component systems, whereas *Bacillus* genomes were dominated by core metabolic and cofactor pathways (Suppl. Table S7; Suppl. Fig. S8–S9). Other genera showed distinct patterns, including moderate secondary metabolism in *Streptomyces* and elevated signals in the nitrogen and carbohydrate pathways in *Pantoea* (Suppl. Table S7). BGC predictions identified lipopeptides in *Bacillus* and *Pseudomonas*, classical antibiotic clusters in *Streptomyces* and *Bacillus*, and siderophore clusters across multiple isolates [68, 69]. Stress-related loci included ectoine clusters in B09, B16, and B24 and carotenoid-associated clusters in B05, B06, B07, B13, B22, B24, and B26 (Suppl. Table S8) [70, 71]. OrthoVenn3 highlighted isolate-specific functions, including benzoate catabolism in B12, phosphonate metabolism in B23, and pyoverdine biosynthesis in B20, consistent with its siderophore activity in culture (Suppl. Table S9; Suppl. Figs. S10–S12).

Pangenome analysis of B20 with publicly available *P. orientalis* genomes (Suppl. Table S10) indicated an open pangenome (γ = 0.3313), with presence/absence matrices separating core and accessory modules (Figure 6A–B). Plant-associated genomes contained more lineage-specific accessory-intermediate genes than free-living genomes (Figure 6C). Several plant-associated accessory genes mapped to tryptophan-linked pathways, including tryptophan 2-monooxygenase and tryptophan decarboxylase, supporting hormone-related activity observed in culture for B20 (Suppl. Figs. S13–S14). Free-living strains showed higher nucleotide diversity than plant-associated strains (π = 0.021 vs 0.016; P < 0.005; Figure 6D) and included bacteriocin-protection genes, including a colicin-E3 immunity protein, that were absent from the plant-associated set (Suppl. Figs. S13–S14). Together, these patterns suggest that accessory content in *P. orientalis* differs with lifestyle, separating host-associated functions from environmental competition signals.

**Figure 6.**
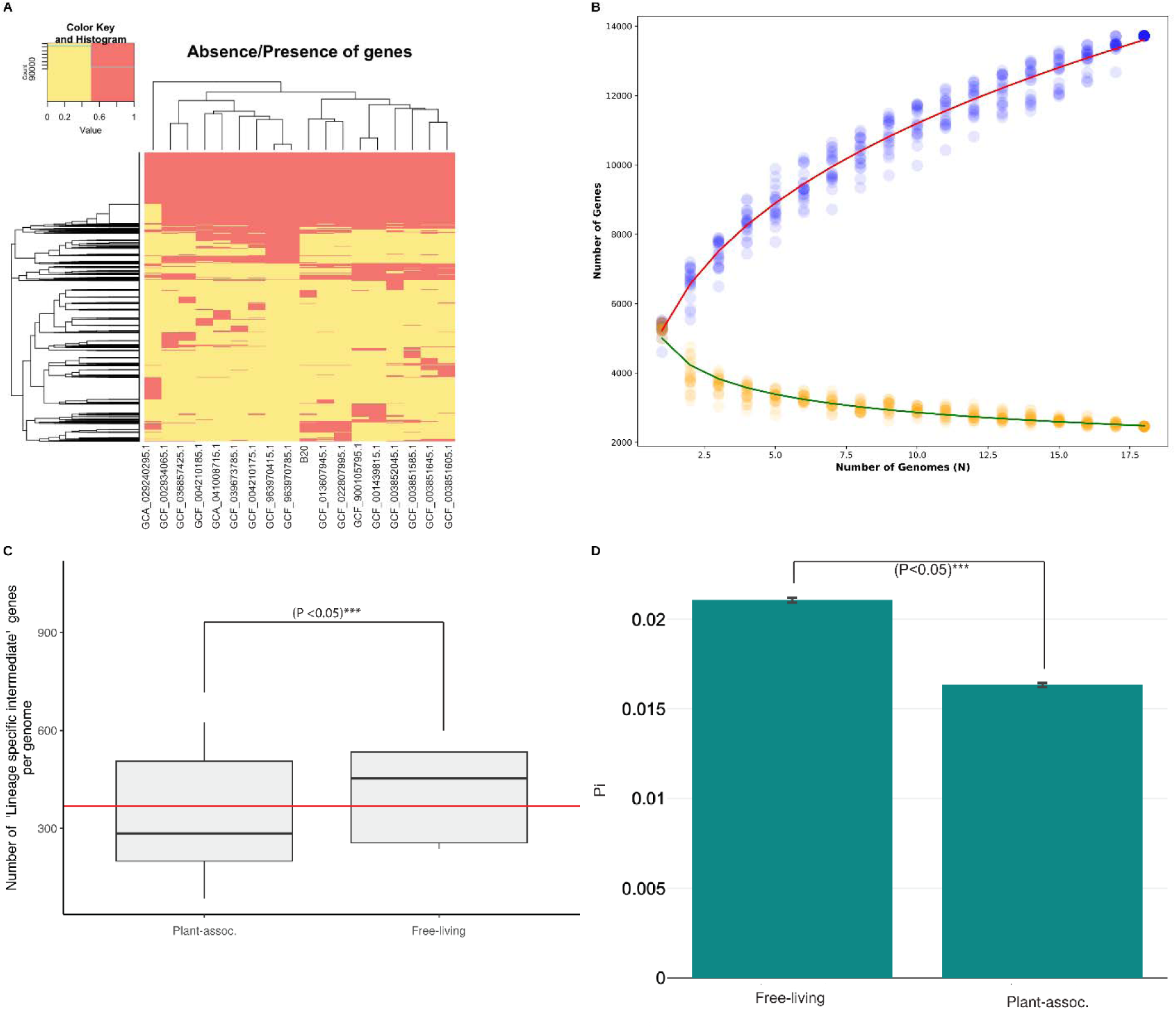
Pangenome analysis of *Pseudomonas orientalis* reveals an open pan-genome and distinct differences between plant-associated and free-living strains. (A) Heatmap displays gene presence and absence (yellow) across genomes, including isolate B20, with hierarchical clustering of core and accessory genes. (B) Curves show gene counts as genome size increases, with fitted trend lines indicating sustained gene acquisition (γ = 0.3313). (C) Bar plot compares lineage-specific accessory genes per genome, with higher counts in plant-associated strains. (D) Nucleotide diversity (π) is higher in free-living strains (π = 0.021) than in plant-associated strains (π = 0.016), indicating greater pan-genome fluidity.

## 4. Discussion

This study examined how the *Citrullus colocynthis* microbiome varies across seasons, tissues, and locations, and how these community patterns connect to stress-related functions. By combining an exploratory 16S rRNA amplicon screen with shotgun metagenomics, culture-based phenotyping, and genome analysis, the study links bacterial community structure to metagenomic taxonomic and functional shifts and to traits observed in representative cultured isolates. This integrated design extends prior single-season or culture-focused work on *C. colocynthis* by linking community dynamics to microbial functions relevant to desert persistence.

The amplicon data established the system’s initial ecological structure and guided the downstream metagenomic focus. Plant-associated bacterial communities were shaped mainly by season and tissue type, with little effect of sampling location; richness declined from winter to summer in leaves and roots, and ordination separated plant tissues by season. In contrast, root-zone soils were comparatively stable across seasons, but differed compositionally from bulk soils. These patterns identified leaves and roots, particularly roots, as the most responsive compartments for deeper metagenomic and functional analysis, while detailed amplicon results are provided in Supplementary Notes S2–3 and Supplementary Figures S1–S4.

Shotgun metagenomics extended the amplicon framework by providing genus-level bacterial resolution and broader microbial taxonomic coverage. Both approaches showed stronger seasonal responsiveness in roots than in leaves, although root Shannon diversity increased in the amplicon data while decreasing in the metagenomic profiles. This reflects the distinct units each method summarizes: ASV-level evenness across a broader low-abundance tail in the amplicon screen versus read-level dominance among classified genera in the shotgun data, where summer expansion of highly abundant actinobacterial genera suppressed effective diversity while observed genus richness remained stable. Together, the datasets that summer root communities retained detectable taxonomic breadth while becoming numerically dominated by stress-tolerant actinobacterial taxa.

Taxonomic results further linked the amplicon screen to the metagenomic findings. Amplicon profiles showed seasonal and tissue-associated bacterial shifts, including changes within Actinomycetota, while shotgun metagenomes confirmed Actinomycetota and Pseudomonadota as dominant bacterial phyla and resolved seasonal restructuring at order and genus levels. In roots, summer enrichment of actinobacterial genera such as *Saccharopolyspora*, *Amycolatopsis*, *Pseudonocardia*, and *Nonomuraea* coincided with declines in *Pseudomonas*, *Massilia*, *Flavobacterium*, and *Clostridium* (Figures 2–4), indicating a seasonal shift toward taxa associated with stress tolerance, secondary metabolism, and persistence under dry or nutrient-limited conditions[22, 72]. Fungal profiles showed descriptive order-level seasonal shifts, archaeal profiles were comparatively stable, and viral assignments were excluded. Similar host-linked adaptability has been reported in halophytes, cacti along aridity gradients, and native Arabian flora [73–76].

Shotgun pathway profiling provided an exploratory view of root functional potential across the available metagenomic root libraries (n = 12). Rather than interpreting individual pathway differences as statistically confirmed functional signatures, we interpreted the functional profiles through effect sizes, directional consistency, and convergence with taxonomic, isolate-genome, and phenotypic data. Under this framework, summer profiles did not collapse into a narrow stress response; dominant pathways remained centered on amino-acid biosynthesis, purine synthesis, glyoxylate shunts, and TCA-linked functions. This suggests maintenance of metabolic breadth during summer stress, consistent with *C. colocynthis* survival in sandy desert soils with episodic water availability and its fast-growing, water-spender strategy rather than the conservative strategy seen in evergreen desert shrubs or date palm [20, 77].

The functional carriers differed by site. In Al Ain, summer profiles shifted toward plant-beneficial *Pseudomonas* taxa, which are associated with exopolysaccharide production, biofilm formation, ACC-deaminase-mediated stress buffering, siderophore production, and root-barrier reinforcement under drought and heat [9, 78–80]. The matching Al Ain pathway signal, including colanic acid and CMP-KDO biosynthesis, thiamine diphosphate, amino-acid metabolism, and glucose/xylose degradation, is consistent with envelope investment, carbon scavenging, and anabolic maintenance. In RAK, summer profiles strengthened *Amycolatopsis methanolica* and *Streptomyces* taxa, consistent with the known association of endophytic actinobacteria with drought and heat adaptation, secondary metabolism, and root growth promotion [9, 81]. RAK summer pathways, including arginine biosynthesis, tetrahydrofolate biosynthesis and salvage, reductive TCA, NAD salvage, and formaldehyde assimilation, point to redox management, cofactor economy, and biosynthetic persistence. These site-specific contrasts should therefore be interpreted as coordinated, hypothesis-generating functional trends rather than confirmed pathway-level effects.

These contrasting profiles suggest that *C. colocynthis* roots may support site-specific summer consortia: a pseudomonad-rich, exopolysaccharide-centered profile in Al Ain and an actinobacterial, cofactor-and central-metabolism-centered profile in RAK. Despite taxonomic differences, both converge on maintenance of root metabolic activity and stress buffering under summer arid conditions, suggesting selection for functional capacity rather than a single taxonomic solution. The lifestyle-associated accessory genome variation observed in *Pseudomonas orientalis* provides one possible mechanism by which flexible bacterial lineages could contribute to different site-specific consortia while supporting convergent root functions.

This metagenome-first design was important because the cultured isolate collection complemented, rather than represented, the full root community. Community profiling captured dominant seasonal taxa regardless of culturability, including *Saccharopolyspora*, *Amycolatopsis*, *Nonomuraea*, and summer-root *Pseudomonas* taxa, while cultured isolates provided phenotypic and genome-resolved depth for representative lineages. *In silico* signals and *in vitro* traits both pointed to stress-protectant pathways, phytohormones, and biosynthetic gene clusters. Lipopeptide and siderophore BGCs in *Pseudomonas orientalis* (B17, B20, B27) and *Bacillus licheniformis* (B04) complemented observed drought, salinity, and heat tolerance, and plant growth-promoting activity (Table 1; Suppl. Table S8). Additional functions, including phytohormone production, phosphate solubilization, and siderophore biosynthesis, were observed in *Paenibacillus lautus* (B09), *Streptomyces mutabilis* (B24), and other isolates. Ectoine and carotenoid genes further support protection against osmotic and oxidative stress, consistent with arid-environment microbiomes [68–71, 82, 83].

Genome-resolved analysis refined these links. The *P. orientalis* pangenome was open and separated into core and accessory components (Figure 6). Plant-associated genomes contained accessory genes associated with tryptophan-dependent IAA pathways, including tryptophan 2-monooxygenase and tryptophan decarboxylase, which matched culture-level IAA signals and known *Pseudomonas* adaptations [28, 30, 31, 84]. Free-living genomes showed higher nucleotide diversity and included competition-related factors, such as a colicin-E3 immunity protein absent from the plant-associated group. These differences indicate that accessory genome content reflects host-associated versus free-living ecological demands within a single species.

Together, the links among seasonal community shifts (Figures 1–4), isolate phenotypes (Table 1), and genome-inferred capabilities highlight the value of desert-native microbiomes for arid agriculture. Stress-tolerant, growth-promoting bacteria can improve nutrient acquisition and crop performance under saline or drought-prone conditions [82, 83], and studies in the Arabian Peninsula show that native desert plants can harbor salt-tolerant, plant-beneficial endophytes [6, 73, 76, 85]. Several isolate–trait combinations point to testable applications under irrigation-limited, saline, and heat-stressed conditions: ectoine and carotenoid loci in B09, B16, and B24 suggest osmotic and oxidative stress-buffering potential; heat-and salt-tolerant *B. licheniformis* B04, carrying lipopeptide BGCs, provides a candidate for stress-resilient biocontrol; and *P. orientalis* B20/B27 combine drought tolerance, siderophore-associated activity, and accessory genome features linked to plant association. These traits should be evaluated in controlled *in planta* and soil-based assays using crops grown on marginal desert soils, where improved root growth, nutrient acquisition, and stress persistence are likely to be most relevant. In this context, *C. colocynthis* is both a model for desert holobiont function and a potential source of strains for microbiome-aware applications.

This study provides an expanded, multi-layered view of the *C. colocynthis* holobiont by extending previous single-season, single-location work to two sites and two seasons. The design prioritized integration across amplicon screening, shotgun metagenomics, culture-based phenotyping, and genome-resolved analysis. Biological replication was modest, and the interpretation was structured accordingly: amplicon screening identified roots as the primary focus, and cross-method convergence among metagenomics, isolate phenotyping, and genomics strengthened the main conclusions. Fungal and archaeal profiles were retained but were underrepresented, so targeted ITS, archaeal marker, or deeper metagenomic profiling will be useful for resolving their dynamics *in* future work.

Priority next steps include confirmed root colonization and controlled in planta evaluation of highlighted strains under field-relevant stress conditions, transcriptomic and metabolomic profiling to test the causal links implied by the functional data, and metagenome-guided isolation of dominant uncultured seasonal taxa including Saccharopolyspora and Amycolatopsis.

## 5. Conclusion

This study shows that bacterial communities associated with *Citrullus colocynthis* are structured primarily by season and tissue type, with roots showing the strongest seasonal restructuring. The 16S rRNA amplicon screen provided an exploratory framework for selecting the main metagenomic focus, while shotgun metagenomics extended the analysis to bacterial, fungal, and archaeal taxonomic profiles and showed that summer root communities were marked by bacterial taxonomic turnover and coordinated functional shifts. Culture-based assays and genome analyses linked representative isolates to stress tolerance, plant growth-promoting traits, and biosynthetic potential. In *Pseudomonas orientalis*, pangenome analysis further showed that accessory genome content differed between plant-associated and free-living lifestyles, implicating tryptophan-dependent IAA-related genes in host association and a bacteriocin-protection gene in environmental competition. Together, these results connect community composition, functional capacity, cultivable microbial traits, and lifestyle-associated genome variation in a desert plant holobiont, identifying desert-native bacteria and pathways to inform future microbiome-guided strategies for arid agriculture.

## 6. List of abbreviations

ABA: Abscisic Acid
ACC: 1-aminocyclopropane-1-carboxylate
ANI: Average Nucleotide Identity
ASV: Amplicon Sequence Variant
BGCs: Biosynthetic Gene Clusters
BUSCO: Benchmarking Universal Single-Copy Orthologs
CAS: Chrome azurol S
CFUs: Colony Forming Units
CMC: Carboxymethyl cellulose
COG: Cluster of orthologous gene
dDDH: Digital DNA-DNA Hybridization
DF: Dworkin and Foster
GA_3_: Gibberellic Acid
GO: Gene Ontology
IAA: Indole-3-Acetic Acid
IBA: Indole-3-Butyric Acid
ICNP: International Code of Nomenclature of Prokaryotes
IDs: Identities
iPA: Isopentyl Adenine
ITS: Internal Transcribed Spacer Region
KEGG: Kyoto Encyclopedia of Genes and Genomes
KO: KEGG Orthology
LB: Luria-Bertani
LC-MS/MS: Liquid chromatography-mass spectrometry/mass spectrometry NA Nutrient Agar
NMDS: Non-Metric Multidimensional Scaling
ONT: Oxford Nanopore Technologies
PCoA: Principal Coordinates Analysis
PDA: Potato Dextrose Agar
PDB: Potato Dextrose Broth
PE: Paired-End
PEG: Polyethylene Glycol
PGP: Plant Growth Promoting
PGPR: Plant Growth Promoting Rhizobacteria
RAK: Ras Al Khaimah
ROS: Reactive Oxygen Species
SA: Salicylic Acid
SNA: Starch Nitrate Agar
TYGS: Type (Strain) Genome Server
w/v: Weight per volume
WGS: Whole Genome Sequencing

## 7. Declarations

### 7.1. Authors’ contributions

Conceptualization and study design: KMH, KMAA; Methodology (Sample collection & processing): MP, BK, SA, FH, KA, IS, KMH, and KMAA; Analysis (Data curation): MP, NS, KMH, and KMAA; Funding acquisition: KMAA, KMH; Writing original draft: MP and KMH; Writing, review and editing: All.

## 7.2 Acknowledgements

The authors would like to thank and acknowledge the United Arab Emirates University (UAEU; Al Ain, UAE) and the Khalifa Center for Genetic Engineering and Biotechnology (KCGEB; Al Ain, UAE) for providing laboratory facilities and additional resources.

## 7.5. Availability of data and materials

The sequencing data generated for bacterial genomes during this study have been deposited in the NCBI-SRA database under the Bioproject ID: PRJNA1199959 and PRJNA1199998. The metagenome (16s amplicon (V3-V4)) data generated during this study have been submitted to the NCBI-SRA database under the BioProject id: PRJNA1345893.

