## Supplemental Notes, Tables, and Figures for "Desert Cucurbit Microbiomes: Spatiotemporal Dynamics and Functional Adaptations"

### Supplementary materials

### **Supplementary Notes**

**S1. Phytohormone screening**

*Extraction procedure*

500 uL of the bacterial sample was added to 50 uL of internal standard (Salicylic acid D4). Afterward, 2500 uL of 1% formic acid in 50% Acetonitrile was added and vortexed for 5 minutes, followed by centrifugation at 4500 rpm for 10 minutes. 1000 uL of the supernatant was removed and mixed with 1000 uL of mobile phase methanol and 0.01% formic acid in water (65:35, v/v). 10 µL was injected in the LC-MS/MS for analysis.

*LC-MS/MS*

LC-MS/MS analysis was conducted using a SHIMADZU LC-30AD (Nexera X2) binary pump and LCMS-8060 Shimadzu mass spectrometer, both manufactured in Japan. The separation was performed on a ZORBAX Eclipse Plus C18 column (100mm x 4.6 mm, 3.5µm) with an injection volume of 10µL. The autosampler temperature was set to 5ºC, and the column temperature was maintained at 30 ºC. The mass spectrometer operated in both ESI positive and Negative ion mode, with nitrogen used for both the heating and drying gas flow. The mobile phase consisted of methanol and 0.01% formic acid in water (65:35, v/v). The rinsing volume was 1500µL, with rinse modes before and after aspiration, using the rinse method of Rinse port only, with a total rinse time of 2 seconds. The total runtime for the analysis was 7.00 minutes. Retention times and MS/MS parameters for each plant hormone can be seen in Supplementary Table S5.

**S2. Amplicon-based Bacterial Diversity Patterns**

Amplicon sequencing produced 3,302,346 reads from leaf and root samples, which were grouped into 2,437 ASVs and rarefied to 115,691 reads per sample. Soil samples yielded 1,535,076 reads and 19,729 ASVs, with a rarefaction depth of 40,741 reads per sample.

Plant-associated bacterial communities showed clear seasonal structuring. Chao1 richness decreased from winter to summer in both leaves (winter 38.5 ± 4.7, summer 3.8 ± 1.7; Mann–Whitney U = 36, p = 0.005) and roots (winter 32.3 ± 7.7, summer 9.2 ± 3.3; U = 36, p = 0.005). Shannon diversity increased from winter to summer in roots (winter 0.423 ± 0.098, summer 0.713 ± 0.091; U = 36, p = 0.002), while leaf evenness did not differ significantly between seasons (U = 27, p = 0.180). Across plant tissues, roots had higher Shannon diversity than leaves overall (U = 122, p = 0.004; Supplementary Figure S1A–B). Jensen–Shannon divergence confirmed strong seasonal structuring of plant-associated communities. Season explained most compositional variation (PERMANOVA: pseudo-F = 14.24, R² = 0.719, p = 0.0008), followed by tissue type (pseudo-F = 4.72, R² = 0.179, p = 0.041), whereas sampling location had no significant effect (pseudo-F = 0.07, p = 0.806). NMDS ordination showed clear seasonal separation across tissue types and locations (stress = 0.016; Supplementary Figure S1C).

Soil bacterial communities were comparatively stable across seasons. Root-zone soil Shannon diversity did not differ significantly between winter and summer overall (winter 4.479 ± 0.294, summer 4.450 ± 0.122; U = 24, p = 0.394), or within Al Ain (U = 9, p = 0.100) and RAK (U = 2, p = 0.400). Chao1 richness was also stable across seasons (winter 395.9 ± 46.5, summer 376.7 ± 43.0; U = 26, p = 0.240). Root-zone soils had higher richness than bulk soils (386.3 ± 43.9 vs 313.1 ± 41.3; U = 33, p = 0.031), while bulk soils had higher Shannon diversity/evenness (4.686 ± 0.053 vs 4.465 ± 0.215; U = 4, p = 0.048). Because bulk soil sampling was limited to RAK and included only three samples, these comparisons were considered exploratory (Supplementary Figure S2A–B).

PERMANOVA detected no significant seasonal effect in soil communities (pseudo-F = 0.363, p = 0.865), but root-zone and bulk soil communities differed compositionally (pseudo-F = 10.722, R² = 0.541, p = 0.001). NMDS ordination showed no clear seasonal clustering but separated root-zone and bulk soil samples (stress = 0.086; Supplementary Figure S2C).

**S3. Amplicon-based Bacterial Taxonomic Patterns**

Amplicon-based taxonomic profiles showed tissue-, season-, and compartment-associated bacterial patterns. Leaf and root communities were dominated by Bacteroidota and Pseudomonadota, with lower contributions from Actinomycetota and Bacillota, whereas soil and root-zone communities were dominated by Actinomycetota, Pseudomonadota, and Bacillota. Soil-associated Bacteroidota and Chloroflexota varied by site and season, with contrasting root-zone trends between Al Ain and Ras Al Khaimah (Suppl. Fig. S3).

At finer taxonomic resolution, Actinomycetota orders showed stronger seasonal and sample-type variability than was evident at the phylum level (Suppl. Fig. S4). Mycobacteriales and Propionibacteriales were more abundant in summer, whereas Actinomycetales and Coriobacteriales increased in winter, with root-zone soils often showing contrasting patterns. These order-level shifts supported the use of amplicon profiles as an exploratory screen and provided context for the metagenomic taxonomic analyses.

### **Supplementary Tables**

Supplementary Table S1: Seasonal alpha diversity comparisons for metagenome-derived bacterial communities across tissue types and sampling locations.

| **Location** | **Tissue** | **Metric** | **Winter mean** | **Summer mean** | **Δ (Summer − Winter)** | **Cliff's δ** | **Mann-Whitney U** | **p-value** | **FDR** |
| --- | --- | --- | --- | --- | --- | --- | --- | --- | --- |
| Al Ain | Leaf | Shannon diversity | 6.334 | 6.42 | 0.086 | 1 | 0 | 0.1 | 0.2 |
| Al Ain | Leaf | Observed genera | 2891 | 2842.333 | -48.667 | -1 | 9 | 0.1 | 0.2 |
| Al Ain | Leaf | Pielou evenness | 0.795 | 0.807 | 0.012 | 1 | 0 | 0.1 | 0.2 |
| Al Ain | Root | Shannon diversity | 5.924 | 4.784 | -1.14 | -0.556 | 7 | 0.4 | 0.533 |
| Al Ain | Root | Observed genera | 2964.333 | 2971.667 | 7.333 | 0.111 | 4 | 1 | 1 |
| Al Ain | Root | Pielou evenness | 0.741 | 0.598 | -0.143 | -0.556 | 7 | 0.4 | 0.533 |
| Ras Al Khaimah | Leaf | Shannon diversity | 6.481 | 6.473 | -0.008 | -0.556 | 7 | 0.4 | 0.533 |
| Ras Al Khaimah | Leaf | Observed genera | 2835.333 | 2831.333 | -4 | -0.111 | 5 | 1 | 1 |
| Ras Al Khaimah | Leaf | Pielou evenness | 0.815 | 0.814 | -0.001 | -0.111 | 5 | 1 | 1 |
| Ras Al Khaimah | Root | Shannon diversity | 5.962 | 4.916 | -1.046 | -1 | 9 | 0.1 | 0.2 |
| Ras Al Khaimah | Root | Observed genera | 2904 | 3102.333 | 198.333 | 1 | 0 | 0.1 | 0.2 |
| Ras Al Khaimah | Root | Pielou evenness | 0.748 | 0.611 | -0.137 | -1 | 9 | 0.1 | 0.2 |

Supplementary Table S2: Seasonal shifts in relative abundance of the 40 most abundant bacterial genera across tissue types and sampling locations.

| **Location** | **Tissue** | **Genus** | **Winter mean rel. abund.** | **Summer mean rel. abund.** | **log₂FC (Summer vs Winter)** | **Cliff's δ** | **Mann-Whitney U** | **p-value** | **FDR** |
| --- | --- | --- | --- | --- | --- | --- | --- | --- | --- |
| Al Ain | Leaf | *Bradyrhizobium* | 0.0043 | 0.0095 | 1.157 | 1 | 0 | 0.1 | 0.186 |
| Al Ain | Leaf | uncultured bacterium | 0.0066 | 0.0118 | 0.826 | 1 | 0 | 0.1 | 0.186 |
| Al Ain | Leaf | *Vibrio* | 0.0254 | 0.0308 | 0.277 | 0.778 | 1 | 0.2 | 0.305 |
| Al Ain | Leaf | *Mycoplasmopsis* | 0.0041 | 0.0049 | 0.257 | 1 | 0 | 0.1 | 0.186 |
| Al Ain | Leaf | *Helicobacter* | 0.0044 | 0.0052 | 0.257 | 1 | 0 | 0.1 | 0.186 |
| Al Ain | Leaf | *Campylobacter* | 0.0057 | 0.0068 | 0.253 | 1 | 0 | 0.1 | 0.186 |
| Al Ain | Leaf | *Spiroplasma* | 0.0069 | 0.0083 | 0.253 | 1 | 0 | 0.1 | 0.186 |
| Al Ain | Leaf | *Nostoc* | 0.004 | 0.0048 | 0.24 | 1 | 0 | 0.1 | 0.186 |
| Al Ain | Leaf | *Chryseobacterium* | 0.0109 | 0.0129 | 0.238 | 1 | 0 | 0.1 | 0.186 |
| Al Ain | Leaf | *Streptococcus* | 0.0053 | 0.0062 | 0.237 | 1 | 0 | 0.1 | 0.186 |
| Al Ain | Leaf | *Polaribacter* | 0.0059 | 0.0069 | 0.227 | 1 | 0 | 0.1 | 0.186 |
| Al Ain | Leaf | *Staphylococcus* | 0.0097 | 0.0114 | 0.227 | 1 | 0 | 0.1 | 0.186 |
| Al Ain | Leaf | *Bacteroides* | 0.0052 | 0.0061 | 0.219 | 1 | 0 | 0.1 | 0.186 |
| Al Ain | Leaf | *Bacillus* | 0.013 | 0.0151 | 0.217 | 1 | 0 | 0.1 | 0.186 |
| Al Ain | Leaf | *Acinetobacter* | 0.0091 | 0.0105 | 0.209 | 1 | 0 | 0.1 | 0.186 |
| Al Ain | Leaf | *Clostridium* | 0.0309 | 0.0356 | 0.204 | 1 | 0 | 0.1 | 0.186 |
| Al Ain | Leaf | *Enterococcus* | 0.0049 | 0.0055 | 0.183 | 1 | 0 | 0.1 | 0.186 |
| Al Ain | Leaf | *Fusobacterium* | 0.0047 | 0.0054 | 0.182 | 1 | 0 | 0.1 | 0.186 |
| Al Ain | Leaf | *Bacteroidales* | 0.0049 | 0.0056 | 0.178 | 1 | 0 | 0.1 | 0.186 |
| Al Ain | Leaf | *Sphingobacterium* | 0.0043 | 0.0048 | 0.175 | 1 | 0 | 0.1 | 0.186 |
| Al Ain | Leaf | *Flavobacterium* | 0.0195 | 0.022 | 0.174 | 1 | 0 | 0.1 | 0.186 |
| Al Ain | Leaf | *Paenibacillus* | 0.0106 | 0.0119 | 0.173 | 1 | 0 | 0.1 | 0.186 |
| Al Ain | Leaf | *Shewanella* | 0.0049 | 0.0054 | 0.161 | 1 | 0 | 0.1 | 0.186 |
| Al Ain | Leaf | *Prevotella* | 0.0035 | 0.0037 | 0.098 | 0.778 | 1 | 0.2 | 0.305 |
| Al Ain | Leaf | *Mycolicibacterium* | 0.0038 | 0.0035 | -0.124 | -1 | 9 | 0.1 | 0.186 |
| Al Ain | Leaf | *Buchnera* | 0.0142 | 0.0125 | -0.187 | 0.333 | 3 | 0.7 | 0.752 |
| Al Ain | Leaf | *Micromonospora* | 0.0027 | 0.0022 | -0.308 | -1 | 9 | 0.1 | 0.186 |
| Al Ain | Leaf | *Amycolatopsis* | 0.0024 | 0.0019 | -0.368 | -1 | 9 | 0.1 | 0.186 |
| Al Ain | Leaf | *Actinomadura* | 0.0011 | 0.0008 | -0.401 | -1 | 9 | 0.1 | 0.186 |
| Al Ain | Leaf | *Rhizobium* | 0.0039 | 0.0027 | -0.566 | -1 | 9 | 0.1 | 0.186 |
| Al Ain | Leaf | *Nocardioides* | 0.0027 | 0.0014 | -0.932 | -1 | 9 | 0.1 | 0.186 |
| Al Ain | Leaf | *Streptomyces* | 0.0468 | 0.0227 | -1.042 | -1 | 9 | 0.1 | 0.186 |
| Al Ain | Leaf | *Pseudomonas* | 0.0246 | 0.012 | -1.042 | -1 | 9 | 0.1 | 0.186 |
| Al Ain | Leaf | *Microbacterium* | 0.0054 | 0.0021 | -1.367 | -1 | 9 | 0.1 | 0.186 |
| Al Ain | Leaf | *Nonomuraea* | 0.0019 | 0.0004 | -2.222 | -1 | 9 | 0.1 | 0.186 |
| Al Ain | Leaf | *Pseudonocardia* | 0.0017 | 0.0004 | -2.257 | -1 | 9 | 0.1 | 0.186 |
| Al Ain | Leaf | *Pseudoduganella* | 0.0043 | 0.0005 | -3.13 | -1 | 9 | 0.1 | 0.186 |
| Al Ain | Leaf | *Massilia* | 0.0063 | 0.0007 | -3.268 | -1 | 9 | 0.1 | 0.186 |
| Al Ain | Leaf | *Saccharopolyspora* | 0.0068 | 0.0003 | -4.293 | -1 | 9 | 0.1 | 0.186 |
| Al Ain | Leaf | *Saccharothrix* | 0.0083 | 0.0002 | -5.644 | -1 | 9 | 0.1 | 0.186 |
| Al Ain | Root | *Saccharopolyspora* | 0.016 | 0.2398 | 3.906 | 0.778 | 1 | 0.2 | 0.305 |
| Al Ain | Root | *Amycolatopsis* | 0.0029 | 0.0279 | 3.277 | 1 | 0 | 0.1 | 0.186 |
| Al Ain | Root | *Pseudonocardia* | 0.0025 | 0.0235 | 3.236 | 1 | 0 | 0.1 | 0.186 |
| Al Ain | Root | *Actinomadura* | 0.0011 | 0.0071 | 2.686 | 1 | 0 | 0.1 | 0.186 |
| Al Ain | Root | *Nonomuraea* | 0.0032 | 0.0129 | 2.002 | 0.333 | 3 | 0.7 | 0.752 |
| Al Ain | Root | *Mycolicibacterium* | 0.0036 | 0.0082 | 1.204 | 1 | 0 | 0.1 | 0.186 |
| Al Ain | Root | *Micromonospora* | 0.0033 | 0.0065 | 0.986 | 1 | 0 | 0.1 | 0.186 |
| Al Ain | Root | *Nocardioides* | 0.0045 | 0.0088 | 0.97 | 0.778 | 1 | 0.2 | 0.305 |
| Al Ain | Root | *Streptomyces* | 0.0852 | 0.102 | 0.259 | 0.333 | 3 | 0.7 | 0.752 |
| Al Ain | Root | *Bradyrhizobium* | 0.0042 | 0.0043 | 0.052 | -0.333 | 6 | 0.7 | 0.752 |
| Al Ain | Root | uncultured bacterium | 0.0044 | 0.0033 | -0.426 | -0.556 | 7 | 0.4 | 0.478 |
| Al Ain | Root | *Vibrio* | 0.019 | 0.0129 | -0.553 | -0.556 | 7 | 0.4 | 0.478 |
| Al Ain | Root | *Microbacterium* | 0.009 | 0.006 | -0.582 | -0.556 | 7 | 0.4 | 0.478 |
| Al Ain | Root | *Clostridium* | 0.0216 | 0.0142 | -0.61 | -0.556 | 7 | 0.4 | 0.478 |
| Al Ain | Root | *Spiroplasma* | 0.0049 | 0.0032 | -0.625 | -0.556 | 7 | 0.4 | 0.478 |
| Al Ain | Root | *Campylobacter* | 0.0041 | 0.0026 | -0.632 | -0.556 | 7 | 0.4 | 0.478 |
| Al Ain | Root | *Bacteroides* | 0.0038 | 0.0024 | -0.637 | -0.556 | 7 | 0.4 | 0.478 |
| Al Ain | Root | *Helicobacter* | 0.0032 | 0.002 | -0.643 | -0.556 | 7 | 0.4 | 0.478 |
| Al Ain | Root | *Staphylococcus* | 0.007 | 0.0045 | -0.646 | -0.556 | 7 | 0.4 | 0.478 |
| Al Ain | Root | *Mycoplasmopsis* | 0.003 | 0.0019 | -0.649 | -0.556 | 7 | 0.4 | 0.478 |
| Al Ain | Root | *Enterococcus* | 0.0034 | 0.0022 | -0.659 | -0.556 | 7 | 0.4 | 0.478 |
| Al Ain | Root | *Bacillus* | 0.0094 | 0.0059 | -0.662 | -0.556 | 7 | 0.4 | 0.478 |
| Al Ain | Root | *Polaribacter* | 0.0043 | 0.0027 | -0.663 | -0.556 | 7 | 0.4 | 0.478 |
| Al Ain | Root | *Chryseobacterium* | 0.0081 | 0.005 | -0.681 | -0.778 | 8 | 0.2 | 0.305 |
| Al Ain | Root | *Streptococcus* | 0.0038 | 0.0024 | -0.685 | -0.556 | 7 | 0.4 | 0.478 |
| Al Ain | Root | *Fusobacterium* | 0.0034 | 0.0021 | -0.686 | -0.556 | 7 | 0.4 | 0.478 |
| Al Ain | Root | *Nostoc* | 0.003 | 0.0018 | -0.7 | -0.556 | 7 | 0.4 | 0.478 |
| Al Ain | Root | *Shewanella* | 0.0035 | 0.0022 | -0.701 | -0.778 | 8 | 0.2 | 0.305 |
| Al Ain | Root | *Paenibacillus* | 0.0079 | 0.0048 | -0.703 | -0.778 | 8 | 0.2 | 0.305 |
| Al Ain | Root | *Acinetobacter* | 0.0066 | 0.004 | -0.707 | -0.778 | 8 | 0.2 | 0.305 |
| Al Ain | Root | *Bacteroidales* | 0.0036 | 0.0022 | -0.74 | -1 | 9 | 0.1 | 0.186 |
| Al Ain | Root | *Prevotella* | 0.0025 | 0.0015 | -0.762 | -0.778 | 8 | 0.2 | 0.305 |
| Al Ain | Root | *Sphingobacterium* | 0.0032 | 0.0019 | -0.775 | -1 | 9 | 0.1 | 0.186 |
| Al Ain | Root | *Buchnera* | 0.009 | 0.0048 | -0.916 | -1 | 9 | 0.1 | 0.186 |
| Al Ain | Root | *Flavobacterium* | 0.0168 | 0.0086 | -0.965 | -1 | 9 | 0.1 | 0.186 |
| Al Ain | Root | *Saccharothrix* | 0.0172 | 0.0085 | -1.008 | -0.556 | 7 | 0.4 | 0.478 |
| Al Ain | Root | *Rhizobium* | 0.0063 | 0.0029 | -1.101 | -0.556 | 7 | 0.4 | 0.478 |
| Al Ain | Root | *Pseudomonas* | 0.0419 | 0.0072 | -2.536 | -1 | 9 | 0.1 | 0.186 |
| Al Ain | Root | *Pseudoduganella* | 0.0135 | 0.0007 | -4.214 | -1 | 9 | 0.1 | 0.186 |
| Al Ain | Root | *Massilia* | 0.0261 | 0.001 | -4.744 | -1 | 9 | 0.1 | 0.186 |
| Ras Al Khaimah | Leaf | *Bradyrhizobium* | 0.0039 | 0.0077 | 0.976 | 1 | 0 | 0.1 | 0.186 |
| Ras Al Khaimah | Leaf | uncultured bacterium | 0.0076 | 0.0099 | 0.387 | 1 | 0 | 0.1 | 0.186 |
| Ras Al Khaimah | Leaf | *Nonomuraea* | 0.0005 | 0.0005 | 0.087 | 0.556 | 2 | 0.4 | 0.478 |
| Ras Al Khaimah | Leaf | *Spiroplasma* | 0.0082 | 0.0086 | 0.072 | 0.778 | 1 | 0.2 | 0.305 |
| Ras Al Khaimah | Leaf | *Prevotella* | 0.0067 | 0.0069 | 0.059 | -0.111 | 5 | 1 | 1 |
| Ras Al Khaimah | Leaf | *Polaribacter* | 0.0072 | 0.0074 | 0.028 | 0.333 | 3 | 0.7 | 0.752 |
| Ras Al Khaimah | Leaf | *Staphylococcus* | 0.012 | 0.0122 | 0.024 | 0.333 | 3 | 0.7 | 0.752 |
| Ras Al Khaimah | Leaf | *Shewanella* | 0.0057 | 0.0058 | 0.016 | 0.333 | 3 | 0.7 | 0.752 |
| Ras Al Khaimah | Leaf | *Bacillus* | 0.0153 | 0.0154 | 0.01 | -0.111 | 5 | 1 | 1 |
| Ras Al Khaimah | Leaf | *Paenibacillus* | 0.0126 | 0.0127 | 0.007 | 0.111 | 4 | 1 | 1 |
| Ras Al Khaimah | Leaf | *Mycoplasmopsis* | 0.005 | 0.005 | 0.003 | -0.333 | 6 | 0.7 | 0.752 |
| Ras Al Khaimah | Leaf | *Chryseobacterium* | 0.013 | 0.013 | 0.001 | -0.111 | 5 | 1 | 1 |
| Ras Al Khaimah | Leaf | *Campylobacter* | 0.0068 | 0.0068 | -0.001 | -0.111 | 5 | 1 | 1 |
| Ras Al Khaimah | Leaf | *Bacteroidales* | 0.0061 | 0.0061 | -0.006 | -0.333 | 6 | 0.7 | 0.752 |
| Ras Al Khaimah | Leaf | *Mycolicibacterium* | 0.0039 | 0.0038 | -0.008 | -0.111 | 5 | 1 | 1 |
| Ras Al Khaimah | Leaf | *Buchnera* | 0.0123 | 0.0122 | -0.012 | -1 | 9 | 0.1 | 0.186 |
| Ras Al Khaimah | Leaf | *Rhizobium* | 0.0028 | 0.0027 | -0.016 | -0.333 | 6 | 0.7 | 0.752 |
| Ras Al Khaimah | Leaf | *Sphingobacterium* | 0.0048 | 0.0048 | -0.017 | -0.111 | 5 | 1 | 1 |
| Ras Al Khaimah | Leaf | *Acinetobacter* | 0.0106 | 0.0105 | -0.019 | -1 | 9 | 0.1 | 0.186 |
| Ras Al Khaimah | Leaf | *Flavobacterium* | 0.022 | 0.0216 | -0.027 | -1 | 9 | 0.1 | 0.186 |
| Ras Al Khaimah | Leaf | *Streptococcus* | 0.0063 | 0.0062 | -0.028 | -0.333 | 6 | 0.7 | 0.752 |
| Ras Al Khaimah | Leaf | *Nostoc* | 0.0048 | 0.0047 | -0.031 | -0.333 | 6 | 0.7 | 0.752 |
| Ras Al Khaimah | Leaf | *Enterococcus* | 0.0055 | 0.0054 | -0.032 | -1 | 9 | 0.1 | 0.186 |
| Ras Al Khaimah | Leaf | *Clostridium* | 0.0352 | 0.0342 | -0.041 | -0.333 | 6 | 0.7 | 0.752 |
| Ras Al Khaimah | Leaf | *Nocardioides* | 0.0014 | 0.0014 | -0.041 | -0.111 | 5 | 1 | 1 |
| Ras Al Khaimah | Leaf | *Pseudomonas* | 0.013 | 0.0126 | -0.043 | -0.778 | 8 | 0.2 | 0.305 |
| Ras Al Khaimah | Leaf | *Fusobacterium* | 0.0062 | 0.006 | -0.059 | -0.556 | 7 | 0.4 | 0.478 |
| Ras Al Khaimah | Leaf | *Micromonospora* | 0.0021 | 0.002 | -0.074 | -0.556 | 7 | 0.4 | 0.478 |
| Ras Al Khaimah | Leaf | *Helicobacter* | 0.0055 | 0.0051 | -0.092 | -1 | 9 | 0.1 | 0.186 |
| Ras Al Khaimah | Leaf | *Bacteroides* | 0.006 | 0.0057 | -0.092 | -0.778 | 8 | 0.2 | 0.305 |
| Ras Al Khaimah | Leaf | *Streptomyces* | 0.0251 | 0.0232 | -0.108 | -1 | 9 | 0.1 | 0.186 |
| Ras Al Khaimah | Leaf | *Microbacterium* | 0.0022 | 0.002 | -0.116 | -0.556 | 7 | 0.4 | 0.478 |
| Ras Al Khaimah | Leaf | *Actinomadura* | 0.0009 | 0.0008 | -0.19 | -1 | 9 | 0.1 | 0.186 |
| Ras Al Khaimah | Leaf | *Vibrio* | 0.0181 | 0.0156 | -0.211 | -0.778 | 8 | 0.2 | 0.305 |
| Ras Al Khaimah | Leaf | *Massilia* | 0.0007 | 0.0006 | -0.223 | -0.556 | 7 | 0.4 | 0.478 |
| Ras Al Khaimah | Leaf | *Amycolatopsis* | 0.0027 | 0.0021 | -0.371 | -0.556 | 7 | 0.4 | 0.478 |
| Ras Al Khaimah | Leaf | *Pseudoduganella* | 0.0005 | 0.0004 | -0.398 | -0.778 | 8 | 0.2 | 0.305 |
| Ras Al Khaimah | Leaf | *Saccharothrix* | 0.0003 | 0.0002 | -0.516 | -0.778 | 8 | 0.2 | 0.305 |
| Ras Al Khaimah | Leaf | *Saccharopolyspora* | 0.0005 | 0.0003 | -0.548 | -0.778 | 8 | 0.2 | 0.305 |
| Ras Al Khaimah | Leaf | *Pseudonocardia* | 0.0006 | 0.0003 | -0.822 | -0.556 | 7 | 0.4 | 0.478 |
| Ras Al Khaimah | Root | *Nonomuraea* | 0.0014 | 0.0464 | 5.044 | 1 | 0 | 0.1 | 0.186 |
| Ras Al Khaimah | Root | *Actinomadura* | 0.0018 | 0.0149 | 3.017 | 0.778 | 1 | 0.2 | 0.305 |
| Ras Al Khaimah | Root | *Saccharopolyspora* | 0.0073 | 0.0312 | 2.09 | 1 | 0 | 0.1 | 0.186 |
| Ras Al Khaimah | Root | *Amycolatopsis* | 0.0564 | 0.1986 | 1.818 | 0.556 | 2 | 0.4 | 0.478 |
| Ras Al Khaimah | Root | *Micromonospora* | 0.0032 | 0.0075 | 1.233 | 1 | 0 | 0.1 | 0.186 |
| Ras Al Khaimah | Root | *Bradyrhizobium* | 0.0037 | 0.0066 | 0.839 | 1 | 0 | 0.1 | 0.186 |
| Ras Al Khaimah | Root | *Pseudonocardia* | 0.0171 | 0.0265 | 0.63 | 0.111 | 4 | 1 | 1 |
| Ras Al Khaimah | Root | *Mycolicibacterium* | 0.0046 | 0.0065 | 0.48 | 0.556 | 2 | 0.4 | 0.478 |
| Ras Al Khaimah | Root | *Streptomyces* | 0.0733 | 0.0945 | 0.367 | 0.333 | 3 | 0.7 | 0.752 |
| Ras Al Khaimah | Root | *Nocardioides* | 0.0071 | 0.0084 | 0.246 | 0.111 | 4 | 1 | 1 |
| Ras Al Khaimah | Root | *Rhizobium* | 0.0041 | 0.0032 | -0.387 | -0.556 | 7 | 0.4 | 0.478 |
| Ras Al Khaimah | Root | *Saccharothrix* | 0.007 | 0.0047 | -0.561 | -0.333 | 6 | 0.7 | 0.752 |
| Ras Al Khaimah | Root | *Pseudomonas* | 0.0117 | 0.0078 | -0.576 | -0.778 | 8 | 0.2 | 0.305 |
| Ras Al Khaimah | Root | uncultured bacterium | 0.0047 | 0.0029 | -0.678 | -1 | 9 | 0.1 | 0.186 |
| Ras Al Khaimah | Root | *Bacteroides* | 0.0042 | 0.0022 | -0.953 | -0.778 | 8 | 0.2 | 0.305 |
| Ras Al Khaimah | Root | *Paenibacillus* | 0.0088 | 0.0044 | -0.997 | -1 | 9 | 0.1 | 0.186 |
| Ras Al Khaimah | Root | *Prevotella* | 0.0045 | 0.0022 | -1.016 | -1 | 9 | 0.1 | 0.186 |
| Ras Al Khaimah | Root | *Sphingobacterium* | 0.0034 | 0.0016 | -1.047 | -1 | 9 | 0.1 | 0.186 |
| Ras Al Khaimah | Root | *Bacteroidales* | 0.0041 | 0.002 | -1.051 | -1 | 9 | 0.1 | 0.186 |
| Ras Al Khaimah | Root | *Bacillus* | 0.0107 | 0.0049 | -1.108 | -1 | 9 | 0.1 | 0.186 |
| Ras Al Khaimah | Root | *Shewanella* | 0.0039 | 0.0018 | -1.111 | -1 | 9 | 0.1 | 0.186 |
| Ras Al Khaimah | Root | *Chryseobacterium* | 0.0091 | 0.0041 | -1.142 | -1 | 9 | 0.1 | 0.186 |
| Ras Al Khaimah | Root | *Nostoc* | 0.0033 | 0.0015 | -1.146 | -1 | 9 | 0.1 | 0.186 |
| Ras Al Khaimah | Root | *Buchnera* | 0.0084 | 0.0038 | -1.154 | -1 | 9 | 0.1 | 0.186 |
| Ras Al Khaimah | Root | *Polaribacter* | 0.005 | 0.0022 | -1.156 | -1 | 9 | 0.1 | 0.186 |
| Ras Al Khaimah | Root | *Spiroplasma* | 0.0058 | 0.0026 | -1.164 | -1 | 9 | 0.1 | 0.186 |
| Ras Al Khaimah | Root | *Staphylococcus* | 0.0083 | 0.0037 | -1.167 | -1 | 9 | 0.1 | 0.186 |
| Ras Al Khaimah | Root | *Streptococcus* | 0.0044 | 0.002 | -1.171 | -1 | 9 | 0.1 | 0.186 |
| Ras Al Khaimah | Root | *Helicobacter* | 0.0037 | 0.0016 | -1.176 | -1 | 9 | 0.1 | 0.186 |
| Ras Al Khaimah | Root | *Campylobacter* | 0.0047 | 0.0021 | -1.182 | -1 | 9 | 0.1 | 0.186 |
| Ras Al Khaimah | Root | *Enterococcus* | 0.0038 | 0.0017 | -1.184 | -1 | 9 | 0.1 | 0.186 |
| Ras Al Khaimah | Root | *Flavobacterium* | 0.0157 | 0.0069 | -1.193 | -1 | 9 | 0.1 | 0.186 |
| Ras Al Khaimah | Root | *Acinetobacter* | 0.0074 | 0.0032 | -1.2 | -1 | 9 | 0.1 | 0.186 |
| Ras Al Khaimah | Root | *Clostridium* | 0.0247 | 0.0107 | -1.203 | -1 | 9 | 0.1 | 0.186 |
| Ras Al Khaimah | Root | *Mycoplasmopsis* | 0.0035 | 0.0015 | -1.204 | -1 | 9 | 0.1 | 0.186 |
| Ras Al Khaimah | Root | *Fusobacterium* | 0.0041 | 0.0018 | -1.223 | -1 | 9 | 0.1 | 0.186 |
| Ras Al Khaimah | Root | *Microbacterium* | 0.017 | 0.0072 | -1.239 | -0.111 | 5 | 1 | 1 |
| Ras Al Khaimah | Root | *Vibrio* | 0.0125 | 0.0049 | -1.35 | -1 | 9 | 0.1 | 0.186 |
| Ras Al Khaimah | Root | *Massilia* | 0.0098 | 0.0014 | -2.825 | -1 | 9 | 0.1 | 0.186 |
| Ras Al Khaimah | Root | *Pseudoduganella* | 0.008 | 0.0009 | -3.091 | -0.556 | 7 | 0.4 | 0.478 |

Supplementary Table S3: Quality-control summary of filtered HUMAnN feature recovery by location-season group.

| **Group** | **n** | **Mean ECs** | **Min ECs** | **Mean gene families** | **Min gene families** | **Mean pathways** | **Min pathways** | **Mean reads** | **Mean mapped taxa** |
| --- | --- | --- | --- | --- | --- | --- | --- | --- | --- |
| Al Ain Winter | 3 | 1,026.7 | 647 | 163,939.3 | 60,562 | 206.0 | 109 | 1,195,730.5 | 25.0 |
| Al Ain Summer | 3 | 1,169.0 | 1,014 | 189,433.3 | 130,555 | 250.3 | 199 | 465,891.7 | 24.0 |
| RAK Winter | 3 | 891.0 | 405 | 131,390.0 | 15,685 | 178.7 | 51 | 458,765.0 | 16.0 |
| RAK Summer | 3 | 1,211.3 | 1,025 | 241,591.3 | 143,139 | 272.7 | 229 | 1,163,331.7 | 63.7 |

Supplementary Table S4: Representative high-effect pathway shifts within Al Ain and RAK after filtering.

| **Location** | **Pathway** | **log2FC (Aug/Mar)** | **CLR effect** | **FDR** |
| --- | --- | --- | --- | --- |
| Al Ain | colanic acid building blocks biosynthesis | 20.22 | 14.01 | 0.634 |
| Al Ain | CMP-3-deoxy-D-manno-octulosonate biosynthesis | 20.16 | 13.97 | 0.634 |
| Al Ain | superpathway of thiamine diphosphate biosynthesis I | 19.53 | 13.54 | 0.875 |
| Al Ain | superpathway of cytosolic glycolysis (plants), pyruvate dehydrogenase and TCA cycle | 19.47 | 13.49 | 0.634 |
| Al Ain | superpathway of L-aspartate and L-asparagine biosynthesis | 19.46 | 13.49 | 0.634 |
| Al Ain | formaldehyde assimilation II (assimilatory RuMP Cycle) | -19.90 | -13.80 | 0.634 |
| Al Ain | thiamine diphosphate salvage IV (yeast) | -19.43 | -13.47 | 0.875 |
| Al Ain | factor 420 biosynthesis I (archaea) | -19.33 | -13.40 | 0.634 |
| Al Ain | NAD salvage pathway V (PNC V cycle) | -19.05 | -13.21 | 0.634 |
| Al Ain | sucrose degradation IV (sucrose phosphorylase) | -18.67 | -12.94 | 0.634 |
| RAK | L-arginine biosynthesis IV (archaebacteria) | 20.96 | 14.53 | 1.000 |
| RAK | thiamine diphosphate salvage IV (yeast) | 19.42 | 13.46 | 1.000 |
| RAK | superpathway of glycolysis, pyruvate dehydrogenase, TCA, and glyoxylate bypass | 19.37 | 13.43 | 1.000 |
| RAK | superpathway of tetrahydrofolate biosynthesis and salvage | 19.19 | 13.30 | 1.000 |
| RAK | reductive TCA cycle I | 19.15 | 13.27 | 1.000 |
| RAK | superpathway of pyridoxal 5'-phosphate biosynthesis and salvage | -19.14 | -13.27 | 1.000 |
| RAK | pyridoxal 5'-phosphate biosynthesis I | -18.43 | -12.78 | 1.000 |
| RAK | superpathway of menaquinol-8 biosynthesis II | -18.09 | -12.54 | 1.000 |
| RAK | CMP-legionaminate biosynthesis I | -17.88 | -12.39 | 1.000 |
| RAK | superpathway of ornithine degradation | -16.63 | -11.53 | 1.000 |

Supplementary Table S5: The optimum conditions and retention time for plant hormones by LC-MS/MS.

| **Compound names** | **Retention time** | **Q1** | **Q3** | **Dwell** | **Q1 pre Bias** | **CE** | **Q3 pre Bias** | **Ionization** |
| --- | --- | --- | --- | --- | --- | --- | --- | --- |
| Indole-3-acetic acid | 2.921 | 176.000 | 130.100 | 100 | -17 | -15 | -22 | Positive |
|  |  | 176.000 | 77.100 | 100 | -16 | -42 | -15 |  |
|  |  | 176.000 | 103.050 | 100 | -16 | -29 | -23 |  |
| Isopentenyl adenine | 3.545 | 204.150 | 136.000 | 100 | -20 | -15 | -20 | Positive |
|  |  | 204.150 | 148.000 | 100 | -10 | -13 | -14 |  |
|  |  | 204.150 | 119.000 | 100 | -10 | -31 | -11 |  |
| Naphthalene acetic acid | 5.07 | 184.900 | 117.050 | 100 | 22 | 11 | 26 | Negative |
|  |  | 184.900 | 100.050 | 100 | 20 | 34 | 38 |  |
|  |  | 184.900 | 141.100 | 100 | 22 | 10 | 20 |  |
| Indole-3-butyric acid | 3.523 | 202.050 | 134.150 | 100 | 22 | 16 | 20 | Negative |
|  |  | 202.050 | 133.150 | 100 | 22 | 22 | 20 |  |
|  |  | 202.050 | 132.200 | 100 | 22 | 30 | 28 |  |
| 6-Benzyl aminopurine | 3.381 | 224.050 | 133.150 | 100 | 24 | 23 | 26 | Negative |
|  |  | 224.050 | 132.150 | 100 | 24 | 32 | 26 |  |
|  |  | 224.050 | 188.000 | 100 | 26 | 13 | 22 |  |
| Gibberalic acid | 2.412 | 345.250 | 143.200 | 100 | 17 | 29 | 13 | Negative |
|  |  | 345.250 | 239.200 | 100 | 17 | 16 | 30 |  |
|  |  | 345.250 | 221.350 | 100 | 17 | 25 | 22 |  |
| Salicylic acid | 3.096 | 137.200 | 93.150 | 100 | 13 | 21 | 11 | Negative |
|  |  | 137.200 | 65.200 | 100 | 14 | 28 | 15 |  |
|  |  | 137.200 | 100.000 | 100 | 15 | 27 | 26 |  |
| Abscisic acid | 3.34 | 263.100 | 153.250 | 100 | 28 | 12 | 20 | Negative |
|  |  | 263.100 | 219.250 | 100 | 28 | 14 | 24 |  |
|  |  | 263.100 | 204.300 | 100 | 28 | 20 | 26 |  |
| Salicylic acid D4  (Internal standard) | 2.901 | 141.100 | 97.100 | 100 | 14 | 21 | 12 | Negative |
|  |  | 141.100 | 59.100 | 100 | 14 | 12 | 14 |  |
|  |  | 141.100 | 69.150 | 100 | 14 | 30 | 8 |  |

Supplementary Table S6. Information on the Bacterial Genome Sequencing Data Generated

| **Isolate** | **Technology** | **Bases** | **Sum of bases** | **Genome size (bp)** | **x Coverage** | **N50** | **%GC** | **BUSCO Completeness %** | **Plasmid(s)** |
| --- | --- | --- | --- | --- | --- | --- | --- | --- | --- |
| B03 | WGS (Illumina) | 1793869800 | 2591108820 | 6508542 | 398 | 6508542 | 61 | 100 |  |
|  | WGS (Nanopore) | 797239020 |  |  |  |  |  |  |  |
| B04 | WGS (Illumina) | 1295876400 | 2798185423 | 4332882 | 646 | 4332882 | 46 | 100 |  |
|  | WGS (Nanopore) | 1502309023 |  |  |  |  |  |  |  |
| B05 | WGS (Illumina) | 1758603600 | 2701274910 | 5310272 | 509 | 5310272 | 62 | 100 |  |
|  | WGS (Nanopore) | 942671310 |  |  |  |  |  |  |  |
| B06 | WGS (Illumina) | 1773381900 | 2366985759 | 5255261 | 450 | 5255261 | 62 | 100 | 1 |
|  | WGS (Nanopore) | 593603859 |  |  |  |  |  |  |  |
| B07 | WGS (Illumina) | 1795218600 | 3360508698 | 4059395 | 828 | 4059395 | 69 | 96.8 (F:1, M:1.6) |  |
|  | WGS (Nanopore) | 1565290098 |  |  |  |  |  |  |  |
| B08 | WGS (Illumina) | 1785532500 | 5634499599 | 5226767 | 1078 | 5226767 | 35 | 99.2 (S:95.2, D:4, F: 0.8) | 3 |
|  | WGS (Nanopore) | 3848967099 |  |  |  |  |  |  |  |
| B09 | WGS (Illumina) | 1986611400 | 3068043506 | 7181186 | 427 | 7181186 | 51 | 100 (S:99.2, D:0.8) |  |
|  | WGS (Nanopore) | 1081432106 |  |  |  |  |  |  |  |
| B10 | WGS (Illumina) | 1554118500 | 3452007054 | 7085723 | 487 | 7085723 | 61 | 100 |  |
|  | WGS (Nanopore) | 1897888554 |  |  |  |  |  |  |  |
| B11 | WGS (Illumina) | 1338192000 | 3707801944 | 4332882 | 856 | 4332882 | 46 | 100 |  |
|  | WGS (Nanopore) | 2369609944 |  |  |  |  |  |  |  |
| B12 | WGS (Illumina) | 1345484700 | 4047280197 | 5176446 | 782 | 5176446 | 35 | 99.2 (S:95.2, D:4, F: 0.8) |  |
|  | WGS (Nanopore) | 2701795497 |  |  |  |  |  |  |  |
| B13 | WGS (Illumina) | 1776483000 | 2754397953 | 5255247 | 524 | 5255247 | 62 | 100 |  |
|  | WGS (Nanopore) | 977914953 |  |  |  |  |  |  |  |
| B14 | WGS (Illumina) | 1496088600 | 2356589573 | 3751757 | 628 | 3751757 | 41 | 100 (S:99.2, D:0.8) | 1 |
|  | WGS (Nanopore) | 860500973 |  |  |  |  |  |  |  |
| B15 | WGS (Illumina) | 1988909700 | 2502679311 | 6928004 | 361 | 7081257 | 61 | 100 |  |
|  | WGS (Nanopore) | 513769611 |  |  |  |  |  |  |  |
| B16 | WGS (Illumina) | 1428717900 | 5940992784 | 6045102 | 983 | 6045102 | 50 | 100 | 1 |
|  | WGS (Nanopore) | 4512274884 |  |  |  |  |  |  |  |
| B17 | WGS (Illumina) | 1508180100 | 1766842279 | 5969873 | 296 | 5969873 | 60 | 100 | 1 |
|  | WGS (Nanopore) | 258662179 |  |  |  |  |  |  |  |
| B18 | WGS (Illumina) | 1469341500 | 2963597780 | 5226250 | 567 | 5226250 | 35 | 99.2 (S:95.2, D:4, F: 0.8) | 3 |
|  | WGS (Nanopore) | 1494256280 |  |  |  |  |  |  |  |
| B19 | WGS (Illumina) | 1137226500 | 2376352624 | 5226646 | 455 | 5226646 | 35 | 99.2 (S:95.2, D:4, F: 0.8) | 3 |
|  | WGS (Nanopore) | 1239126124 |  |  |  |  |  |  |  |
| B20 | WGS (Illumina) | 2010952500 | 4224442413 | 5974169 | 707 | 5974169 | 60 | 100 | 1 |
|  | WGS (Nanopore) | 2213489913 |  |  |  |  |  |  |  |
| B22 | WGS (Illumina) | 1604466900 | 3243298251 | 4100771 | 791 | 4100771 | 55 | 99.2 (S:99.2, F:0.8) | 3 |
|  | WGS (Nanopore) | 1638831351 |  |  |  |  |  |  |  |
| B23 | WGS (Illumina) | 1805075400 | 4698810520 | 6839629 | 687 | 6839629 | 61 | 100 |  |
|  | WGS (Nanopore) | 2893735120 |  |  |  |  |  |  |  |
| B24 | WGS (Illumina) | 1796503800 | 2553172950 | 8157816 | 313 | 8019266 | 72 | 100 (S:99.2, D:0.8) | 1 |
|  | WGS (Nanopore) | 756669150 |  |  |  |  |  |  |  |
| B26 | WGS (Illumina) | 1772991900 | 2354539013 | 6020760 | 391 | 6020760 | 61 | 100 |  |
|  | WGS (Nanopore) | 581547113 |  |  |  |  |  |  |  |
| B27 | WGS (Illumina) | 1959642600 | 4674373461 | 5979496 | 782 | 5979496 | 60 | 100 | 1 |
|  | WGS (Nanopore) | 2714730861 |  |  |  |  |  |  |  |
| F02 | WGS (Illumina) | 1519977900 | 2537473027 | 3763092 | 674 | 3763092 | 65 | 100 (S:99.2, D:0.8) |  |
|  | WGS (Nanopore) | 1017495127 |  |  |  |  |  |  |  |

Supplementary Table S7. *Top 20 KEGG Pathway Hits for Bacterial Isolates.*

|  | ***Pseudomonas*** | | | | | | | ***Bacillus*** | | | | | ***Plantibacter*** | ***Brevibacillus*** | ***Pantoea*** | ***Streptomyces*** | ***Arthrobacter*** |
| --- | --- | --- | --- | --- | --- | --- | --- | --- | --- | --- | --- | --- | --- | --- | --- | --- | --- |
| **Pathway** | **B03** | **B05** | **B06** | **B10** | **B20** | **B23** | **B26** | **B04** | **B08** | **B09** | **B12** | **B14** | **B07** | **B16** | **B22** | **B24** | **F2** |
| Metabolic pathways |  |  |  |  |  |  |  |  |  |  |  |  |  |  |  |  |  |
| Biosynthesis of secondary metabolites |  |  |  |  |  |  |  |  |  |  |  |  |  |  |  |  |  |
| Microbial metabolism in diverse environments |  |  |  |  |  |  |  |  |  |  |  |  |  |  |  |  |  |
| ABC transporters |  |  |  |  |  |  |  |  |  |  |  |  |  |  |  |  |  |
| Two-component system |  |  |  |  |  |  |  |  |  |  |  |  |  |  |  |  |  |
| Biosynthesis of cofactors |  |  |  |  |  |  |  |  |  |  |  |  |  |  |  |  |  |
| Biosynthesis of amino acids |  |  |  |  |  |  |  |  |  |  |  |  |  |  |  |  |  |
| Carbon metabolism |  |  |  |  |  |  |  |  |  |  |  |  |  |  |  |  |  |
| Purine metabolism |  |  |  |  |  |  |  |  |  |  |  |  |  |  |  |  |  |
| Biofilm formation - Pseudomonas aeruginosa |  |  |  |  |  |  |  |  |  |  |  |  |  |  |  |  |  |
| Ribosome |  |  |  |  |  |  |  |  |  |  |  |  |  |  |  |  |  |
| Oxidative phosphorylation |  |  |  |  |  |  |  |  |  |  |  |  |  |  |  |  |  |
| Glycine, serine and threonine metabolism |  |  |  |  |  |  |  |  |  |  |  |  |  |  |  |  |  |
| Nucleotide metabolism |  |  |  |  |  |  |  |  |  |  |  |  |  |  |  |  |  |
| Flagellar assembly |  |  |  |  |  |  |  |  |  |  |  |  |  |  |  |  |  |
| Porphyrin metabolism |  |  |  |  |  |  |  |  |  |  |  |  |  |  |  |  |  |
| Quorum sensing |  |  |  |  |  |  |  |  |  |  |  |  |  |  |  |  |  |
| Biosynthesis of nucleotide sugars |  |  |  |  |  |  |  |  |  |  |  |  |  |  |  |  |  |
| Pyruvate metabolism |  |  |  |  |  |  |  |  |  |  |  |  |  |  |  |  |  |
| Glyoxylate and dicarboxylate metabolism |  |  |  |  |  |  |  |  |  |  |  |  |  |  |  |  |  |

Supplementary Table S8: Biosynthetic Gene Clusters (BGCs) in Bacteria with Similarity to Known Clusters Identified using antiSMASH

| **Isolate** | **Region** | **Type** | **From** | **To** | **Most similar known cluster** | **Similarity** |
| --- | --- | --- | --- | --- | --- | --- |
| B03 | 1 | lanthipeptide-class-ii | 1 | 18345 |  |  |
| B03 | 2 | NRPS-like | 422827 | 466369 | fragin | 0.37 |
| B03 | 3 | arylpolyene | 753667 | 797278 | APE Vf | 0.4 |
| B03 | 4 | RiPP-like | 1634576 | 1645445 |  |  |
| B03 | 5 | NAGGN | 2036207 | 2051102 |  |  |
| B03 | 6 | NRPS | 2179369 | 2232373 | Pf-5 pyoverdine | 0.1 |
| B03 | 7 | hydrogen-cyanide | 2752907 | 2765888 | hydrogen cyanide | 1 |
| B03 | 8 | ranthipeptide | 2798592 | 2820022 | Pf-5 pyoverdine | 0.08 |
| B03 | 9 | NRPS-like,T1PKS,NRP-metallophore,NRPS | 2947101 | 3054409 | histicorrugatin | 0.84 |
| B03 | 10 | RiPP-like | 3178790 | 3189635 |  |  |
| B03 | 11 | NRP-metallophore,NRPS | 3507563 | 3571337 | enantio-pyochelin | 0.4 |
| B03 | 12 | hserlactone,NRPS | 3816809 | 3983964 | syringomycin | 1 |
| B03 | 13 | hydrogen-cyanide | 4001862 | 4014671 |  |  |
| B03 | 14 | butyrolactone | 4196612 | 4210022 | grimoviridin | 0.18 |
| B03 | 15 | betalactone | 4484229 | 4507479 | fengycin | 0.13 |
| B03 | 16 | hydrogen-cyanide | 4508333 | 4521257 |  |  |
| B03 | 17 | NRP-metallophore,NRPS | 4653681 | 4752406 | pyoverdine SMX-1 | 0.58 |
| B03 | 18 | redox-cofactor | 6156914 | 6179061 | lankacidin C | 0.13 |
| B04 | 1 | NRPS | 362089 | 427538 | lichenysin | 1 |
| B04 | 2 | thiopeptide,LAP | 1035524 | 1076814 | butirosin A/butirosin B | 0.07 |
| B04 | 3 | NI-siderophore | 1183953 | 1217419 | schizokinen | 0.6 |
| B04 | 4 | betalactone | 2048774 | 2077288 | fengycin | 0.53 |
| B04 | 5 | terpene | 2162026 | 2183924 |  |  |
| B04 | 6 | T3PKS | 2376059 | 2417156 |  |  |
| B04 | 7 | CDPS | 3530202 | 3550950 | pulcherriminic acid | 0.5 |
| B04 | 8 | NRP-metallophore,NRPS | 3828482 | 3880235 | bacillibactin/bacillibactin E/bacillibactin F | 1 |
| B04 | 9 | lassopeptide,RRE-containing | 3979833 | 4002566 |  |  |
| B04 | 10 | lanthipeptide-class-ii | 4066448 | 4093409 | lichenicidin VK21 A1/lichenicidin VK21 A2 | 1 |
| B05 | 1 | redox-cofactor | 430692 | 452851 | lankacidin C | 0.13 |
| B05 | 2 | ectoine | 549724 | 560161 |  |  |
| B05 | 3 | NAGGN | 1345758 | 1360543 |  |  |
| B05 | 4 | RiPP-like | 2154546 | 2166741 |  |  |
| B05 | 5 | terpene | 2840892 | 2864468 | carotenoid | 1 |
| B05 | 6 | NRP-metallophore,RiPP-like,NRPS | 3205488 | 3279776 | pyoverdine SMX-1 | 0.22 |
| B05 | 7 | NRP-metallophore,NRPS | 3491619 | 3547140 | Pf-5 pyoverdine | 0.08 |
| B05 | 8 | arylpolyene | 4933633 | 4977240 | APE Vf | 0.4 |
| B06 | 1.1 | redox-cofactor | 442184 | 464343 | lankacidin C | 0.13 |
| B06 | 1.2 | NAGGN | 1351529 | 1366314 |  |  |
| B06 | 1.3 | RiPP-like | 2094019 | 2106214 |  |  |
| B06 | 1.4 | terpene | 2699664 | 2723240 | carotenoid | 1 |
| B06 | 1.5 | NRP-metallophore,RiPP-like,NRPS | 3072649 | 3147065 | pyoverdine SMX-1 | 0.22 |
| B06 | 1.6 | NRP-metallophore,NRPS | 3360372 | 3415071 | Pf-5 pyoverdine | 0.06 |
| B06 | 1.7 | arylpolyene | 4854506 | 4898113 | APE Vf | 0.4 |
| B06 | 2.1 | NAGGN | 78620 | 93405 |  |  |
| B07 | 1 | hydrogen-cyanide | 116712 | 129772 | aborycin | 0.21 |
| B07 | 2 | T3PKS | 831075 | 872202 |  |  |
| B07 | 3 | betalactone | 1284432 | 1310304 | microansamycin | 0.07 |
| B07 | 4 | NAPAA | 1617984 | 1651988 | ε-Poly-L-lysine | 1 |
| B07 | 5 | T3PKS | 1777996 | 1819189 | 5-acetyl-5,10-dihydrophenazine-1-carboxylic acid/5-(2-hydroxyacetyl)-5,10-dihydrophenazine-1-carboxylic acid/endophenazine A1/endophenazine F/endophenazine G | 0.17 |
| B07 | 6 | RiPP-like | 2838423 | 2849226 |  |  |
| B07 | 7 | lanthipeptide-class-iii | 3038476 | 3061082 | dechlorocuracomycin | 0.08 |
| B07 | 8 | T1PKS | 3226378 | 3271081 | meilingmycin | 0.02 |
| B07 | 9 | terpene | 3772320 | 3793288 | carotenoid | 0.21 |
| B08 | 1.1 | LAP | 1219089 | 1242596 |  |  |
| B08 | 1.2 | NI-siderophore | 1852416 | 1884122 | petrobactin | 1 |
| B08 | 1.3 | NRP-metallophore,NRPS | 2215323 | 2267055 | bacillibactin | 0.85 |
| B08 | 1.4 | betalactone | 2396812 | 2422050 | fengycin | 0.4 |
| B08 | 1.5 | RiPP-like | 2521628 | 2531894 |  |  |
| B08 | 1.6 | NRPS | 2545793 | 2592803 |  |  |
| B08 | 1.7 | terpene | 3372158 | 3394011 | molybdenum cofactor | 0.17 |
| B08 | 1.8 | lassopeptide,RRE-containing | 4698084 | 4720586 |  |  |
| B08 | 4.1 | RRE-containing | 15296 | 33987 |  |  |
| B09 | 1 | proteusin | 1653652 | 1673891 |  |  |
| B09 | 2 | cyclic-lactone-autoinducer | 2173148 | 2193814 |  |  |
| B09 | 3 | NRPS-like | 2986677 | 3030402 |  |  |
| B09 | 4 | T3PKS | 3093456 | 3134613 |  |  |
| B09 | 5 | opine-like-metallophore | 3211892 | 3234012 | bacillopaline | 1 |
| B09 | 6 | terpene | 3831895 | 3852680 |  |  |
| B09 | 7 | NRP-metallophore,NRPS | 4980316 | 5032328 | bacillibactin | 1 |
| B09 | 8 | RRE-containing | 5963275 | 5983556 | O&K-antigen | 0.04 |
| B09 | 9 | ectoine | 6291650 | 6302042 | ectoine | 1 |
| B10 | 1 | RiPP-like | 162724 | 173569 |  |  |
| B10 | 2 | NRPS-like | 395639 | 439007 | ambactin | 0.25 |
| B10 | 3 | arylpolyene | 744711 | 788286 | APE Vf | 0.4 |
| B10 | 4 | NRPS | 2206286 | 2259182 | Pf-5 pyoverdine | 0.09 |
| B10 | 5 | NAGGN | 2294775 | 2309556 |  |  |
| B10 | 6 | betalactone | 2813686 | 2836870 | fengycin | 0.13 |
| B10 | 7 | methanobactin | 3444216 | 3465603 | methanobactin | 0.66 |
| B10 | 8 | NRP-metallophore,NRPS | 3668762 | 3722347 | pyochelin | 0.92 |
| B10 | 9 | RiPP-like | 4003068 | 4015266 | lipopolysaccharide | 0.05 |
| B10 | 10 | NRP-metallophore,NRPS | 4218450 | 4292675 | Pf-5 pyoverdine | 0.07 |
| B10 | 11 | hydrogen-cyanide | 4296668 | 4309392 |  |  |
| B10 | 12 | RiPP-like | 5636584 | 5647462 |  |  |
| B10 | 13 | redox-cofactor | 6708610 | 6730757 | lankacidin C | 0.13 |
| B11 | 1 | NRPS | 362089 | 427538 | lichenysin | 1 |
| B11 | 2 | thiopeptide,LAP | 1035524 | 1076814 | butirosin A/butirosin B | 0.07 |
| B11 | 3 | NI-siderophore | 1183953 | 1217419 | schizokinen | 0.6 |
| B11 | 4 | betalactone | 2048774 | 2077288 | fengycin | 0.53 |
| B11 | 5 | terpene | 2162026 | 2183924 |  |  |
| B11 | 6 | T3PKS | 2376059 | 2417156 |  |  |
| B11 | 7 | CDPS | 3530202 | 3550950 | pulcherriminic acid | 0.5 |
| B11 | 8 | NRP-metallophore,NRPS | 3828482 | 3880235 | bacillibactin/bacillibactin E/bacillibactin F | 1 |
| B11 | 9 | lassopeptide,RRE-containing | 3979833 | 4002566 |  |  |
| B11 | 10 | lanthipeptide-class-ii | 4066448 | 4093409 | lichenicidin VK21 A1/lichenicidin VK21 A2 | 1 |
| B12 | 1 | LAP | 1201406 | 1224913 |  |  |
| B12 | 2 | NI-siderophore | 1860210 | 1891915 | petrobactin | 1 |
| B12 | 3 | NRP-metallophore,NRPS | 2189035 | 2240774 | bacillibactin | 0.85 |
| B12 | 4 | betalactone | 2340886 | 2366124 | fengycin | 0.4 |
| B12 | 5 | RiPP-like | 2408233 | 2418442 |  |  |
| B12 | 6 | RiPP-like | 2465012 | 2475278 |  |  |
| B12 | 7 | terpene | 3288916 | 3310769 | molybdenum cofactor | 0.17 |
| B12 | 8 | lassopeptide | 3435941 | 3459866 | paeninodin | 1 |
| B13 | 1 | redox-cofactor | 442184 | 464343 | lankacidin C | 0.13 |
| B13 | 2 | NAGGN | 1351528 | 1366313 |  |  |
| B13 | 3 | RiPP-like | 2094018 | 2106213 |  |  |
| B13 | 4 | terpene | 2699650 | 2723226 | carotenoid | 1 |
| B13 | 5 | NRP-metallophore,RiPP-like,NRPS | 3072635 | 3147051 | pyoverdine SMX-1 | 0.22 |
| B13 | 6 | NRP-metallophore,NRPS | 3360358 | 3415057 | Pf-5 pyoverdine | 0.06 |
| B13 | 7 | arylpolyene | 4854492 | 4898099 | APE Vf | 0.4 |
| B14 | 1 | NRPS | 351991 | 435716 | lichenysin | 0.85 |
| B14 | 2 | RRE-containing | 875418 | 896323 |  |  |
| B14 | 3 | terpene,NI-siderophore | 1045300 | 1082897 | schizokinen | 0.6 |
| B14 | 4 | betalactone | 1825351 | 1853762 | fengycin | 0.53 |
| B14 | 5 | terpene | 1914929 | 1936806 |  |  |
| B14 | 6 | T3PKS | 1975217 | 2016314 | laterocidine | 0.05 |
| B14 | 7 | RiPP-like | 2325016 | 2335342 |  |  |
| B14 | 8 | betalactone | 2472914 | 2505331 | bottromycin A2 | 0.06 |
| B14 | 9 | other | 3415470 | 3456891 | bacilysin | 0.85 |
| B15 | 1.1 | betalactone | 371120 | 394304 | fengycin | 0.13 |
| B15 | 1.2 | methanobactin | 1001070 | 1022457 | methanobactin | 0.66 |
| B15 | 1.3 | NRP-metallophore,NRPS | 1225616 | 1279201 | pyochelin | 0.92 |
| B15 | 1.4 | RiPP-like | 1559922 | 1572120 | lipopolysaccharide | 0.05 |
| B15 | 1.5 | NRP-metallophore,NRPS | 1775304 | 1849529 | Pf-5 pyoverdine | 0.07 |
| B15 | 1.6 | hydrogen-cyanide | 1853522 | 1866246 |  |  |
| B15 | 1.7 | RiPP-like | 3193429 | 3204307 |  |  |
| B15 | 1.8 | redox-cofactor | 4265559 | 4287706 | lankacidin C | 0.13 |
| B15 | 1.9 | RiPP-like | 4801625 | 4812470 |  |  |
| B15 | 1.10 | NAGGN | 5122104 | 5136885 |  |  |
| B15 | 1.11 | NRPS | 5172478 | 5225374 | Pf-5 pyoverdine | 0.09 |
| B15 | 1.12 | arylpolyene | 6643524 | 6687099 | APE Vf | 0.4 |
| B15 | 3.1 | NRPS-like | 1 | 26104 | fragin | 0.25 |
| B16 | 1.1 | NRPS | 2368790 | 2412161 | molybdenum cofactor | 0.11 |
| B16 | 1.2 | T3PKS | 3633394 | 3674461 |  |  |
| B16 | 1.3 | terpene | 3813590 | 3835500 |  |  |
| B16 | 1.4 | RiPP-like | 4199327 | 4210175 |  |  |
| B16 | 1.5 | NI-siderophore | 4340285 | 4375051 | surfactin | 0.08 |
| B16 | 1.6 | ectoine | 4662727 | 4673110 | ectoine | 0.75 |
| B16 | 1.7 | LAP | 5293511 | 5317045 | tauramamide | 0.13 |
| B16 | 1.8 | cyclic-lactone-autoinducer | 5378794 | 5399393 |  |  |
| B17 | 1.1 | RiPP-like | 157642 | 168487 |  |  |
| B17 | 1.2 | NRPS-like | 312578 | 355946 | ambactin | 0.25 |
| B17 | 1.3 | arylpolyene | 697109 | 740684 | APE Vf | 0.4 |
| B17 | 1.4 | T1PKS,NRPS | 1425690 | 1479379 | alginate | 0.91 |
| B17 | 1.5 | RiPP-like | 1829841 | 1840719 |  |  |
| B17 | 1.6 | NRPS | 2650491 | 2716368 | pyoverdine SMX-1 | 0.22 |
| B17 | 1.7 | RiPP-like | 2967809 | 2980007 | lipopolysaccharide | 0.05 |
| B17 | 1.8 | terpene | 3101807 | 3124026 | MA026 | 0.03 |
| B17 | 1.9 | hserlactone,phenazine | 3167746 | 3190506 | endophenazine A/endophenazine B | 0.38 |
| B17 | 1.10 | NRP-metallophore,NRPS | 3567343 | 3644820 | viscosin | 0.56 |
| B17 | 1.11 | betalactone | 3910006 | 3933084 | fengycin | 0.13 |
| B17 | 1.12 | NRP-metallophore,NRPS | 3993380 | 4061412 | pyoverdine SMX-1 | 0.16 |
| B17 | 1.13 | NAGGN | 4263390 | 4278282 |  |  |
| B17 | 1.14 | NRPS | 4313688 | 4366599 | Pf-5 pyoverdine | 0.09 |
| B17 | 1.15 | NI-siderophore | 4437649 | 4467574 | mevalagmapeptide A/mevalagmapeptide B/mevalagmapeptide C/mevalagmapeptide D | 0.04 |
| B17 | 1.16 | thioamitides | 4823997 | 4845953 |  |  |
| B17 | 1.17 | NRPS-like,betalactone | 4884003 | 4927479 | pyoverdine SMX-1 | 0.12 |
| B17 | 1.18 | redox-cofactor | 5618235 | 5640382 |  |  |
| B18 | 1.1 | LAP | 1219079 | 1242586 |  |  |
| B18 | 1.2 | NI-siderophore | 1852244 | 1883950 | petrobactin | 1 |
| B18 | 1.3 | NRP-metallophore,NRPS | 2215151 | 2266883 | bacillibactin | 0.85 |
| B18 | 1.4 | betalactone | 2396640 | 2421878 | fengycin | 0.4 |
| B18 | 1.5 | RiPP-like | 2521456 | 2531722 |  |  |
| B18 | 1.6 | NRPS | 2545621 | 2592631 |  |  |
| B18 | 1.7 | terpene | 3371986 | 3393839 | molybdenum cofactor | 0.17 |
| B18 | 1.8 | lassopeptide,RRE-containing | 4697912 | 4720414 |  |  |
| B18 | 4.1 | RRE-containing | 15304 | 33995 |  |  |
| B19 | 1.1 | LAP | 1219085 | 1242592 |  |  |
| B19 | 1.2 | NI-siderophore | 1852430 | 1884136 | petrobactin | 1 |
| B19 | 1.3 | NRP-metallophore,NRPS | 2215337 | 2267069 | bacillibactin | 0.85 |
| B19 | 1.4 | betalactone | 2396826 | 2422064 | fengycin | 0.4 |
| B19 | 1.5 | RiPP-like | 2521642 | 2531908 |  |  |
| B19 | 1.6 | NRPS | 2545807 | 2592817 |  |  |
| B19 | 1.7 | terpene | 3372160 | 3394013 | molybdenum cofactor | 0.17 |
| B19 | 1.8 | lassopeptide,RRE-containing | 4698086 | 4720588 |  |  |
| B19 | 4.1 | RRE-containing | 19289 | 33995 |  |  |
| B20 | 1.1 | RiPP-like | 157654 | 168499 |  |  |
| B20 | 1.2 | NRPS-like | 312601 | 355969 | ambactin | 0.25 |
| B20 | 1.3 | arylpolyene | 697132 | 740707 | APE Vf | 0.4 |
| B20 | 1.4 | T1PKS,NRPS | 1426577 | 1480266 | alginate | 0.91 |
| B20 | 1.5 | RiPP-like | 1830730 | 1841608 |  |  |
| B20 | 1.6 | NRPS | 2651338 | 2717215 | pyoverdine SMX-1 | 0.22 |
| B20 | 1.7 | RiPP-like | 2968914 | 2981112 | lipopolysaccharide | 0.05 |
| B20 | 1.8 | terpene | 3102913 | 3125132 | MA026 | 0.03 |
| B20 | 1.9 | hserlactone,phenazine | 3168852 | 3191612 | endophenazine A/endophenazine B | 0.38 |
| B20 | 1.10 | NRP-metallophore,NRPS | 3568474 | 3645951 | viscosin | 0.56 |
| B20 | 1.11 | betalactone | 3914584 | 3937662 | fengycin | 0.13 |
| B20 | 1.12 | NRP-metallophore,NRPS | 3997965 | 4065997 | pyoverdine SMX-1 | 0.16 |
| B20 | 1.13 | NAGGN | 4268065 | 4282957 |  |  |
| B20 | 1.14 | NRPS | 4318363 | 4371274 | Pf-5 pyoverdine | 0.09 |
| B20 | 1.15 | NI-siderophore | 4442324 | 4472249 | mevalagmapeptide A/mevalagmapeptide B/mevalagmapeptide C/mevalagmapeptide D | 0.04 |
| B20 | 1.16 | thioamitides | 4828672 | 4850628 |  |  |
| B20 | 1.17 | NRPS-like,betalactone | 4888678 | 4932154 | pyoverdine SMX-1 | 0.12 |
| B20 | 1.18 | redox-cofactor | 5622531 | 5644678 |  |  |
| B22 | 1.1 | redox-cofactor | 2155346 | 2177512 | lankacidin C | 0.13 |
| B22 | 1.2 | arylpolyene,hserlactone | 2626873 | 2686711 | aryl polyenes | 0.94 |
| B22 | 1.3 | thiopeptide | 2752659 | 2778915 | O-antigen | 0.14 |
| B22 | 1.4 | hserlactone | 3572538 | 3593176 |  |  |
| B22 | 1.5 | NRP-metallophore,NRPS | 3668758 | 3722449 | frederiksenibactin | 0.84 |
| B22 | 2.1 | NI-siderophore | 170299 | 200653 | desferrioxamine E | 1 |
| B22 | 2.2 | terpene | 381734 | 405295 | carotenoid | 1 |
| B23 | 1 | NRPS-like | 3074 | 46451 | pyoverdine DC3000 | 0.07 |
| B23 | 2 | arylpolyene | 313825 | 357400 | APE Vf | 0.45 |
| B23 | 3 | NRPS-like | 669184 | 712561 | pyoverdine DC3000 | 0.07 |
| B23 | 4 | RiPP-like | 898180 | 909025 |  |  |
| B23 | 5 | NRPS-like,betalactone | 1046426 | 1094028 | kutzneride 2 | 0.13 |
| B23 | 6 | NRPS-like | 1104577 | 1146355 | coronatine | 0.87 |
| B23 | 7 | redox-cofactor | 1417496 | 1439643 | lankacidin C | 0.13 |
| B23 | 8 | arylpolyene | 2520640 | 2561902 | fulvuthiacene A/fulvuthiacene B | 0.08 |
| B23 | 9 | NRPS | 2837796 | 2890692 | Pf-5 pyoverdine | 0.09 |
| B23 | 10 | NAGGN | 2923357 | 2938103 |  |  |
| B23 | 11 | NRPS,NRP-metallophore | 3480221 | 3569782 | Pf-5 pyoverdine | 0.09 |
| B23 | 12 | RiPP-like | 3847454 | 3859652 |  |  |
| B23 | 13 | NRP-metallophore,NRPS | 3976579 | 4030159 | pyochelin | 0.92 |
| B23 | 14 | terpene | 4348826 | 4371051 | MA026 | 0.05 |
| B23 | 15 | betalactone | 4917556 | 4945895 | fengycin | 0.13 |
| B23 | 16 | methanobactin,betalactone | 5120260 | 5157064 | methanobactin | 0.66 |
| B23 | 17 | RiPP-like | 5950844 | 5961722 |  |  |
| B24 | 1.1 | lanthipeptide-class-i | 4137 | 30593 |  |  |
| B24 | 1.2 | terpene,T1PKS,NRPS-like,hglE-KS,lanthipeptide-class-i | 213343 | 430857 | candicidin | 0.95 |
| B24 | 1.3 | indole | 582697 | 603812 | 5-dimethylallylindole-3-acetonitrile | 1 |
| B24 | 1.4 | terpene | 668447 | 692662 | carotenoid | 0.54 |
| B24 | 1.5 | NRP-metallophore,NRPS | 1236901 | 1300360 | coelibactin | 1 |
| B24 | 1.6 | ectoine | 1866900 | 1877298 | ectoine | 1 |
| B24 | 1.7 | RRE-containing | 1907732 | 1927986 | citrulassin E | 1 |
| B24 | 1.8 | melanin | 2764913 | 2775479 | istamycin | 0.04 |
| B24 | 1.9 | NI-siderophore | 2850129 | 2879901 | desferrioxamin B/desferrioxamine E | 1 |
| B24 | 1.10 | lanthipeptide-class-iii,RiPP-like | 4174450 | 4197152 | cacaoidin | 0.11 |
| B24 | 1.11 | lanthipeptide-class-i | 4372338 | 4397120 | planosporicin | 1 |
| B24 | 1.12 | terpene | 5174446 | 5195537 | albaflavenone | 1 |
| B24 | 1.13 | T2PKS | 5245384 | 5317932 | spore pigment | 0.66 |
| B24 | 1.14 | NI-siderophore | 5803746 | 5833770 | kinamycin | 0.19 |
| B24 | 1.15 | RiPP-like | 6066011 | 6077336 |  |  |
| B24 | 1.16 | terpene | 6097233 | 6119413 | geosmin | 1 |
| B24 | 1.17 | NI-siderophore | 6245093 | 6276248 | paulomycin | 0.13 |
| B24 | 1.18 | NRPS | 6288699 | 6368692 | CDA1b/CDA2a/CDA2b/CDA3a/CDA3b/CDA4a/CDA4b | 0.75 |
| B24 | 1.19 | terpene | 6794133 | 6820918 | hopene | 1 |
| B24 | 1.20 | hydrogen-cyanide | 6933052 | 6945907 | aborycin | 0.21 |
| B24 | 1.21 | terpene | 7089002 | 7110033 | versipelostatin | 0.05 |
| B24 | 1.22 | RiPP-like | 7126371 | 7136586 | informatipeptin | 0.42 |
| B24 | 1.23 | NRP-metallophore,NRPS | 7390559 | 7449013 | coelichelin | 1 |
| B24 | 1.24 | lanthipeptide-class-iii | 7658988 | 7681570 | SapB | 1 |
| B24 | 1.25 | lanthipeptide-class-i | 7958991 | 7985447 |  |  |
| B24 | 2.1 | T2PKS | 34896 | 123733 | fluostatins M-Q | 0.67 |
| B26 | 1 | RiPP-like | 492697 | 503572 |  |  |
| B26 | 2 | arylpolyene | 1457848 | 1501423 | APE Vf | 0.4 |
| B26 | 3 | NRPS-like | 1791488 | 1834865 | ambactin | 0.25 |
| B26 | 4 | RiPP-like | 2012866 | 2023711 |  |  |
| B26 | 5 | redox-cofactor | 2520642 | 2542789 | lankacidin C | 0.13 |
| B26 | 6 | NRPS | 3784987 | 3837868 | Pf-5 pyoverdine | 0.09 |
| B26 | 7 | NAGGN | 3874692 | 3889543 |  |  |
| B26 | 8 | terpene | 3917309 | 3940960 | carotenoid | 1 |
| B26 | 9 | NI-siderophore | 4076347 | 4106152 |  |  |
| B26 | 10 | hydrogen-cyanide | 4142414 | 4155231 |  |  |
| B26 | 11 | betalactone | 4247469 | 4270879 | fengycin | 0.13 |
| B26 | 12 | NRPS | 4934171 | 5024749 | bananamide 1/bananamide 2/bananamide 3 | 1 |
| B26 | 13 | NRP-metallophore,NRPS | 5165161 | 5241870 | pyoverdine SMX-1 | 0.35 |
| B26 | 14 | RiPP-like | 5364549 | 5376747 | lipopolysaccharide | 0.05 |
| B27 | 1.1 | NRP-metallophore,NRPS | 255200 | 332677 | viscosin | 0.56 |
| B27 | 1.2 | betalactone | 598237 | 621315 | fengycin | 0.13 |
| B27 | 1.3 | NRP-metallophore,NRPS | 681617 | 749649 | pyoverdine SMX-1 | 0.16 |
| B27 | 1.4 | NAGGN | 951717 | 966609 |  |  |
| B27 | 1.5 | NRPS | 1002015 | 1054926 | Pf-5 pyoverdine | 0.09 |
| B27 | 1.6 | NI-siderophore | 1125976 | 1155901 | mevalagmapeptide A/mevalagmapeptide B/mevalagmapeptide C/mevalagmapeptide D | 0.04 |
| B27 | 1.7 | thioamitides | 1512324 | 1534280 |  |  |
| B27 | 1.8 | NRPS-like,betalactone | 1572330 | 1615806 | pyoverdine SMX-1 | 0.12 |
| B27 | 1.9 | redox-cofactor | 2306187 | 2328334 |  |  |
| B27 | 1.10 | RiPP-like | 2815478 | 2826323 |  |  |
| B27 | 1.11 | NRPS-like | 2970425 | 3013793 | ambactin | 0.25 |
| B27 | 1.12 | arylpolyene | 3354956 | 3398531 | APE Vf | 0.4 |
| B27 | 1.13 | T1PKS,NRPS | 4084420 | 4138109 | alginate | 0.91 |
| B27 | 1.14 | RiPP-like | 4488573 | 4499451 |  |  |
| B27 | 1.15 | NRPS | 5309255 | 5375132 | pyoverdine SMX-1 | 0.22 |
| B27 | 1.16 | RiPP-like | 5626772 | 5638970 | lipopolysaccharide | 0.05 |
| B27 | 1.17 | terpene | 5761294 | 5783513 | MA026 | 0.03 |
| B27 | 1.18 | hserlactone,phenazine | 5827233 | 5849993 | endophenazine A/endophenazine B | 0.38 |
| F02 | 1 | LAP | 175036 | 196484 |  |  |
| F02 | 2 | NRPS-like | 298478 | 341051 |  |  |
| F02 | 3 | NI-siderophore | 529716 | 559572 | legonoxamine A/desferrioxamine B/legonoxamine B | 0.66 |
| F02 | 4 | NAPAA | 1475959 | 1509846 | stenothricin | 0.31 |
| F02 | 5 | betalactone | 1544899 | 1570315 | microansamycin | 0.07 |
| F02 | 6 | terpene | 1826516 | 1847427 | carotenoid | 0.66 |
| F02 | 7 | T3PKS | 1934644 | 1975858 |  |  |
| F02 | 8 | NAPAA | 2725883 | 2759800 | ε-Poly-L-lysine | 1 |

Supplementary Table S9: Unique GO Enriched Elements for Bacteria as per OrthoVenn

| **Isolate** | **GO ID** | **Namespace** | **Name** | **Count** | **p-value** |
| --- | --- | --- | --- | --- | --- |
| B03 | GO:0006355 | biological_process | regulation of transcription, DNA-templated | 2 | 3.80628E-26 |
|  | [GO:0003700](http://amigo.geneontology.org/amigo/term/GO:0003700) | molecular_function | sequence-specific DNA binding transcription factor activity | 2 | 6.72828E-05 |
|  | [GO:0009289](http://amigo.geneontology.org/amigo/term/GO:0009289) | cellular_component | pilus | 2 | 0.000277435 |
|  | [GO:0006865](http://amigo.geneontology.org/amigo/term/GO:0006865) | biological_process | amino acid transport | 2 | 0.000709542 |
|  | [GO:0006351](http://amigo.geneontology.org/amigo/term/GO:0006351) | biological_process | transcription, DNA-templated | 2 | 0.004367285 |
| B06 | [GO:0003677](http://amigo.geneontology.org/amigo/term/GO:0003677) | molecular_function | DNA binding | 2 | 0.002961218 |
| B09 | [GO:0030435](http://amigo.geneontology.org/amigo/term/GO:0030435) | biological_process | sporulation resulting in formation of a cellular spore | 2 | 1.59E-07 |
|  | [GO:0006355](http://amigo.geneontology.org/amigo/term/GO:0006355) | biological_process | regulation of transcription, DNA-templated | 2 | 0.000936305 |
|  | [GO:0009635](http://amigo.geneontology.org/amigo/term/GO:0009635) | biological_process | response to herbicide | 2 | 0.005199409 |
| B12 | [GO:0006388](http://amigo.geneontology.org/amigo/term/GO:0006388) | biological_process | tRNA splicing, via endonucleolytic cleavage and ligation | 2 | 4.52424E-05 |
|  | [GO:0043640](http://amigo.geneontology.org/amigo/term/GO:0043640) | biological_process | benzoate catabolic process via hydroxylation | 2 | 4.52424E-05 |
| B16 | [GO:0006313](http://amigo.geneontology.org/amigo/term/GO:0006313) | biological_process | transposition, DNA-mediated | 4 | 0.000735153 |
|  | [GO:0005524](http://amigo.geneontology.org/amigo/term/GO:0005524) | molecular_function | ATP binding | 5 | 0.001299941 |
|  | [GO:0015942](http://amigo.geneontology.org/amigo/term/GO:0015942) | biological_process | formate metabolic process | 2 | 0.003198059 |
| B20 | [GO:0097040](http://amigo.geneontology.org/amigo/term/GO:0002049) | biological_process | phthiocerol biosynthetic process | 2 | 0.000222432 |
|  | GO:0002049 | biological_process | pyoverdine biosynthetic process | 2 | 0.014848085 |
|  | GO:0009432 | biological_process | SOS response | 3 | 0.026076706 |
| B22 | [GO:0055114](http://amigo.geneontology.org/amigo/term/GO:0055114) | biological_process | oxidation-reduction process | 2 | 0.000454132 |
| B23 | [GO:0055085](http://amigo.geneontology.org/amigo/term/GO:0055085) | biological_process | transmembrane transport | 2 | 1.24114E-26 |
|  | [GO:0003700](http://amigo.geneontology.org/amigo/term/GO:0003700) | molecular_function | sequence-specific DNA binding transcription factor activity | 2 | 9.30E-12 |
|  | [GO:0006313](http://amigo.geneontology.org/amigo/term/GO:0006313) | biological_process | transposition, DNA-mediated | 13 | 2.6311E-05 |
|  | [GO:0050609](http://amigo.geneontology.org/amigo/term/GO:0050609) | molecular_function | phosphonate dehydrogenase activity | 2 | 0.00128958 |
|  | [GO:0015416](http://amigo.geneontology.org/amigo/term/GO:0015416) | molecular_function | organic phosphonate transmembrane-transporting ATPase activity | 2 | 0.00128958 |
|  | [GO:0032196](http://amigo.geneontology.org/amigo/term/GO:0032196) | biological_process | transposition | 8 | 0.002206643 |
|  | [GO:0006397](http://amigo.geneontology.org/amigo/term/GO:0006397) | biological_process | mRNA processing | 2 | 0.00373395 |

Supplementary Table S10: Pseudomonas orientalis genomes used in the pangenome analysis.

| **Assembly Accession** | **Strain/ Isolate** | **Assembly Level** | **WGS project accession** | **No. of Scaffolds** | **Host** | **Country** |
| --- | --- | --- | --- | --- | --- | --- |
| GCF_039673785.1 | TWI666 | Contig | JBDJMH01 |  | river water | Australia: Torrens River |
| GCF_004210185.1 | 16NI | Contig | SGFD01 |  | wheat grain | Germany: Detmold |
| GCF_004210175.1 | 133NRW | Scaffold | SGFE01 | 153 | wheat grain | Germany: Detmold |
| GCF_022807995.1 | YB-76 | Complete Genome |  | 2 | soil at 5cm depth | China: Henan Academy of Agricultural Sciences |
| GCF_001439815.1 | DSM 17489 | Contig | JYLM01 |  | spring water | Lebanon |
| GCF_900105795.1 | LMG 23660 | Chromosome |  | 1 | missing | missing |
| GCF_036857425.1 | I3 | Contig | JAXQQL01 |  | Citrus, phyllosphere | Tunisia: National Agronomic Instiute of Tunisia |
| GCF_963970785.1 | PseudoSSchier:P117 | Contig | CAXAPP01 |  | Artemisia maritima, endosphere | Netherlands |
| GCF_963970415.1 | PseudoSSchier:P114 | Contig | CAXAOR01 |  | Artemisia maritima, endosphere | Netherlands |
| GCF_013607945.1 | CDVBN20 | Scaffold | VDLW01 | 209 | Brassica napus root | Spain: Municipality in the province of Salamanca |
| GCF_002934065.1 | F9 | Complete Genome |  | 1 | apple blossom | Switzerland: Kanton Zuerich |
| GCF_003852045.1 | 8B | Complete Genome |  | 1 | Triticum sp. wheat, rhizosphere | Canada: University in Moncton |
| GCF_003851585.1 | L1-3-08 | Complete Genome |  | 1 | Triticum sp. wheat, rhizosphere | Canada: University in Moncton |
| GCF_003851605.1 | R2-66-08W | Complete Genome |  | 1 | Triticum sp. wheat, rhizosphere | Canada: University in Moncton |
| GCF_003851645.1 | R4-35-08 | Complete Genome |  | 1 | Triticum sp. wheat, rhizosphere | Canada: University in Moncton |
| GCA_029240295.1 | L_E1_T5_bin.8 | Contig | JAREAO01 |  | Agricultural soil | USA: Colorado, Fort Collins |
| GCA_041008715.1 | Phyllo_205 | Contig | JAUKOD01 |  | Pisum sativum, leaf phyllosphere | USA: New York, Ithaca |
| GCA_046628125.1 | MPB20 | Complete Genome |  | 2 | Citrullus colocytis, endosphere | UAE: Al Ain |

#### ***Supplementary Figures***

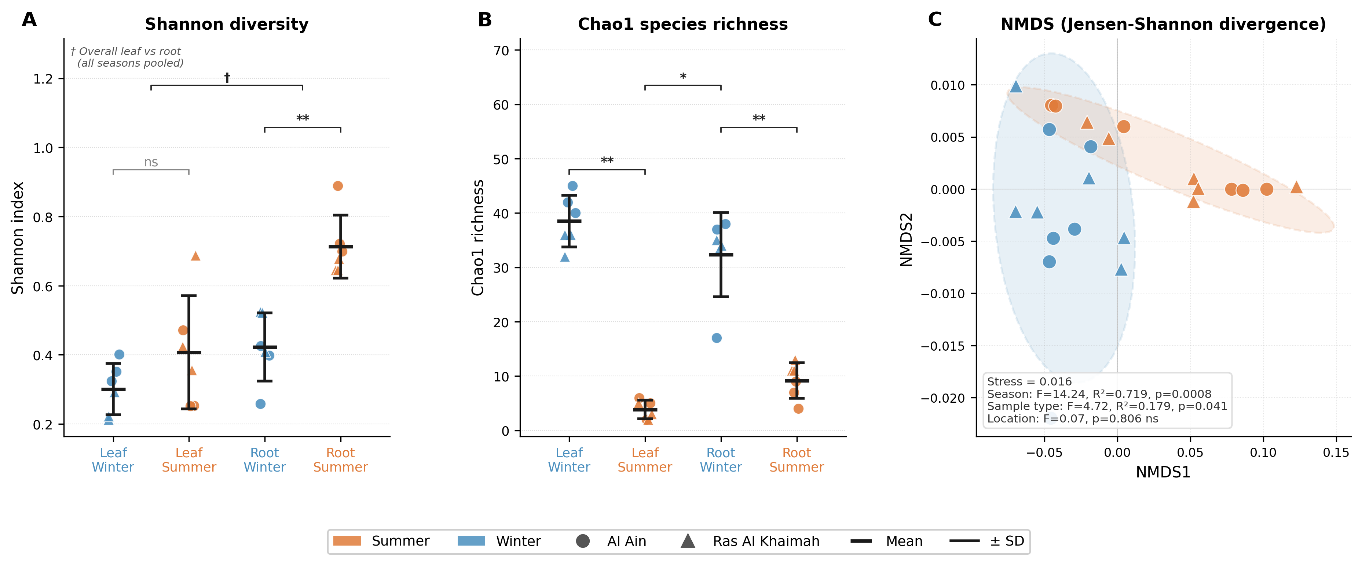

Supplementary Figure S1. Alpha and beta diversity of plant tissue-associated microbial communities. (A) Shannon diversity index and (B) Chao1 species richness of leaf- and root-associated communities across seasons and sampling locations. Individual points represent biological replicates (n = 6 per group; 3 Al Ain, 3 Ras Al Khaimah); horizontal bars and error bars indicate mean ± SD. Significance brackets denote pairwise Mann-Whitney U test results (** p < 0.01; * p < 0.05; ns, not significant). † indicates the overall comparison between leaf and root communities pooled across seasons (U = 122, p = 0.004 for Shannon; U = 3, p = 0.020 for Chao1 in summer only). (C) Non-metric multidimensional scaling (NMDS) ordination of community composition based on Jensen-Shannon divergence distances calculated at species level. Dashed ellipses represent 95% confidence intervals around seasonal group centroids. PERMANOVA results (9,999 permutations) are shown in the annotation box. Colour indicates season (orange, summer; blue, winter); marker shape indicates sampling location (circle, Al Ain; triangle, Ras Al Khaimah).

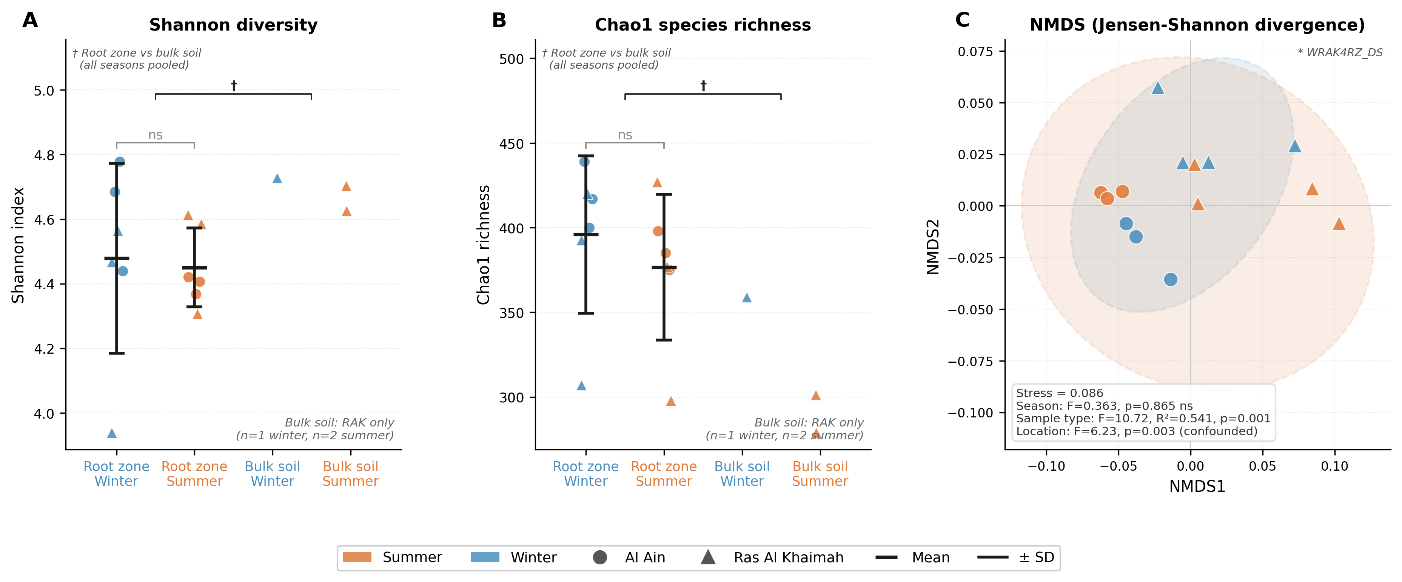

Supplementary Figure S2. Alpha and beta diversity of soil microbial communities. (A) Shannon diversity index and (B) Chao1 species richness of root zone and bulk soil communities across seasons. Root zone samples: n = 6 per season (3 Al Ain, 3 Ras Al Khaimah); bulk soil samples: n = 1 (winter) and n = 2 (summer), Ras Al Khaimah only. For root zone groups, horizontal bars and error bars indicate mean ± SD; mean and SD are not shown for bulk soil groups owing to insufficient replication. Significance brackets denote Mann-Whitney U test results (ns, not significant; * p < 0.05). † indicates the comparison between root zone and bulk soil communities pooled across seasons, which should be interpreted with caution, given the limited bulk soil sample size and single-location sampling. (C) NMDS ordination of community composition based on Jensen-Shannon divergence distances at the species level. Dashed ellipses represent 95% confidence intervals around seasonal group centroids. PERMANOVA results (9,999 permutations) are shown in the annotation box; the location result is noted as confounded with sample type, as all Al Ain samples are root zone, while Ras Al Khaimah includes both root zone and bulk soil. * denotes sample WRAK4RZ_DS, which showed an atypical ordination position and was retained in all analyses. Color indicates season (orange, summer; blue, winter); marker shape indicates sampling location (circle, Al Ain; triangle, Ras Al Khaimah).

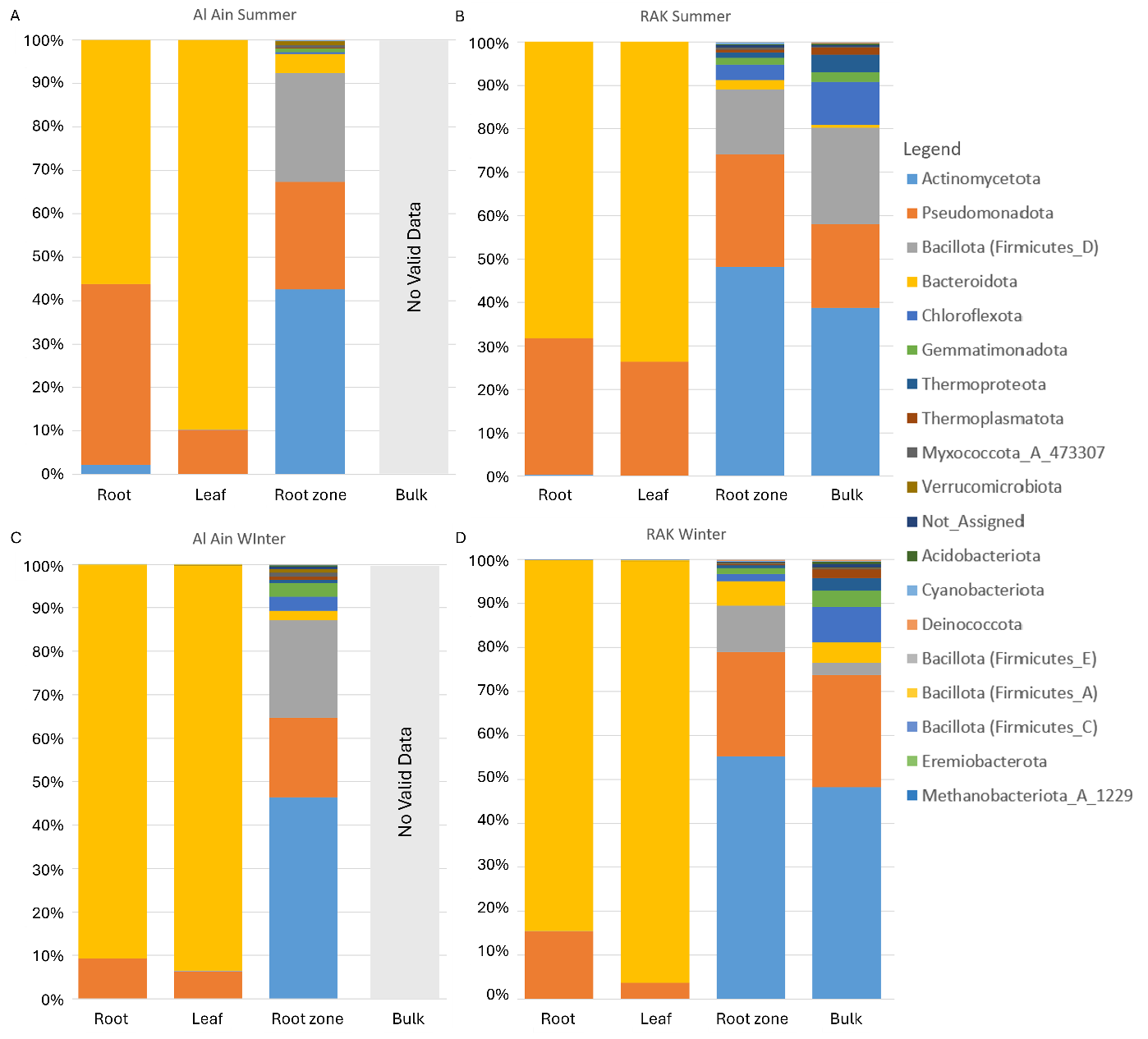

Supplementary Figure S3. Seasonal Variation in the Relative Abundance of Bacterial Phyla in Al Ain and Ras Al Khaimah (RAK) Based on 16S rDNA Amplicon Data. (A, B) Relative abundance of bacterial phyla during summer in Al Ain and RAK, respectively. (C, D) Relative abundance during winter in Al Ain and RAK, respectively. Missing bulk soil samples are labeled as “No valid data.”. To note, Firmicutes are shown as two separate taxa due to the polyphyletic lineages present within the phylum in the Greengenes database reference tree. Other datasets show the phylum as a monophyletic lineage, as per the Greengenes FAQ page (https://gtdb.ecogenomic.org/faq), and the International Code of Nomenclature of Prokaryotes (ICNP) has updated the naming conventions for prokaryotic phyla, standardizing names with the suffix "-ota" derived from the type genus (Oren & Garrity, 2021). This change had not been reflected in the Greengenes database, which still presents Bacillota as “Firmicutes” at the time of our analysis.

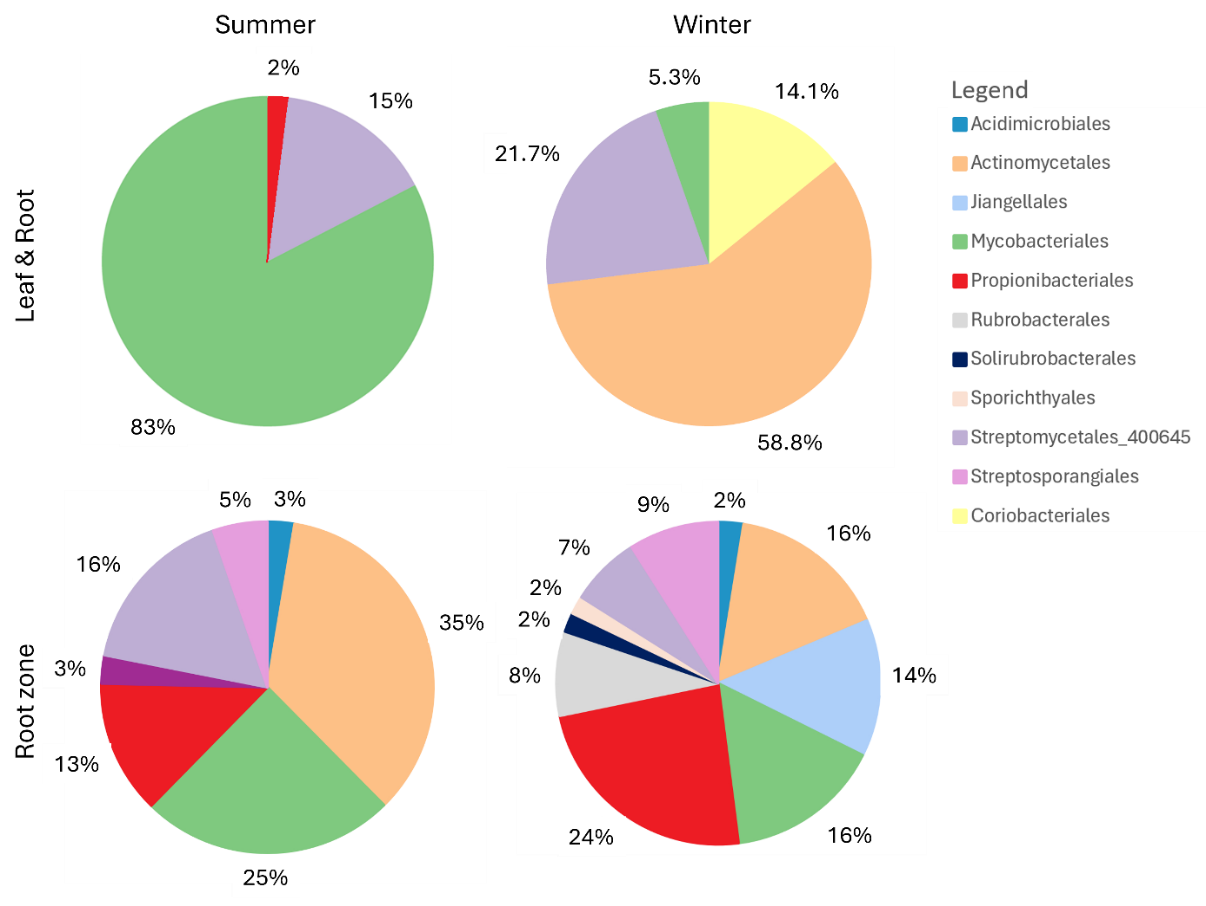

Supplementary Figure S4. Seasonal variation in the relative abundance of Actinobacteriota orders across different sample types. Pie charts show the community composition of Actinobacteriota at the order level in leaf, root, and root zone soil samples during summer and winter. The charts highlight shifts in the relative abundance of dominant and minor orders across seasons and sample types.

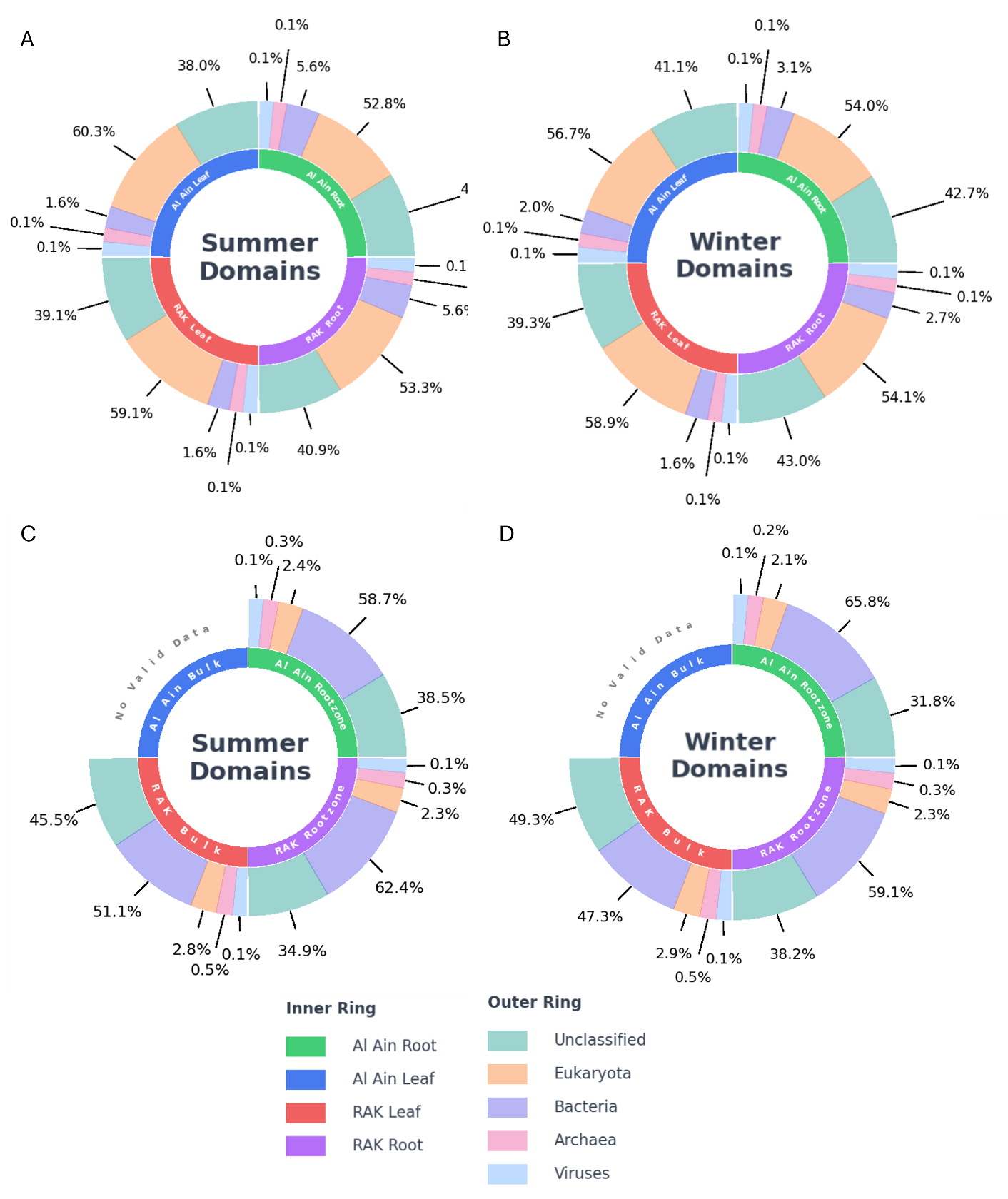

Supplementary Figure S5. Domain-level read-based metagenomic profiling of Al Ain and Ras Al Khaimah (RAK) samples across seasons. Donut charts show the proportion of sequencing reads assigned to major domains—Bacteria, Archaea, Eukaryota, Viruses, and Unclassified—in leaf and root samples during summer (A) and winter (B), and in root zone and bulk soil samples during summer (C) and winter (D). Inner rings represent individual sample types (Al Ain and RAK), while outer rings indicate the relative abundance of each domain. “No valid data” is shown for missing bulk soil samples.

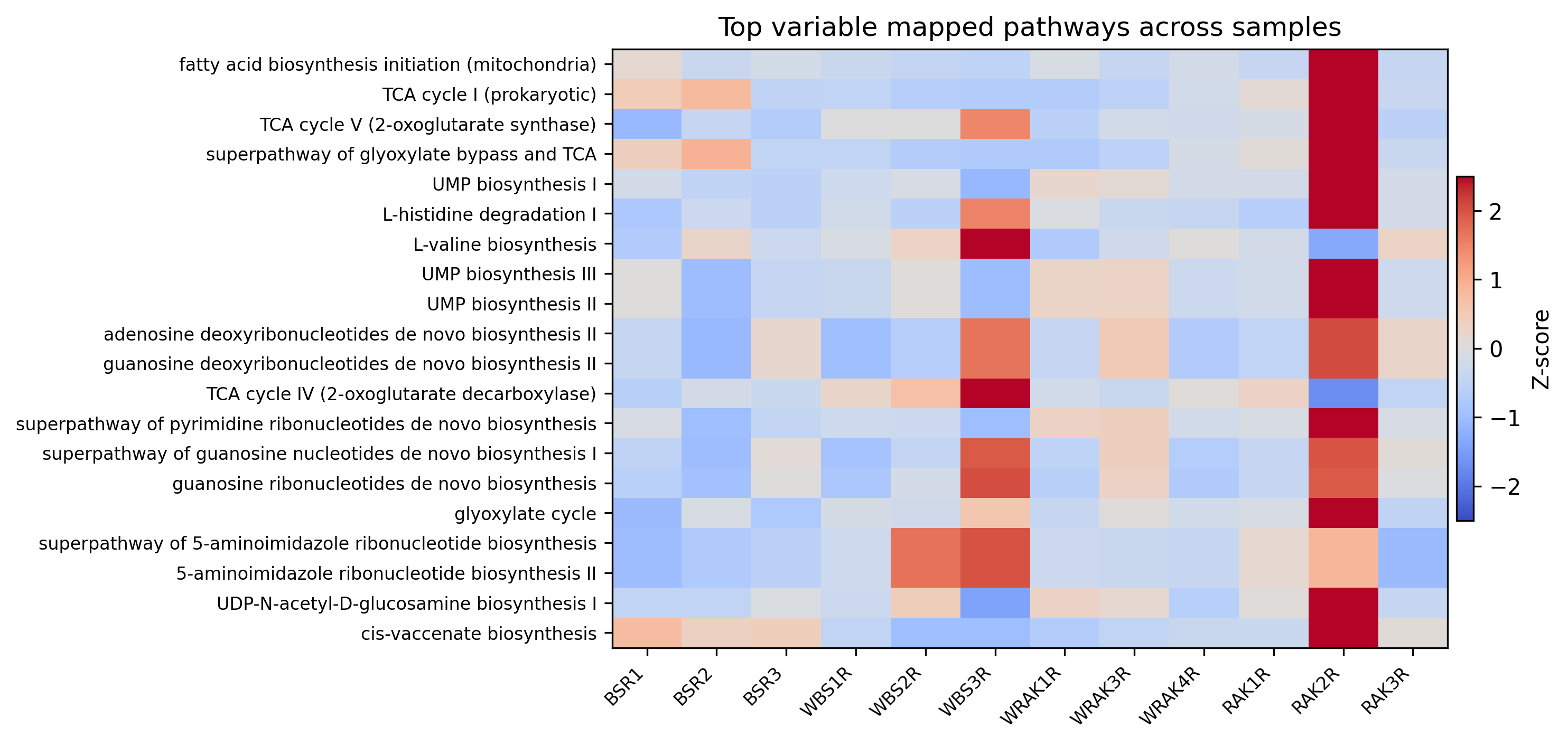

Supplementary Figure S6. Heatmap of the most variable mapped functional pathways across individual root samples, based on HUMAnN pathway profiling. Values are row-scaled to highlight relative enrichment and depletion across samples. Samples are grouped by location and season; the color scale indicates relative abundance after row scaling (warmer colors indicate enrichment, cooler colors indicate depletion).

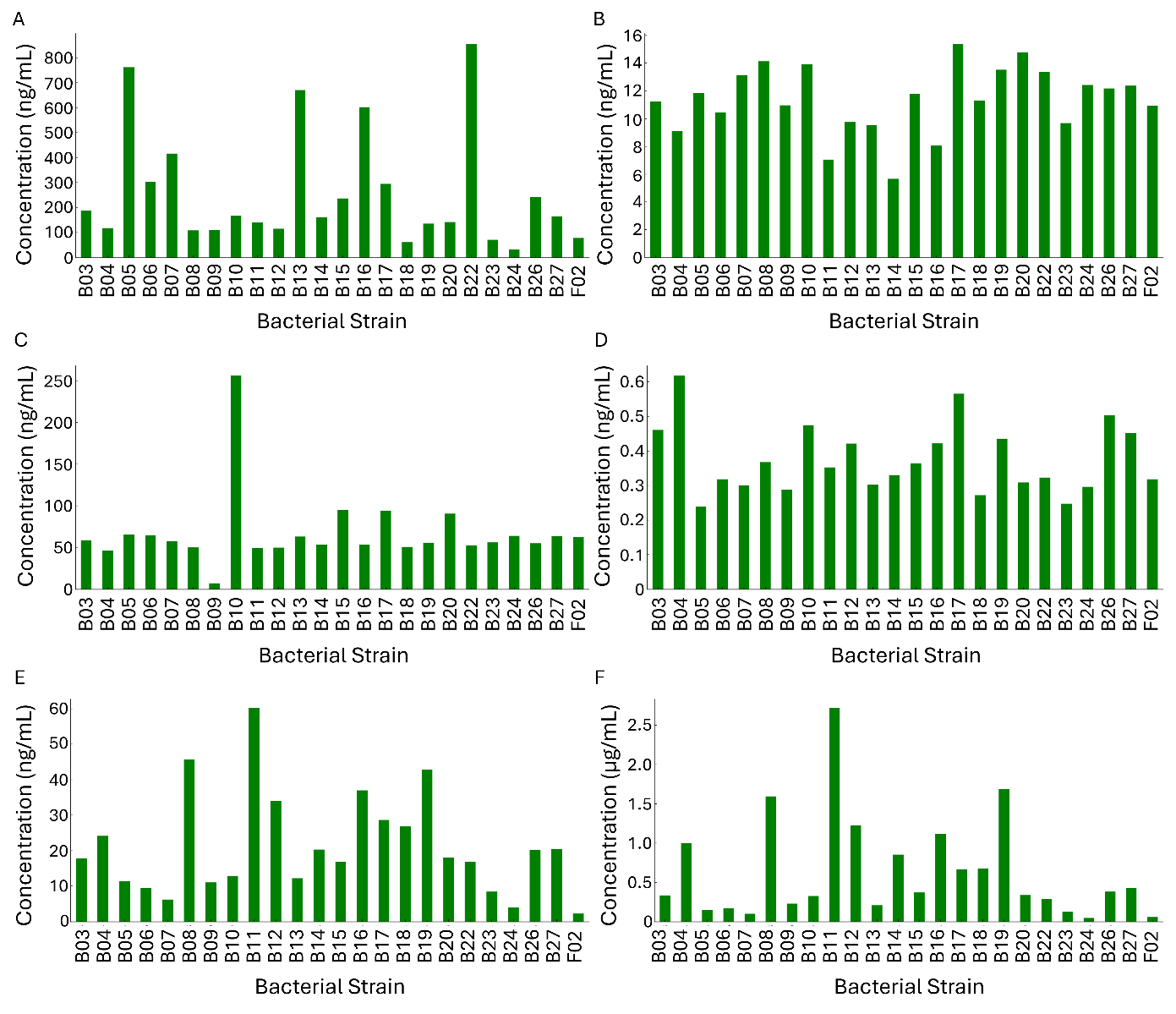

Supplementary Figure S7. Phytohormone production by bacterial isolates measured using LC-MS/MS. (A) Indole-3-acetic acid (IAA), (B) Gibberellic acid (GA_3_), (C) Salicylic acid (SA), (D) Abscisic acid (ABA), (E) Isopentyl adenine (iPA), and (F) Indole-3-butyric acid (IBA).

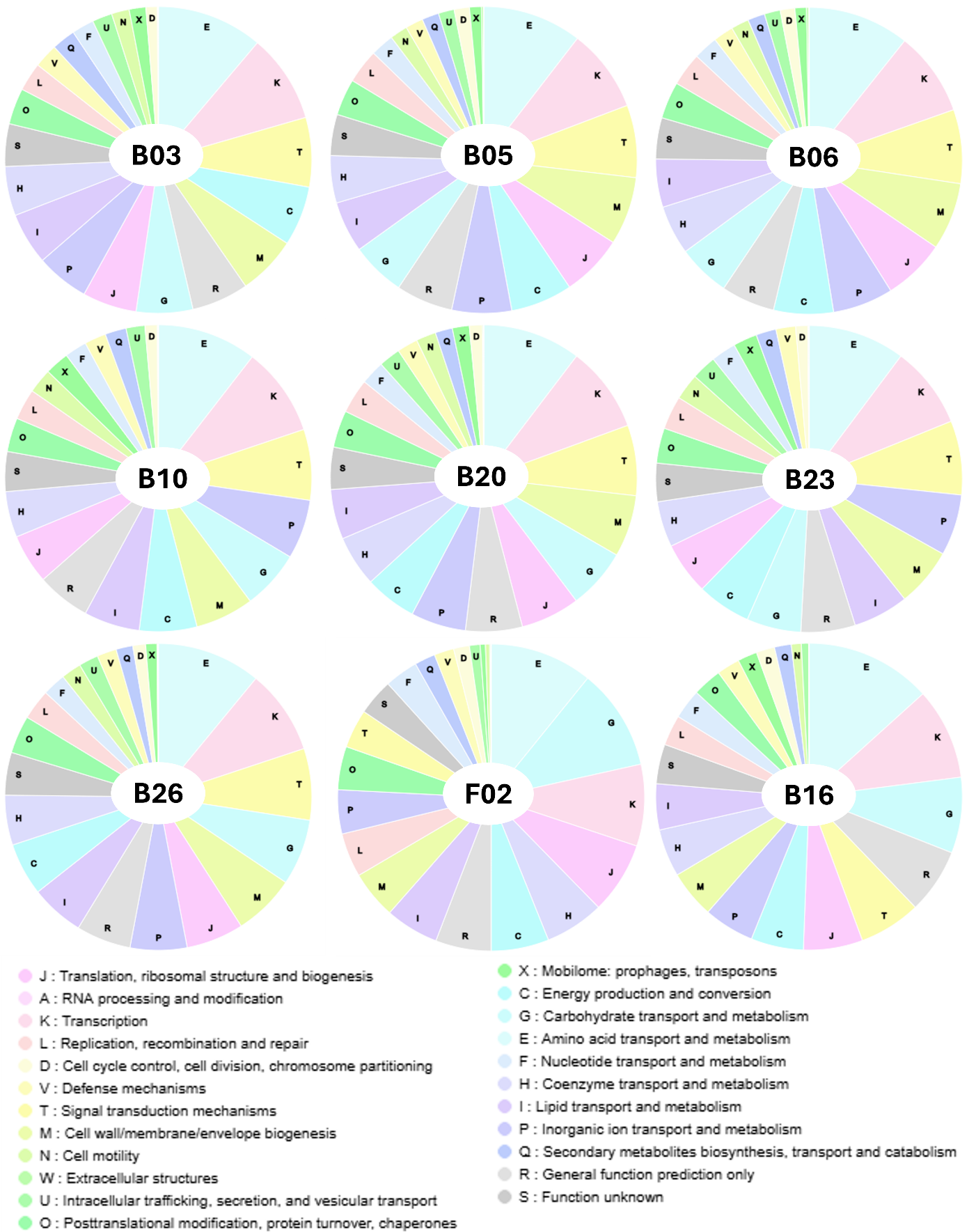

Supplementary Figure S8. COG Functional Classification of Pseudomonas, Arthrobacter, and Brevibacillus isolates. Pie charts represent the distribution of predicted proteins of Pseudomonas (B03, B05, B06, B10, B20, B23, B26), Arthrobacter (F02), and Brevibacillus (B16) isolates across different Clusters of Orthologous Groups (COGs) functional categories, based on the number of sequences grouping in each category.

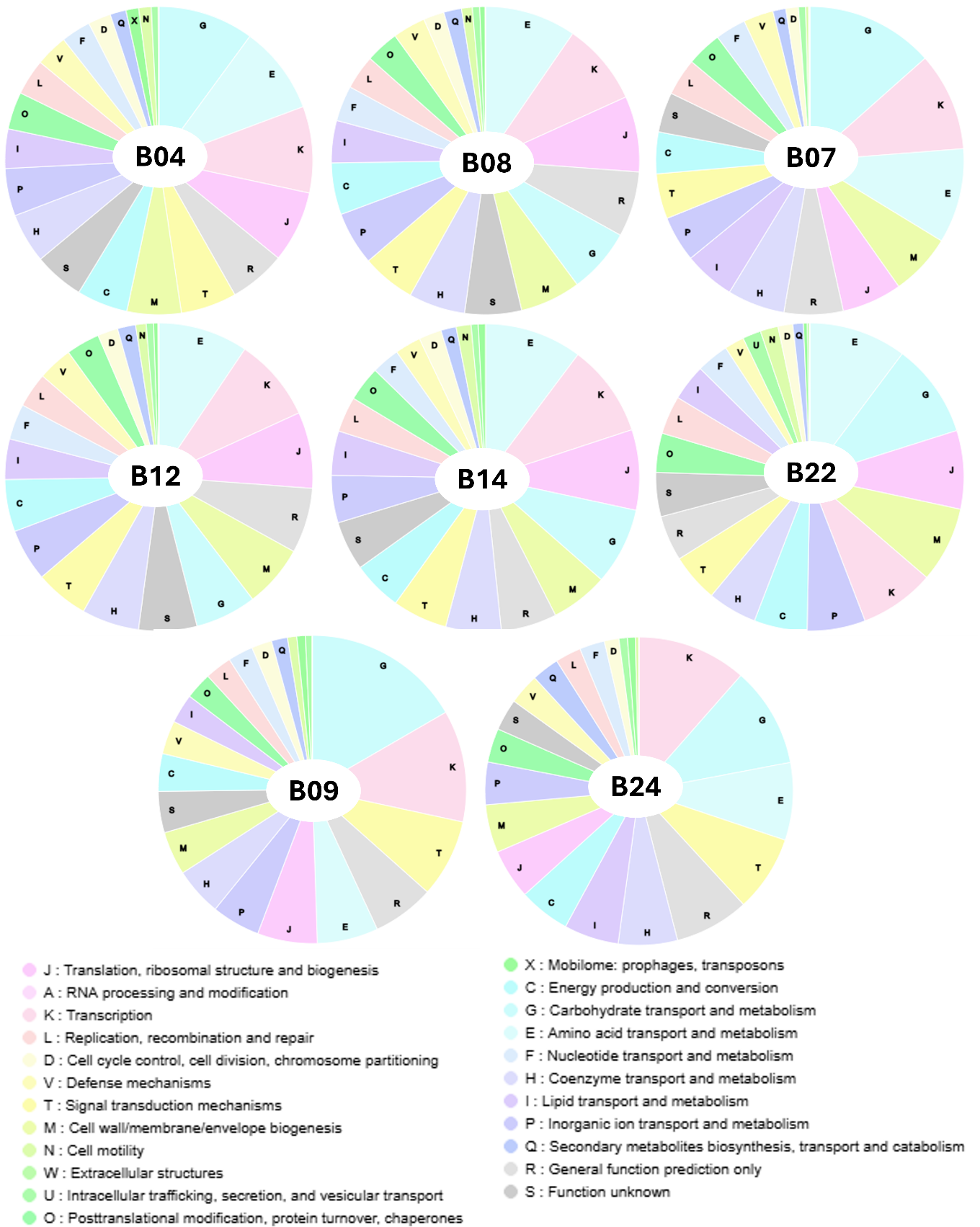

Supplementary Figure S9. COG functional classification of Bacillus, Plantibacter, Pantoea, Paenibacillus, and Streptomyces isolates. Pie charts represent the distribution of predicted proteins of Bacillus (B04, B08, B12, B14), Plantibacter (B07), Pantoea (B22), Paenibacillus (B09), and Streptomyces (B24) isolates across different Clusters of Orthologous Groups (COGs) functional categories, based on the number of sequences grouping in each category.

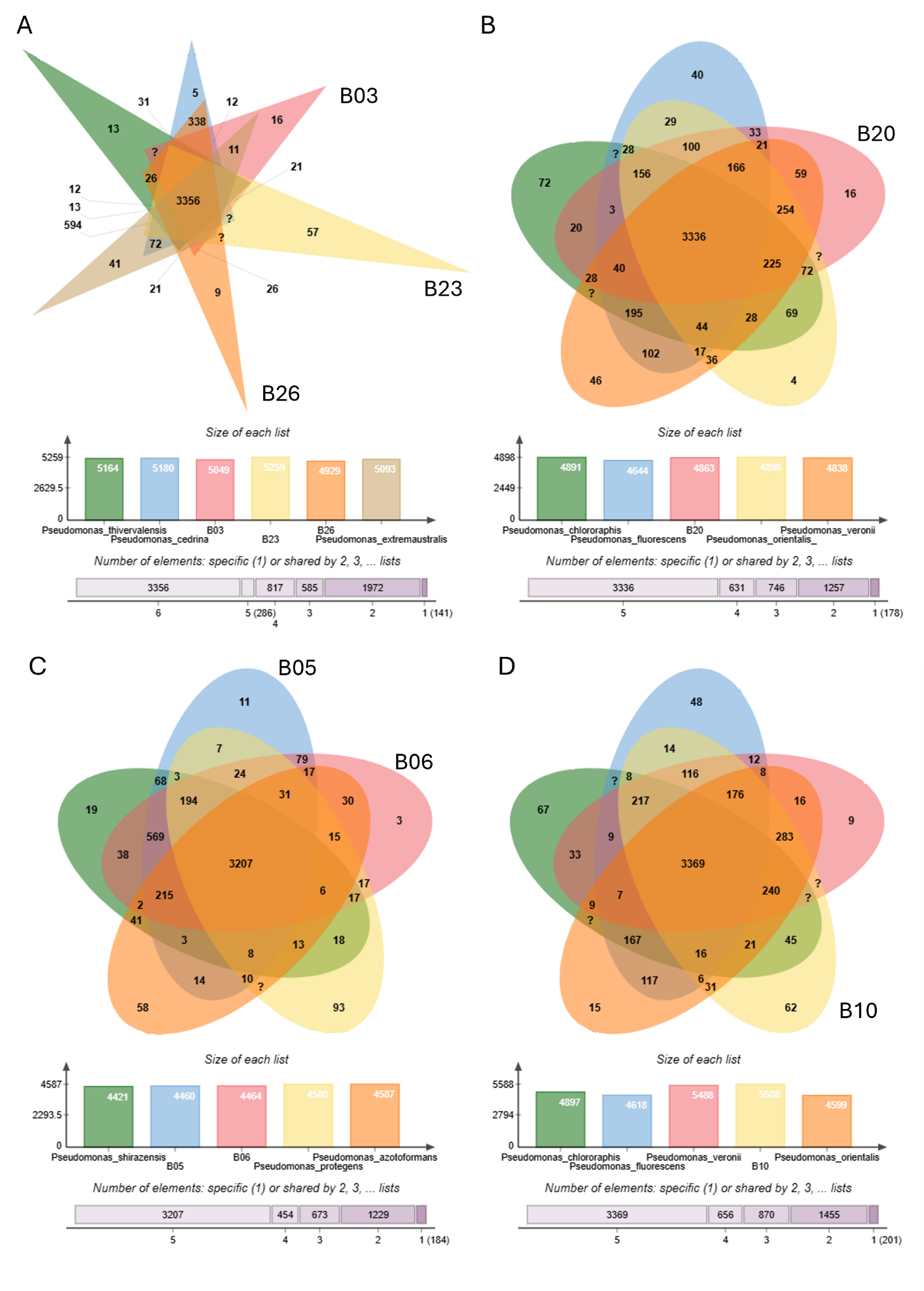

Supplementary Figure S10. Comparative genomics of Pseudomonas isolates and related species using OrthoVenn. OrthoVenn diagrams illustrating the distribution of orthologous gene clusters among (A) Isolates B03, B23, B26, Pseudomonas thivervalensis, P. cedrina, and P. extremaustralis; (B) B20, P. chloraphis, P. fluorescens, P. orientalis, and P. veronii; (C) Isolates B05, B06, P. shirazensis, P. protegens, and P. azotoformans; and (D) Isolate B10, P. chlororaphis, P. fluorescens, P. veronii, and P. orientalis. The intersecting areas represent the number of shared orthologous gene clusters, while non-overlapping areas indicate unique clusters for each species/isolate. Bar plots below each Venn diagram show the total number of orthologous gene clusters for each species/isolate and the number of specific or shared clusters among different numbers of lists. "?" indicates values exceeding the display limit.

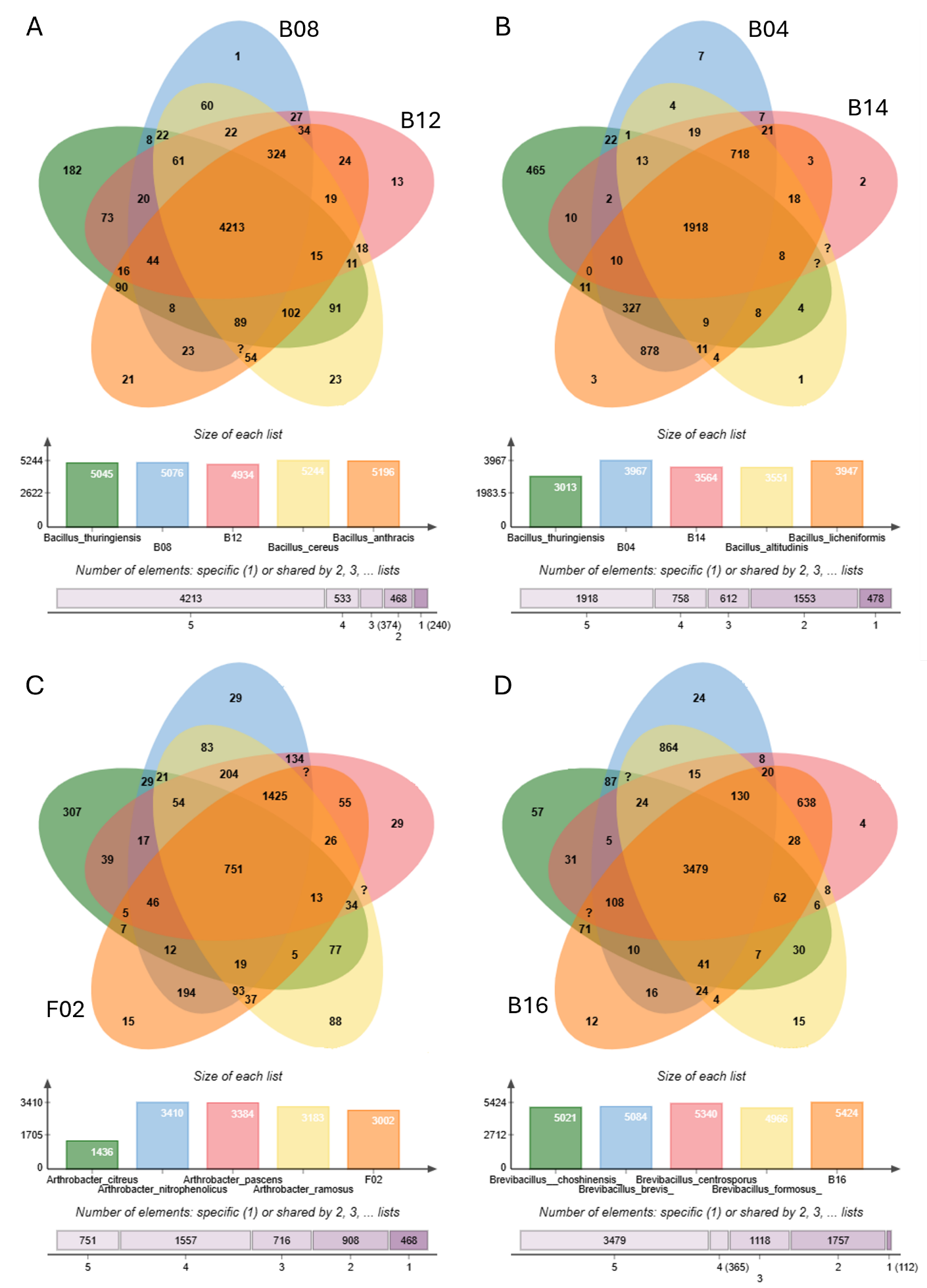

Supplementary Figure S11. Distribution of Orthologous Gene Clusters in Bacillus, Arthobacter, and Brevibacillus Isolates and Related Species. OrthoVenn diagrams illustrate the distribution of orthologous gene clusters among specific bacterial isolates and their closely related species: (A) Isolates B08 and B12, along with Bacillus thuringiensis, B. cereus, and B. anthracis; (B) Isolates B04 and B14, along with B. thuringiensis, B. altitudinis, and B. licheniformis; (C) Isolate F02, along with Arthrobacter citreus, A. nitrophenolicus, A. pascens, and A. ramosus; and (D) Isolate B16, along with Brevibacillus choshinensis, B. brevis, B. centrosporus, and B. fomosus. Intersecting areas within each diagram represent the number of shared orthologous gene clusters, while non-overlapping areas indicate unique clusters for each isolate or species. Bar plots below each Venn diagram show the total number of orthologous gene clusters per isolate/species and the number of specific or shared clusters among different numbers of lists. The symbol "?" indicates values exceeding the display limit of the diagram.

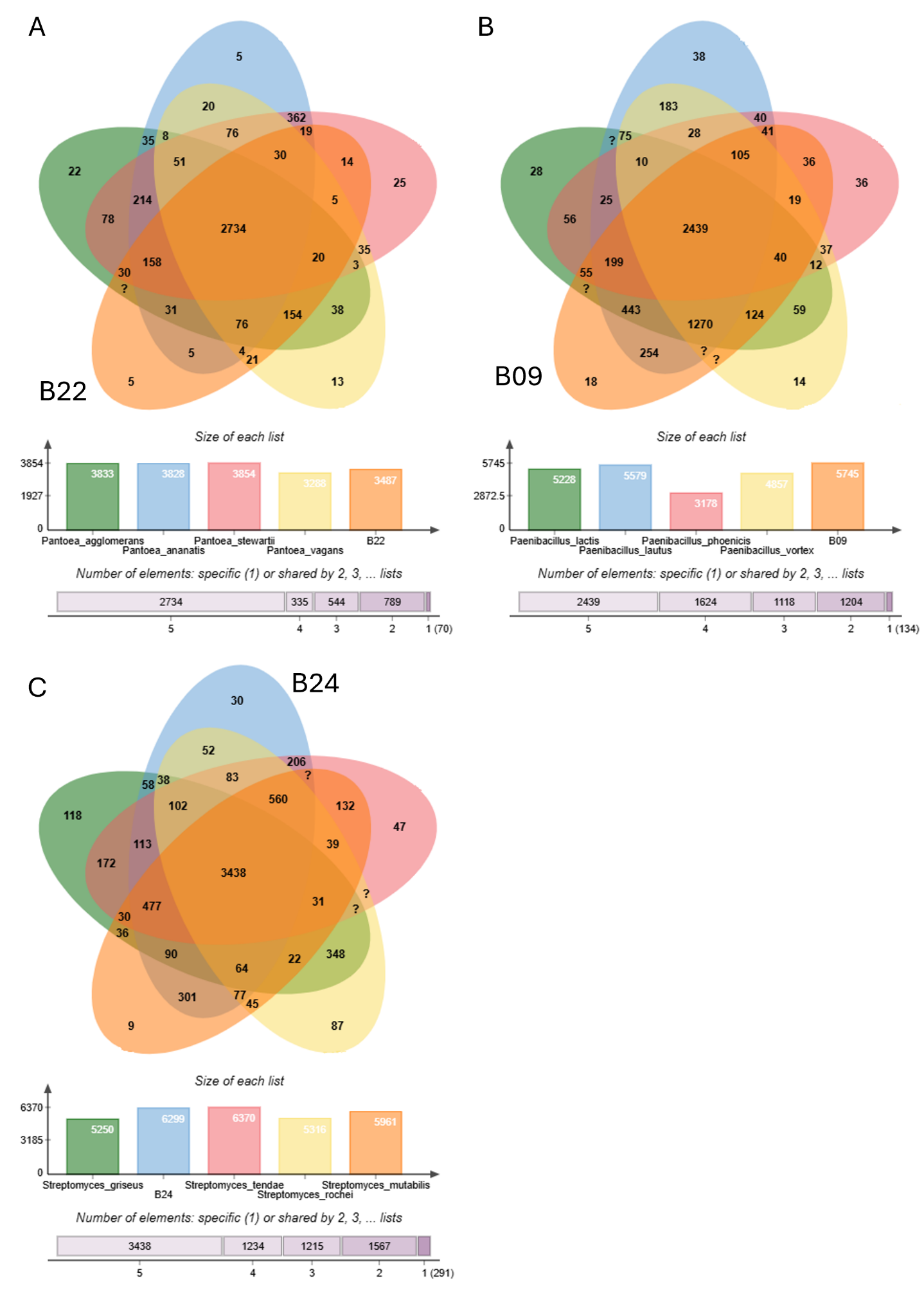

Supplementary Figure S12. Orthologous Gene Cluster Distribution in Pantoea, Paenibacillus, and Streptomyces Isolates and Related Species. OrthoVenn diagrams illustrate the distribution of orthologous gene clusters among: (A) Isolate B22, along with Pantoea agglomerans, Pa. ananatis, Pa. stewartii, and Pa. vagans; (B) Isolate B09, along with Paenibacillus lactis, P. lautus, P. phoenicis, and P. vortex; and (C) Isolate B24, along with Streptomyces griseus, S. tendae, S. rochei, and S. mutabilis. Intersecting areas within each diagram represent the number of shared orthologous gene clusters, while non-overlapping areas indicate unique clusters for each specific isolate or species. Bar plots below each Venn diagram quantify the total number of orthologous gene clusters per isolate/species and the number of specific or shared clusters among different numbers of groups. The symbol "?" indicates values exceeding the display limit of the diagram.

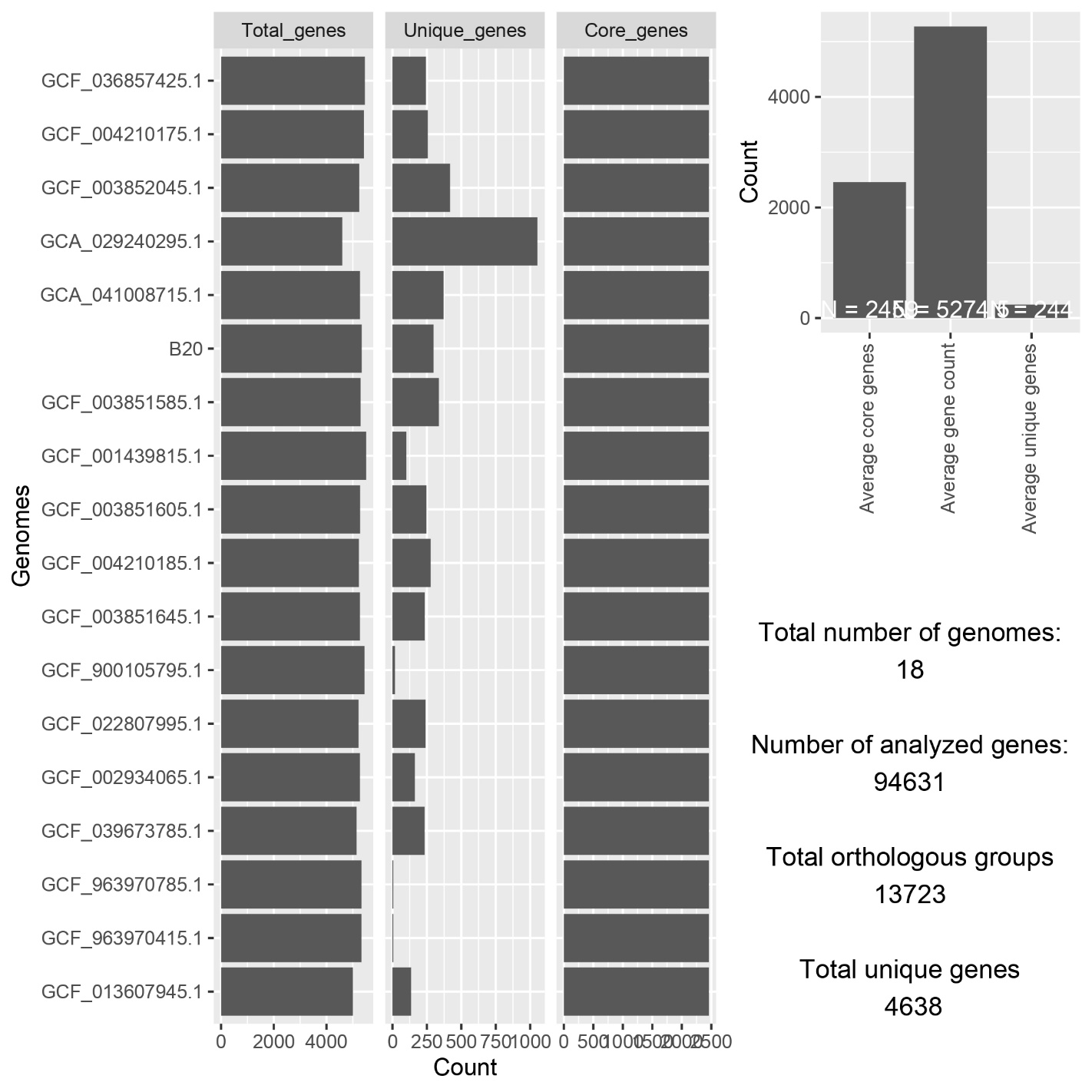

Supplementary Figure S13. Summary statistics for the pan-genome analysis of P. orientalis, providing a comprehensive overview of the gene diversity across 18 genomes. It includes metrics such as the total gene count, core genes, and unique genes. Key statistics reveal that 94,631 genes were analyzed, organized into 13,723 orthologous groups, with 4,638 genes classified as unique. The bar plots illustrate the distribution of total, unique, and core genes across genomes, highlighting variability in genome content. These statistics offer valuable insights into gene conservation, diversity, and the evolutionary dynamics of this group, which are crucial for understanding genomic architecture and adaptation.

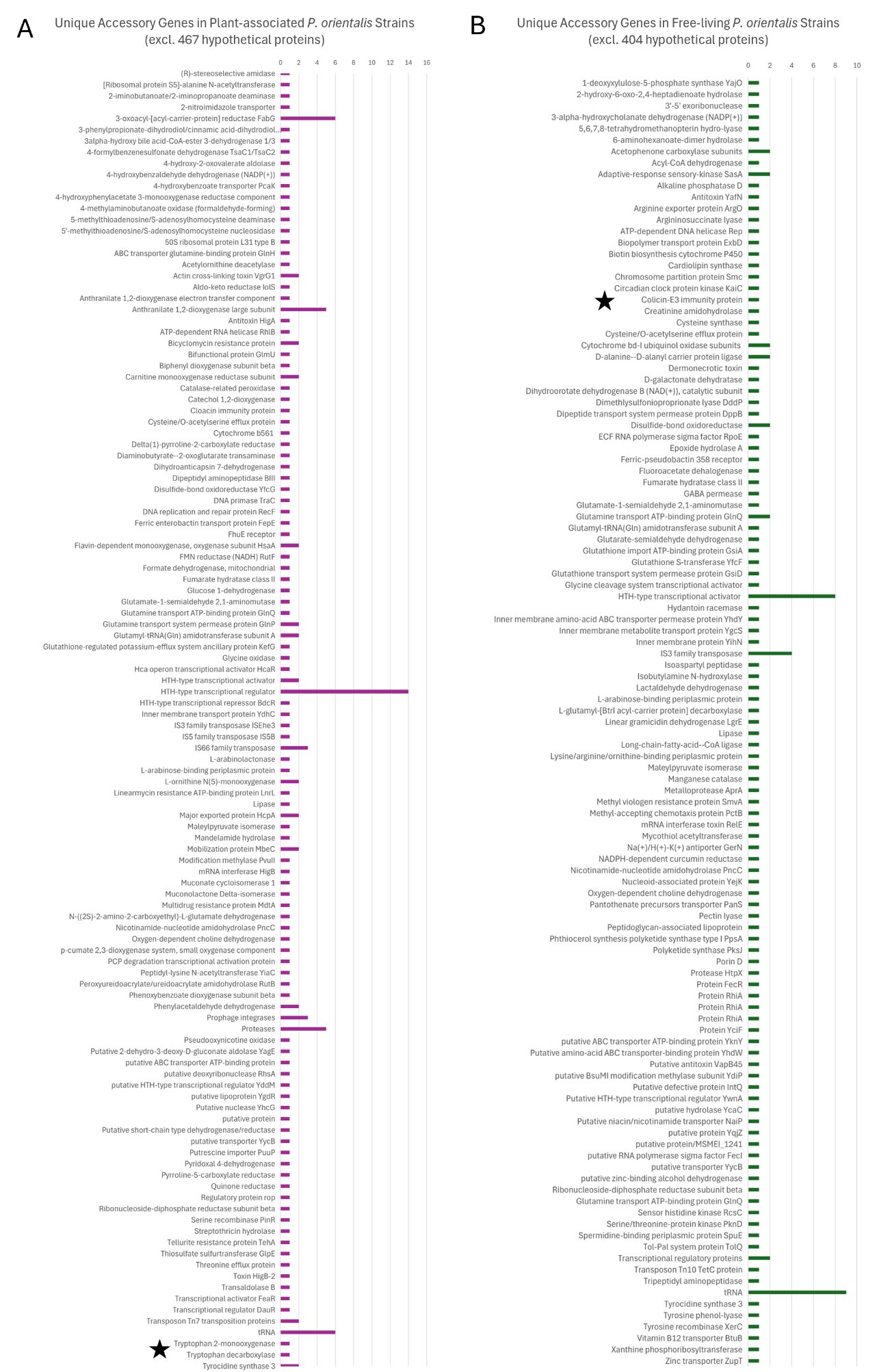

Supplementary Figure S14. Unique Accessory Genes in Plant-associated (A) and Free-living (B) Pseudomonas orientalis strains. Stars indicate the interesting genes discussed.
